# Investigating equations for measuring dissolved inorganic nutrient uptake in oligotrophic conditions

**DOI:** 10.1101/2020.08.30.274449

**Authors:** Michael R. Stukel

**Author notes:** Correspondence to Michael Stukel.

## Abstract

Multiple different equations have been used to quantify nutrient uptake rates from stable isotope tracer label incorporation experiments. Each of these equations implicitly assumes an underlying model for phytoplankton nutrient uptake behavior within the incubation bottle and/or pelagic environment. However, the applicability of different equations remains in question and uncertainty arising from subjective choices of which equation to use is never reported. In this study, I use two approaches to investigate the conditions under which different nutrient uptake equations should be used. First, I utilized a moderate-complexity pelagic ecosystem model that explicitly models the δ^15^N values of all model compartments (NEMURO+^15^N) to conduct simulated nitrate uptake and ammonium uptake incubations and quantify the accuracy of different nutrient uptake equations. Second, I used results of deckboard diel nutrient uptake experiments to quantify the biases of 24-h incubations relative to six consecutive 4-h incubations. Using both approaches, I found that equations that account for nutrient regeneration (i.e., isotope dilution) are more accurate than equations that do not, particularly when nutrient concentrations are low but uptake is relatively high. Furthermore, I find that if the goal is to estimate *in situ* uptake rates it is appropriate to use an *in situ* correction to standard equations. I also present complete equations for quantifying uncertainty in nutrient uptake experiments using each nutrient uptake equation and make all of these calculations available as Excel spreadsheets and Matlab scripts.

## INTRODUCTION

Numerous studies quantifying nitrate, ammonium, and phosphate uptake kinetics in the pelagic are motivated by the fact that low nutrient concentrations often limit phytoplankton growth in the vast oligotrophic regions of the ocean (Dugdale 1967; Mulholland and Lomas 2008). Such studies most frequently use uptake of stable isotope labeled nutrient tracers (e.g., ^15^NO_3_^-^ or ^15^NH_4_^+^) (Dugdale and Wilkerson 1986; Glibert et al. 2018). In such studies, the accumulation of heavy isotope label into particulate organic matter is quantified and uptake rates are typically determined using some version of the following equation:

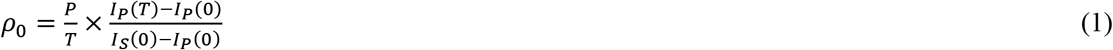

In this equation (originally from Dugdale and Goering 1967), ρ_0_ is the computed nutrient uptake rate (typically units of μmol L^-1^ h^-1^), P is particulate organic nitrogen (PON) concentration, T is the duration of the incubation, I_p_(T) is the isotope ratio of the particulate nitrogen at the end of the incubation, I_S_(0) is the isotope ratio of the substrate pool at the beginning of the experiment, and I_P_(0) is the natural isotope ratio in the particulate pool at the beginning of the experiment (which most frequently is not measured and instead is assumed to be equal to the average isotopic ratio of phytoplankton). Most frequently, P is measured at the end of the experiment and I_S_(0) is calculated as:

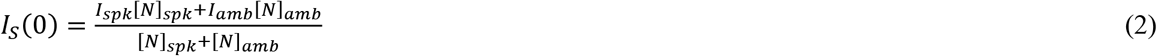

where I_spk_ and [N]_spk_ are the isotope ratio and concentration of the ^15^N-labeled nutrient spike and I_amb_ and [N]_amb_ are the natural isotope ratio of the nutrient pool and the ambient nutrient concentration, respectively (Table 1). However, modifications to this equation are sometimes made based on whether P is determined at the end of the incubation, beginning, or both, and whether or not the isotopic ratio of natural POM is determined (see Dugdale and Wilkerson 1986 for more details). Collos (1987) suggested that when more than one nutrient pool is available (e.g., NO_3_^-^ and NH_4_^+^) only versions of Eq. 1 using the final PON concentration for P are accurate, thus I will use this form for future calculations of p_0_.

**Table 1.** Parameter Definitions

| <u>Variable</u> | <u>Definition</u> | <u>Units</u> |
| --- | --- | --- |
| $\rho_0$ | Calculated incubation nutrient uptake (Eq. 1) | $\text{nmol N L}^{-1} \text{ h}^{-1}$ |
| $\rho_{\text{kan}}$ | Calculated incubation nutrient uptake (Eq. 3) | $\text{nmol N L}^{-1} \text{ h}^{-1}$ |
| $\rho_{\text{reg}}$ | Calculated incubation nutrient uptake (Eq. 13) | $\text{nmol N L}^{-1} \text{ h}^{-1}$ |
| $\rho_{0,\text{is}}$ | Calculated incubation nutrient uptake (Eq. 5) | $\text{nmol N L}^{-1} \text{ h}^{-1}$ |
| $\rho_{\text{kan},\text{is}}$ | Calculated incubation nutrient uptake (Eq. 6) | $\text{nmol N L}^{-1} \text{ h}^{-1}$ |
| $\rho_{\text{regis}}$ | Calculated incubation nutrient uptake (Eq. 14) | $\text{nmol N L}^{-1} \text{ h}^{-1}$ |
| T | Duration of Incubation | hours |
| P | PON (final) | $\mu\text{mol N L}^{-1}$ |
| $I_P(T)$ | Isotope Ratio of PON (final) | atom fraction |
| $I_P(0)$ | Isotope Ratio of PON (initial) | atom fraction |
| $I_{\text{spk}}$ | Isotope Ratio of tracer spike | atom fraction |
| $I_{\text{amb}}$ | Isotope Ratio of ambient nutrient | atom fraction |
| $[N]_{\text{spk}}$ | Final concentration of tracer | $\mu\text{mol N L}^{-1}$ |
| $[N]_{\text{amb}}$ | Ambient Nutrient Concentration | $\mu\text{mol N L}^{-1}$ |
| r | Rate of nutrient regeneration | $\text{nmol N L}^{-1} \text{ h}^{-1}$ |
| a | Ratio of nutrient regeneration to uptake ($=r/\rho$) | unitless |
| b | $\rho_0 \times T / ([N]_{\text{amb}} + [N]_{\text{spk}})$ | unitless |

While Eq. 1 is widely used, it assumes that the isotopic ratio of the labeled nutrient pool remains constant throughout the duration of the experiment. If the nutrient is regenerated during the experiment, the heavy isotope spike will be diluted, lowering the isotopic ratio of the substrate during the experiment. This will lead Eq. 1 to underestimate *in situ* nutrient uptake. Blackburn et al. (1979), Caperon et al. (1979), and Glibert et al. (1982) developed equations for simultaneously quantifying nutrient uptake and regeneration by monitoring substrate concentration and isotope ratio. These measurements, however, are quite technically challenging when substrate concentrations are low and hence not realistic for most studies that quantify nutrient uptake in oligotrophic regions. To address these issues, Kanda et al. (1987) made the assumptions that nutrient regeneration and uptake are constant throughout the duration of an incubation, that regenerated nutrients have an isotopic ratio equal to I_0_, and that ∂I_p_(t)/∂t (the rate of change of isotope ratio in PON) was constant in time. Given these assumptions, Kanda et al. (1987) estimated nutrient uptake as:

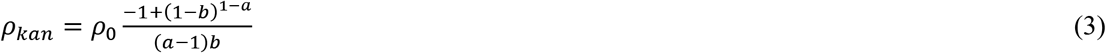

where a is equal to the ratio of nutrient regeneration rate to nutrient uptake rate (r/ρ) and:

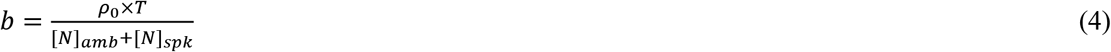

Kanda et al. (1987) originally formulated Eq. 3 for use with NH_4_^+^ uptake measurements, since NH_4_^+^ is rapidly regenerated by biological processes in the surface ocean. Recent evidence documenting shallow water nitrification in many ecosystems (Yool et al. 2007) raises the question of whether Eq. 3 is appropriate for NO_3_^-^ uptake experiments as well. Furthermore, two key assumptions add substantial uncertainty to results derived from Eq. 3. As noted before, Eq. 3 assumes that regenerated nutrients are isotopically equal to the isotopic ratio of the ambient nutrient pool and that ∂I_p_(t)/∂t is constant. Both of these assumptions are problematic when nutrient recycling within the incubation is substantial (i.e. when ρ×T>>[N_spk_+N_amb_]). In the limit as ρ×T/[N_spk_+N_amb_] approaches infinity, I_S_(T) and I_P_(T) will both asymptote to (I_P_(0)×P(0)+Is(0)×S(0))/(P(0)+S(0)), while ∂I_p_(t)/∂t will approach zero at the end of the incubation. Both of these violate the assumptions of Eq. 3, leading to potentially problematic results. An additional complication for using Eq. 3 is that, the ratio of nutrient regeneration to nutrient uptake (r/ρ) must typically be estimated *a priori*, and hence potentially introduces substantial uncertainty.

Another assumption inherent to using Eqs. 1 and 3 to estimate in situ uptake rates is the assumption that nutrient uptake rates in experimental bottle are representative of *in situ* uptake rates. Although bottle effects are potentially substantial and may lead to over- or underestimation of true rates (Laws 2013; Nogueira et al. 2014; Veldhuis and Timmermans 2007), I do not focus on them here. Instead, I consider the possibility that nutrient uptake rates are modified due to the addition of labeled nutrients. General practice considers this potential issue to be negligible if [N]_spk_ is <10% of [N]_amb_ (Dugdale 1967; Dugdale and Wilkerson 1986; McCarthy 1980). While spiking with such low nutrient concentrations may be good practice, it is not always possible if nutrient concentrations are not measured immediately at sea, which can be particularly difficult in oligotrophic areas where low-level nutrient methods may be necessary (e.g., Dore et al. 1996). Several studies have suggested computing a modified *in situ* nutrient uptake rate assuming that phytoplankton uptake follows a Monod-type saturating function, particularly if the tracer spike is >10% of ambient concentrations (Dugdale and Wilkerson 1986; Kanda et al. 2003; Rees et al. 1999). Such a modification can be calculated as:

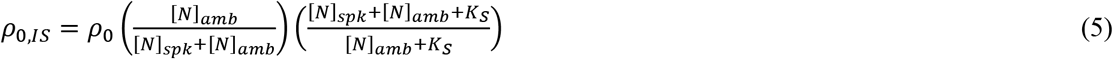

and:

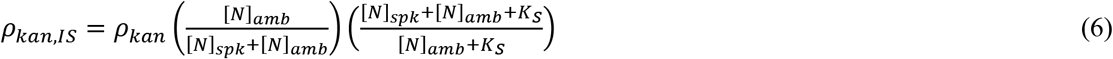

where K_S_ (half-saturation constant for phytoplankton nutrient uptake) is either measured *in situ* by conducting nutrient uptake measurements at multiple nutrient concentrations or assumed *a priori.*

In this study, I develop an unbiased equation (i.e. an equation that should not have systematic error) for estimating nutrient uptake when nutrient regeneration within the incubation bottle is significant. I then investigate the tradeoffs between calculating nutrient uptake using Eqs. 1, 3, 5, and 6 and this new equation in three ways: First, I present equations for measurement uncertainty determined by propagating measurement and parameter uncertainty through each equation (all equations are included in easy to use spreadsheets as supplementary material for this manuscript). I then use a plankton ecosystem model with a ^15^N isotope module to simulate phytoplankton nutrient uptake experiments and compare phytoplankton nutrient uptake calculated with each equation to “true” model nutrient uptake (Appendix 1). I use this approach because “truth” (i.e., actual environmental rates) can never be known for environment samples. However, in a simulated deterministic environment (i.e., the ecosystem model used here) the “true” rates within the simulated ecosystem are known and hence can be compared to the results of simulated incubations conducted in different conditions and using different nutrient uptake equations. My next approach is to use results from diel uptake experiments conducted in two open ocean regions to highlight the differences between these equations (Appendix 2). Finally, in a “recommendations” section, I discuss the situations during which each equation may be most appropriate to use.

## MATERIALS AND PROCEDURES

### Derivation of equation for nutrient uptake with recycled nutrients

If we make the assumption that nutrient uptake rates are constant throughout an incubation (a necessary simplifying assumption also made in Eqs. 1 and 3, although it is unlikely to be strictly true in real experiments), we can write two differential equations governing the concentrations of substrate (S) and particulate organic matter (P) in the incubation:

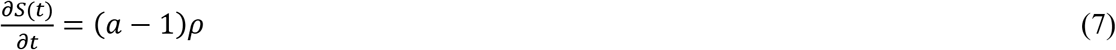

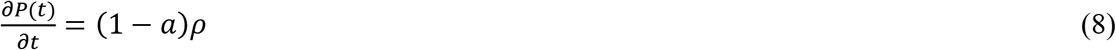

where a is the fraction of nutrient uptake that gets regenerated (as in Eq. 3). The amount of ^15^N (or other labeled substrate) taken up by phytoplankton can be easily estimated as ρ×I_S_(t), where I_S_(t) is the isotope ratio of the substrate (i.e. nutrient pool). The amount of ^15^N regenerated from P is potentially complex because phytoplankton are not the only form of particulate organic matter. Indeed, non-living detritus (which will not take up the labeled substrate) often comprises most of the POM (Riley 1970; Stukel et al. 2014; Yanada and Maita 1995). Thus the actual amount of ^15^N regenerated may depend on complex interactions within different components of particulate organic matter (phytoplankton, zooplankton, bacteria, archaebacteria, and detritus). However, if we make the null assumption that P is a well-mixed reservoir, we can write:

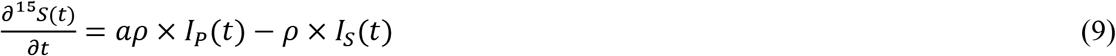

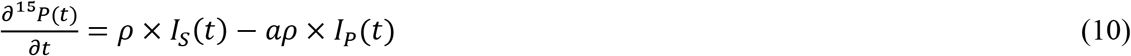

where ^15^S(t) is the total ^15^N in the substrate (nutrient pool) and ^15^P(t) is the total ^15^N in the particulate pool. The rate of change of the isotopic ratios of S and P can then be written as:

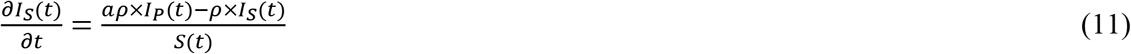

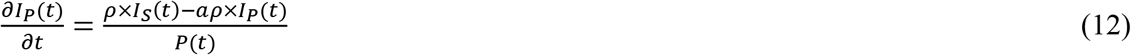

It is not possible to determine a closed-form solution for ρ as a function of only measurements typically made in incubation experiments. However, when ΔS is small relative to S and ΔP is small relative to P (both of which will be true when a≈1), ρ can be approximated by the equation:

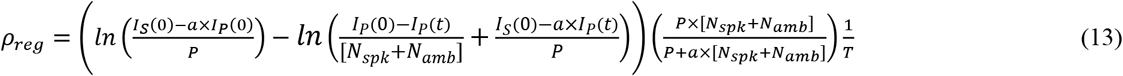

For a full derivation, see Online Supplementary Appendix S1. Following discussions above, I also define an equation for calculating *in situ* nutrient uptake when nutrient regeneration is substantial:

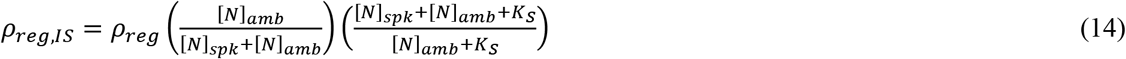

Importantly, for situations with substantial nutrient regeneration, Eq. 13 behaves in a sensible manner in the limit as t→∞. Specifically, with the system of differential equations outlined in Eqs. 7-12, I_P_(t) will approach:

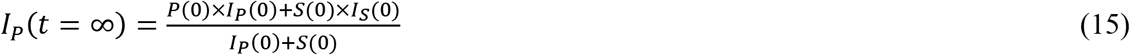

as the isotope label becomes evenly distributed between the substrate and particulate pool. When Ip(T) approaches the value in Eq. 15, the term inside the second natural logarithm in Eq. 13, will approach zero. Thus the term inside the first parentheses approaches infinity while 1/T approaches zero. This means that if the incubation experiment is run too long such that the isotope label becomes evenly distributed throughout the substrate and particulate pools, Eq. 13 becomes undefined reflecting an inability to constrain nutrient uptake rates under these conditions. Uncertainty in nutrient uptake calculated by ρ_reg_ also increases appropriately as I_P_(t) approaches its value in Eq. 15 (see Online Supplementary Appendix S2.5 for derivation of uncertainty). This is not the case for ρ0 or ρ_kan_. Both of these equations have I_S_(0) – I_P_(0) in the denominator and this term will never approach zero. Thus, because they also have time in the denominator, σ_ρ0_ and σ_ρkan_ (measurement uncertainty for ρ_0_ or ρ_kan_, respectively) both approach zero as time approaches infinity (see Online Supplementary Appendices S2.1 and S2.2 for derivation of uncertainty). This is clearly problematic and an argument in favor of using ρ_reg_ (Eq. 13) when recycling is expected to be substantial in the incubation bottle (Fig. 1).

**Fig. 1.**
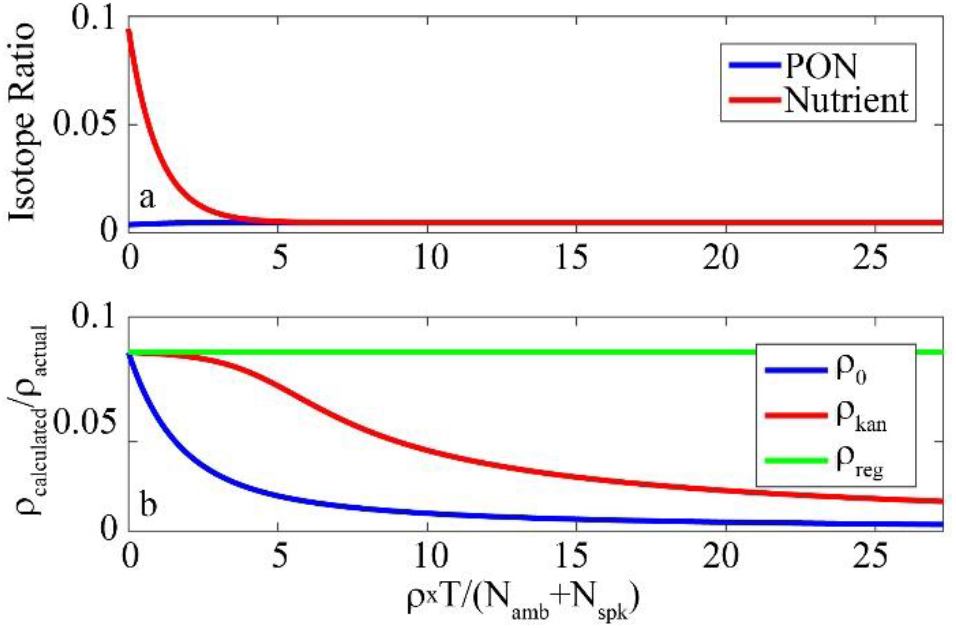
Comparison of nutrient uptake equations with complete nutrient regeneration (a = 1). Results based on numerical solution of system of differential equations described by Eqs. 7 – 12. Initial conditions were PON = 1 μmol L^-1^, N_amb_ = 0.01 μmol L^-1^, N_spk_ = 0.001 μmol L^-1^. Actual nutrient uptake rate was 100 nmol L^-1^ d^-1^. a) Isotope ratio in particulate and nutrient pools over time. b) Ratio of calculated nutrient uptake rate (using ρ_0_, ρ_kan_, or ρ_reg_) to actual nutrient uptake rate. Both are plotted against the number of times the nutrient pool is recycled within the incubation.

**Fig. 2.**
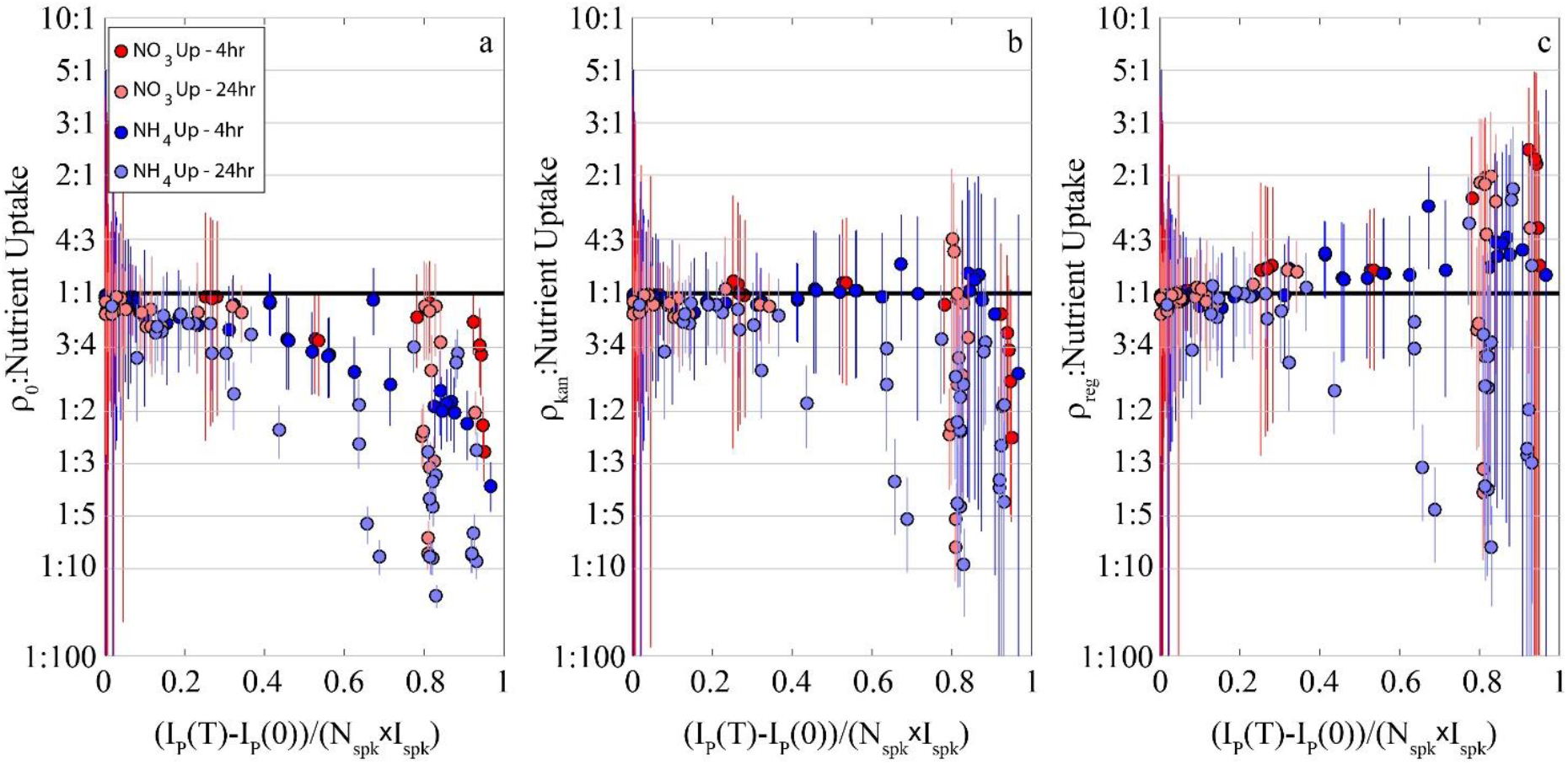
Comparison of nitrate uptake results calculated with ρ_0_, ρ_kan_, and ρ_reg_ to the fraction of labeled isotope present in the particulate pool at the end of the incubation (x-axis = (I_P_(T)-I_P_(0))/(N_spk_×I_spk_)). Results are from NEMURO+^15^N models run to steady-state and “sampled” for 4-h nitrate uptake incubations (dark red), 24-h nitrate uptake incubations (light red), 4-h ammonium uptake incubations (dark blue), and 24-h ammonium uptake incubations (light blue). These are the same experiments plotted in Fig. 6 and 7.

### Derivation of nutrient uptake uncertainty equations

Uncertainty in nutrient uptake experiments calculated from Eqs. 1, 3, 5, 6, 13, and 14, arises from uncertainty in T, P, I_P_(T), I_P_(0), I_spk_, I_amb_, N_spk_, N_amb_, a, and K_S_. However, I have not seen equations that calculate uncertainty in nutrient uptake by propagating uncertainty in each of these parameters through these equations. For each equation, I assume that measurement uncertainty in one parameter (e.g., P) is uncorrelated with measurement uncertainty in another parameter (e.g., I_spk_). I hence calculate propagation of uncertainty following the general chain rule:

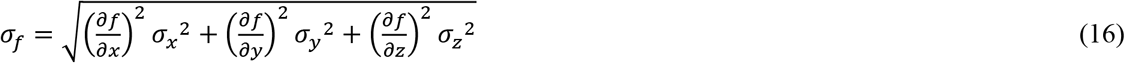

if *f* is a function of *x, y*, and *z.* Because of the complexity of Eqs. 1, 3, 5, 6, 13, and 14, the equations for their uncertainty cannot be displayed here. However, they are available in online Supplementary Appendix S2 in Eqs. B1, B13, B40, B58, B73, and B92, respectively. In the supplementary information I have also included Microsoft Excel spreadsheets and Mathworks Matlab scripts that can be used to calculate ρ and its associated uncertainty for each equation. These files are also available on GitHub at: https://github.com/stukel-lab/NUpCalculations.

Most frequently, studies report uncertainty in nutrient uptake rates as the standard deviation (or standard error) of duplicate or triplicate measurements. This is not, however, an appropriate estimate of true measurement uncertainty, because it does not account for correlated errors across the measurements. For instance with ρ_kan_ and ρ_reg_ the ratio of nutrient regeneration to nutrient uptake (a) must be assumed *a priori*. This *a priori* choice is associated with substantial uncertainty that is not reflected in uncertainty estimates made from triplicate incubations that all assume the same value of a. In online Supplementary Appendix S2.7 I explain how to calculate measurement uncertainty in the case of duplicate or triplicate measurements.

Eq. 16 has an implicit assumption that either higher order derivatives are negligible or the uncertainty in input parameters is small. When this is not the case, asymmetric confidence intervals are more appropriate. To compute asymmetric confidence intervals I used a standard Monte Carlo approach (Anderson 1976). Input parameters were randomly varied based on their assumed or measured uncertainty ranges and ρ_0_, ρ_kan_, ρ_reg_, ρ_0,is_, ρ_kan,is_, or ρ_reg,is_ were calculated based on these randomly distributed input parameters. A total of 1000 iterations were performed. Matlab code for calculating asymmetric confidence intervals for each of these equations is available in the supplemental material and on GitHub. In all plots I showed the symmetric confidence limits (Eq. 16) unless otherwise stated, because symmetric confidence limits are far more commonly used in the literature.

## RESULTS AND DISCUSSION

### Comparison of nutrient uptake equations

I also conducted diel experiments to assess each equation in field conditions by conducting paired 24-h incubations simultaneously with a series of six sequential 4-h incubations using bottles sampled from the same water (see Appendix 2 for full details). The assumption in these experiments is that 4-h incubations will be more accurate, but that with an unbiased equation (i.e. an equation with no systematic error) the results of the 24-h incubation will have higher uncertainty, but the confidence limits should still bracket the more accurate results determined by averaging rates determined from the six 4-h incubations. Results showed generally good agreement between 4-h and 24-h nitrate uptake equations with all equations, although ρ_0_ was more prone to underestimate nitrate uptake in the 24-h incubations (Fig. 3). All equations substantially underestimated ammonium uptake for the 24-h incubations, although the ρ_reg,is_ equation (when used with asymmetric confidence limits) did a much better job of accurately reflecting the uncertainty associated with these 24-h incubations.

**Fig. 3.**
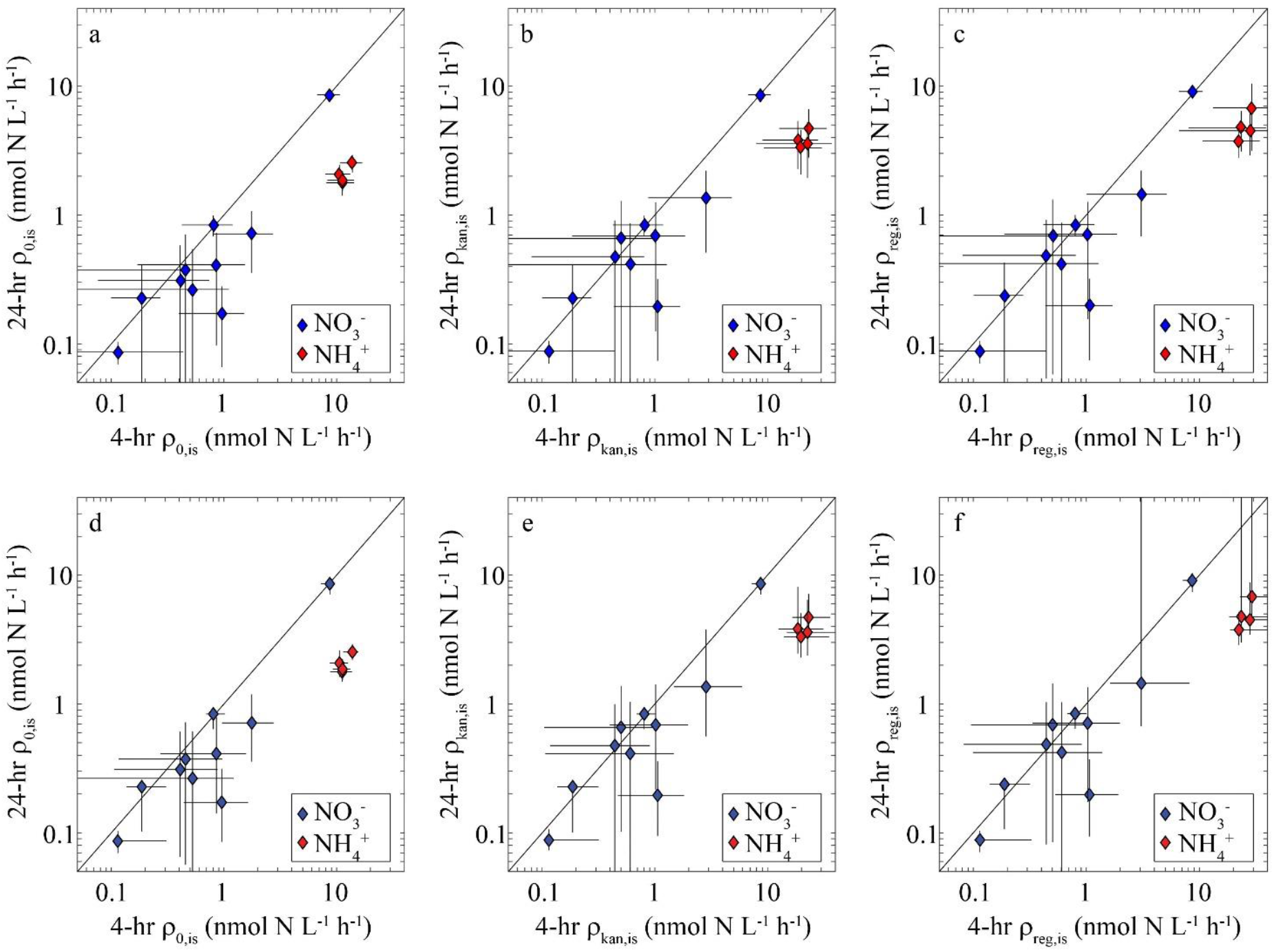
Comparison of 4-h (x-axis) and 24-h (y-axis) nutrient uptake rates conducted on cruises in the Gulf of Mexico and Costa Rica Dome computed using ρ_0,is_ (a,d), ρ_kan,is_ (b,e), and ρ_reg,is_ (c,f). Panels a-c show symmetric confidence intervals calculated using Eq. 16. Panels d-f show asymmetric confidence intervals calculated by Monte Carlo analyses.

### Uncertainty in nutrient uptake measurements

Uncertainty in nutrient uptake measurements can be derived from two distinct sources: parameter uncertainty and model uncertainty. By ‘parameter uncertainty’ I refer to uncertainty in parameters and measurements that are used in nutrient uptake calculations (e.g., K_S_, a, N_amb_, I_P_(T)). Uncertainty in these parameters can be approximated using symmetric confidence limits (Eq. 16) outlined in the online appendix and in the Matlab and excel files that I have included as a supplement to this manuscript. It can also be approximated more accurately using Monte Carlo approaches to quantify asymmetric confidence limits (also available in Matlab files in the supplement). Notably, I have not noted any studies that have accurately computed uncertainty arising from all parameters that are used as inputs to ρ_0_, ρ_kan_, ρ_reg_, ρ_0_,is, ρ_kan_,is, or ρ_reg,is_. Rather, most studies report uncertainty as the standard error of the mean (or standard deviation) of replicate measurements. Crucially, reporting uncertainty in this way only accounts for uncertainty in uncorrelated parameters (typically I_P_(T), P, and N_spk_), while it neglects uncertainty in correlated errors (most notably N_amb_ and, for some equations, a and K_S_). This thus understates the true uncertainty arising from the parameters in the equations. A more accurate approach is to combine the standard error of the mean with the uncertainty in these other parameters as outlined in online Supplementary Appendix S2.7.

Use of tracer label at a concentration < 10% of the ambient nutrient concentration, as recommended originally by Dugdale and Goering (1967), may be advantageous and serves two important purposes. It minimizes differences between *in situ* uptake and bottle uptake (i.e., between ρ_0_ and ρ_0,is_). It also decreases the likelihood that a perturbation will lead to nonlinear uptake rates. However, at very low nutrient concentrations, phytoplankton may be able to completely consume a 10% nutrient spike during the incubation. Hence, Glibert and Capone (1993) have recommended a target tracer concentration of 10% ambient particulate nitrogen.

### Recommendations

Ideally, uptake experiments would keep labeled nutrient tracer spikes < 10% of ambient nutrient and terminate incubations before substantial nutrient regeneration occurs. If such conditions are met, all 6 equations tested in this study give similar results. However, achieving both of these goals can be difficult at low nutrient concentrations, so it may be advisable to spike with tracer equal to ~10% of particulate nitrogen. An ideal experimental design would probably include triplicate 2- to 4-h nutrient uptake incubations conducted at natural light levels and repeated continuously over a full diel period to test for non-linearity. If possible, measurements of substrate isotope dilution are also advisable (Caperon et al. 1979; Glibert et al. 1982; Hansell and Goering 1989). However, for full euphotic zone coverage, the above would likely need to be replicated at 6 – 8 depths. Such experimental designs are certainly not practical in most situations. It is thus important to carefully consider (especially in oligotrophic conditions) which equation to use and how confidence limits should be determined.

First, it is important to consider the goal of the experiment (Fig. 4). When trying to quantify *in situ* nutrient uptake rates ρ_0,is_, ρ_kan,is_, or ρ_reg,is_ should be used. However, if the goal is to investigate nutrient uptake kinetics by manipulating nutrient concentrations, ρ_0_, ρ_kan_, or ρ_reg_ should be used. The next decision depends on whether the nutrient being measured is expected to be recycled in the incubation bottle. Typically, ammonium is recycled during the experiment, while nitrate may or may not be recycled. However, the equations in this experiment can also be used for other uptake experiments including urea uptake (recycling should be assumed), phosphate uptake (also recycled), or bicarbonate uptake (minimal recycling). When recycling is substantial, it is always appropriate to use ρ_kan_ or ρ_reg_ (or ρ_kan,is_ or ρ_reg,is_).

**Fig. 4.**
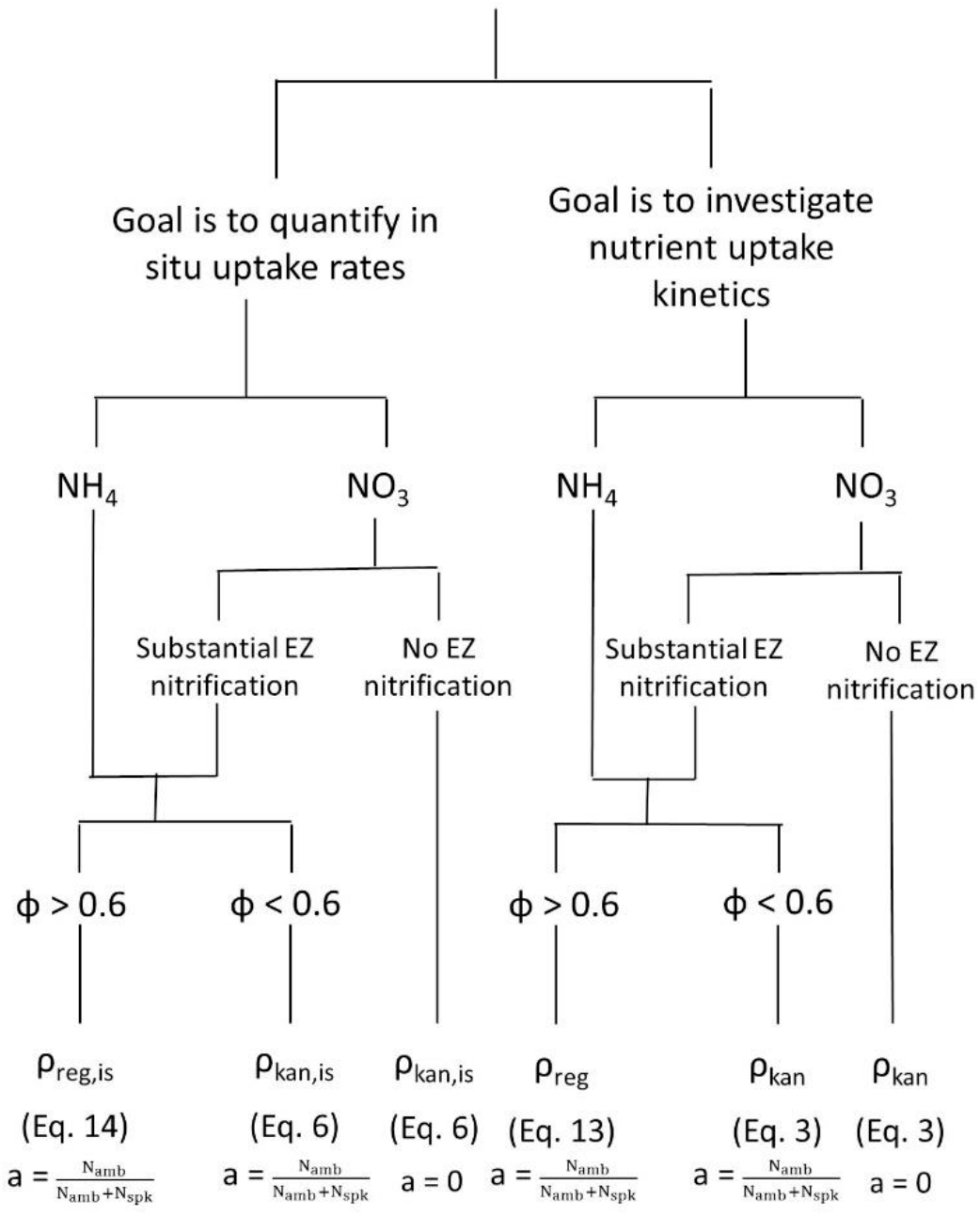
Recommended flow chart for deciding which nutrient uptake equation to use. Φ is the ratio of labeled nutrient incorporated in particulate matter at the end of the experiment and is equal to (I_P_(T)-I_P_(0))/(N_spk_×I_spk_). EZ = Euphotic zone.

It can be unclear whether or not experimentalists should assume that nitrate is being recycled within incubation bottles. While it was originally assumed that nitrate is only regenerated beneath the euphotic zone (Eppley and Peterson 1979), abundant recent evidence suggests that nitrification can occur in the euphotic zone (Yool et al. 2007). Earlier results apparently showing light limitation of nitrification may have resulted from competition between nitrifiers and phytoplankton for ammonium (Smith et al. 2014; Xu et al. 2019). Determining whether substantial nitrate recycling is occurring in the euphotic zone can be challenging. Ideally, independent nitrification rate measurements can be conducted by spiking separate bottles with labeled ammonium and quantifying ^15^NO_3_^-^ (e.g., Buchwald et al. 2015; Santoro et al. 2013). When nitrification rate measurements are not feasible, investigators can use nitrate profiles and reasonable estimates of vertical eddy diffusivity and/or upwelling to estimate whether or not diffusive and advective transport can supply enough nitrate to the euphotic zone to support measured nitrate uptake rates (Caffin et al. 2017; Fernández-Castro et al. 2015; Painter et al. 2013).

It is also important to choose a reasonable estimate for a (the ratio of nutrient recycling to nutrient uptake). If complete recycling is expected, I recommend that investigators assume a = N_amb_/(N_amb_+N_spk_), rather than 1. This assumes that nutrient regeneration rates remain at the levels that would occur prior to the perturbation caused by adding a nutrient spike. Regardless of what value is chosen for a, using appropriate values for σa should ensure that errors in this unknown parameter are properly incorporated.

When only a small fraction of the ^15^N-labeled spike is incorporated into particulate matter during the incubation, ρ_0_, ρ_kan_, and ρ_reg_ give similar results. In such situations, I recommend using ρ_kan_ (or ρ_kan,is_). ρk an notably simplifies to ρ_0_ (the most commonly used equation for nutrient uptake) when a = 0. However, uncertainty limits determined for ρ_kan_ appropriately incorporate uncertainty in the amount of recycling occurring within the incubation bottle. When a large fraction of the ^15^N-labeled spike is incorporated into particulate matter during the incubation, I recommend the use of ρ_reg_ (or ρ_reg,is_), because when Φ > 0.6 (where Φ = (I_P_(T)-I_P_(0))/(N_spk_×I_spk_)) ρ_kan_ has a significant negative bias, while ρ_reg_ is unbiased (Fig. 3). Furthermore, when asymmetric confidence limits are used, it is more likely that uncertainty limits from ρ_reg_ will bracket the true values.

I further recommend that investigators use the Excel or Matlab files included with this manuscript to investigate the sensitivity of their results to choices of which equation to use. Although the symmetric confidence limits are easier to use, investigators should also compute the asymmetric confidence limits to assess whether or not the symmetric confidence limits are appropriate for any particular incubation experiment. Furthermore, I believe that it is important that authors submit not only their estimates of nutrient uptake rates, but also the parameters that go into these estimates (e.g., P, T, I_P_(T), N_amb_, N_spk_) when they archive their data in data repositories and/or as supplements to manuscripts. This will ensure that future investigators can thoroughly evaluate the assumptions made by the original authors if they intend to use this data in the future or compare results from studies using different equations (e.g., Aufdenkampe et al. 2001). Nutrient uptake rate measurements are never exact and it is important that we allow other investigators to independently evaluate our assumptions and related confidence limits.

## Supporting information

Calculation Files for Nutrient Uptake Eqs

NEMURO+15N Files

## APPENDIX 1: Assessment using a simulated ecosystem framework

### Modeling approach

To model the planktonic ecosystem and changing isotopic ratios of ecosystem compartments, I use the NEMURO+^15^N model described in Stukel et al. (2018b). This model combines the NEMURO biogeochemical model (Kishi et al. 2011; Kishi et al. 2007) with the nitrogen isotope model of Yoshikawa et al. (2005). Briefly, NEMURO is a nitrogen currency planktonic ecosystem model with three nutrient pools (nitrate, ammonium, and silicic acid), two phytoplankton (large and small), three zooplankton (large, small, and predatory), dissolved organic nitrogen, detritus, and detrital silica. NEMURO was chosen because it is a well-known model that includes some of the key components necessary for testing the accuracy of nitrate uptake (i.e., two nitrogenous nutrient pools, multiple phytoplankton and zooplankton groups, nitrification). However, it is important to note that NEMURO does not include heterotrophic bacteria that may compete with phytoplankton for NH_4_^+^ or N2 fixation, which can be an important source of new nitrogen that can be converted into recycled DON or NH_4_^+^. These processes are potentially important in oligotrophic regions and may impact the results of nutrient uptake experiments (Dugdale and Goering 1967; Kirchman 1994). The sensitivity of NEMURO has also been extensively investigated (Ito et al. 2010; Rose et al. 2007; Yoshie et al. 2007) and its parameters have been tuned using *in situ* phytoplankton experiments (Li et al. 2011; Li et al. 2010). Stukel et al. (2018b) added a nitrogen isotopic model (Yoshikawa et al. 2005) to the NEMURO framework. This model adds state variables tracking the ^15^N content of each nitrogen-containing state variable and includes isotopic fractionation during nutrient uptake, excretion, egestion, nitrification, and remineralization. For a detailed description of the model, I refer readers to Kishi et al. (2007) and Stukel et al. (2018b).

One minor modification was made to NEMURO: the term for basal phytoplankton respiration was replaced with a basal excretion rate, because in the original configuration of NEMURO, when phytoplankton respiration exceeds phytoplankton growth rates, phytoplankton actually excrete nutrients (nitrate and ammonium) in the proportion to which they would take up those nutrients (if their growth rate exceeded respiration). This leads to unrealistic dynamics in which phytoplankton taxa can serve as a source of nitrate in the mixed layer under some conditions, particularly in oligotrophic regions.

To simulate conditions in the open ocean, I ran NEMURO+^15^N in a simple one-dimensional physical framework from the surface to a depth of 300-m with 2-m vertical resolution and 1-minute temporal resolution. I assumed a constant vertical diffusivity of a 10^-3^ m^2^ s^-1^ (default) within a 20-m mixed layer that gradually decreased to 10^-5^ m^2^ s^-1^ (default) at a depth of twice the mixed layer depth and remained constant with depth below that range. To create multiple different ecosystem configurations, I ran the model 37 times with different physical and biological parameterizations (Fig. 5; Supp. Table 1). I do not argue that any one parameter set is more or less accurate, but rather test the relative accuracy of Eqs. 1, 3, 5, 6. 13 and 14 under these scenarios that capture different phytoplankton and nutrient dynamics.

**Fig. 5.**
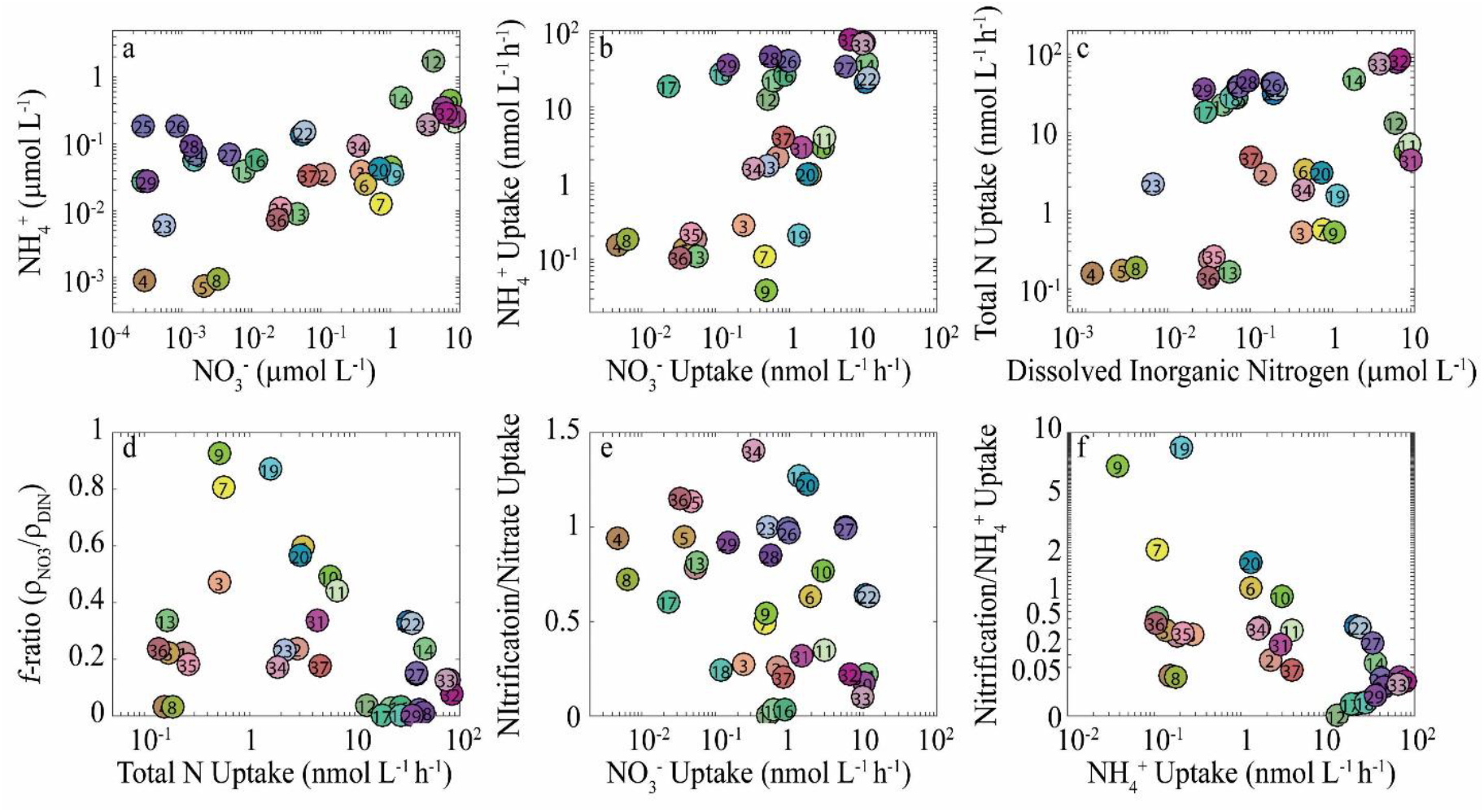
Summary of steady-state conditions for each model simulation. Each data point represents conditions for a specific parameter set.

To assess the accuracy of each nutrient uptake equation, each model configuration was first run to steady-state. At steady-state, simulated *in situ* mixed layer nitrate and ammonium uptake were quantified at a depth of 10-m using the equations within NEMURO. At the same time simulated “bottle incubations” were conducted on water from 10-m depth by: 1) turning off model physics (i.e., turning off the sinking and mixing subroutines in the model), 2) “spiking” the “bottle” by adding additional ^15^N labeled nitrate or ammonium to the appropriate model state variable, 3) running the bottle incubation for a period of 4 or 24 hours, and then 4) quantifying the ^15^N isotope ratio of all particulate matter (phytoplankton, zooplankton, and detritus) at the end of the incubation. Based on my experience with true *in situ* incubations, I “spiked” the simulated incubations with 10% of ambient nutrient concentrations if nutrient concentrations were >100 nmol L^-1^, but spiked with a constant 10 nmoL L^-1^ if ambient nutrients were <100 nmol L^-1^.

ρ_0_, ρ_kan_, ρ_reg_, ρ_0,IS_, ρ_kan,is_, and ρ_reg,IS_ were then calculated from Eqs. 1, 3, 5, 6, 13, and 14 (for both nitrate and ammonium uptake) and compared to “true” model nitrate and ammonium uptake calculated either within the modeled bottle or the model water column. For ρ_kan_, ρ_reg_, ρ_kan,IS_, ρ_reg,IS_ I assumed that a (the ratio of nutrient regeneration rate to nutrient uptake rate) was equal to N_amb_/(N_amb_+N_spk_). This assumption was made because it is rare to have information available to make any better guess than the null assumption that the system is near steady-state (prior to the perturbation of adding labeled substrate). I assumed that uncertainty in a was equal to ±0.5. For ρ_0,IS_, ρ_kan,IS_, and ρ_reg,IS_, I assumed (regardless of what half-saturation constant was used in the model) that K_NO3_ = 0.1 μmol L^-1^ and K_NH4_ = 0.05 μmol L^-1^ based on prior data syntheses (Beltrán-Heredia et al. 2017; Edwards et al. 2012; Harrison et al. 1996; Mutshinda et al. 2017). It must be noted, however, that some studies have called into question the applicability of assuming any Michaelis-Menten half-saturation constant (Bonachela et al. 2011; Tang and Maggi 2012), and that even if this approach is an accurate model, half-saturation constants vary by several of orders of magnitude. I thus introduce variables L10_KNO3_ and L10_KNH4_, which are the log base 10 transformations of K_NO3_ and K_NH4_, respectively, and set L10_KNO3_ = −1 ± 1 and L10_KNH4_ = −1.3 ± 1.

Since Eqs. 3 and 13 (ρKan and ρ_reg_) assume that the nutrient concentrations are at steady state in the incubation bottle, it is possible that evaluating the equations at steady state may unfairly favor ρKan and ρ_reg_. I thus also conducted non-steady-state simulations. These non-steady-state simulations had broadly similar results to the steady-state simulations (see online appendix).

### Results of simulated incubations compared to nutrient uptake within bottles

To investigate the efficacy of each nutrient uptake equation, I ran a series of NEMURO+^15^N simulations and conducted “simulated incubations” by turning off mixing and adding ^15^N-labeled nutrient tracers (^15^NO_3_^-^ or ^15^NH_4_^+^). When estimating nitrate uptake from 4-h simulated incubations ρ_0_, ρ_kan_, and ρ_reg_ all performed similarly well at low nitrate uptake rates (<2 nmol N L^-1^ h^-1^). At these low rates, ρ_0_ and ρk an showed a consistent, very weak negative bias, while ρ_reg_ typically had a similar small negative misfit, but also overestimated true uptake by a similar amount for a couple of simulations (Fig. 6 a-c). At higher nitrate uptake rates results began to diverge. Above 2 nmol N L^-1^ h^-1^ both ρ_0_ and ρ_kan_ underestimated uptake, at times by >50%. The underestimate of true uptake rates was most concerning for ρ_0_, with which uncertainty estimates gave a false sense of confidence in these underestimated values. For one simulation, even the upper limit of the confidence interval was less than 50% of the true value. By contrast, the greater uncertainty limits derived using ρ_kan_ (which has higher uncertainty because it incorporates uncertainty in a, the fraction of nutrients that are regenerated) bracketed the true value for every simulation except one. In contrast, ρ_reg_ started to slightly overestimate true uptake at ~1 nmol N L^-1^ h^-1^, and substantially overestimate it over 2 nmol N L^-1^ h^-1^. However, for all simulations the uncertainty estimates for ρ_reg_ bracketed the true value.

**Fig. 6.**
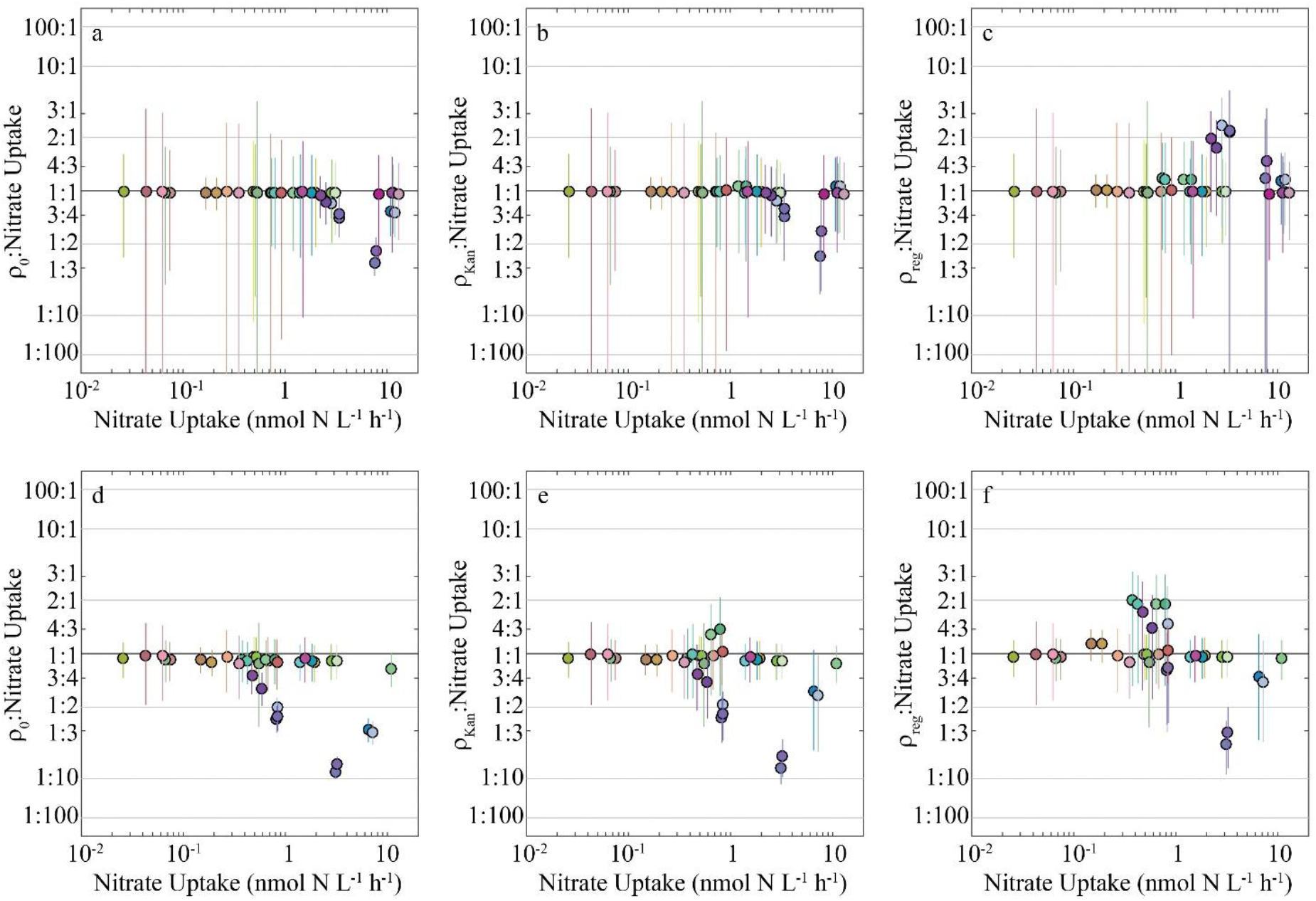
Comparison of nitrate uptake results for 4-h (a – c) and 24-h (d-f) nitrate uptake measurements from steady-state NEMURO+^15^N model runs, calculated using ρ_0_ (a, d, Eq. 1), ρKan (b, e, Eq. 3), and ρ_reg_ (c, f, Eq. 14). Y-axis is the ratio of calculated uptake to “actual” uptake computed inside simulated bottles in the NEMURO+^15^N model. Colors represent different parameter sets and are identical to colors in Fig. 5.

When running 24-h incubations similar patterns emerged, but the errors associated with ρ_0_, ρ_kan_, and ρ_reg_ became more severe (Fig. 6d-f). For ρ_0_, underestimates reached a ratio of 1:10 and uncertainty ranges actually shrank relative to the 4-h incubations. This led to 9 simulations in which uncertainty ranges did not bracket the true values. Estimates based on ρ_kan_ were largely similar to those based on ρ_0_, although the greater uncertainty estimates for ρ_kan_ led to these estimates only failing to bracket the true values for 7 simulations. In contrast, ρ_reg_ overestimated the true uptake rate by a ratio of ~2:1 for several simulations with nitrate uptake of 10 – 20 μmol m^-3^ d^-1^ and underestimated it by a ratio of ~1:4 for two simulations at high nitrate uptake rates (the only simulations for which uncertainty did not bracket the true values).

Greater differences emerged between ρ_0_ and the two equations designed to account for isotope dilution when used to estimate ammonium uptake (Fig. 7). At uptake rates >1 nmol L^-1^ h^-1^, 4-h ρ_0_ underestimates were pronounced, typically at a ratio of 1:2 and uncertainty estimates seldom bracketed the true value. In contrast, ρ_kan_ and ρ_reg_ showed no consistent over- or under-estimate bias and ρ_kan_ uncertainty estimates always bracketed the true value, while ρ_reg_ uncertainty estimates failed to bracket the true value for a single simulation. For 24-h incubations, ammonium uptake estimates were much worse (Fig. 7d-f). The negative bias for ρ_0_ was substantial for most simulations and reached values lower than 1:10. Uncertainty estimates also suggest great confidence in these substantially biased estimates. ρ_kan_ also consistently underestimated the true values, although to a lesser extent. For ρ_reg_ the misfits were not consistent, with a negative misfit on most simulations, but a positive misfit on some. At high nutrient uptake rates (>10 nmol L^-1^ h^-1^) where all three equations underestimated true nutrient uptake, ρ_re_g performed the best.

**Fig. 7.**
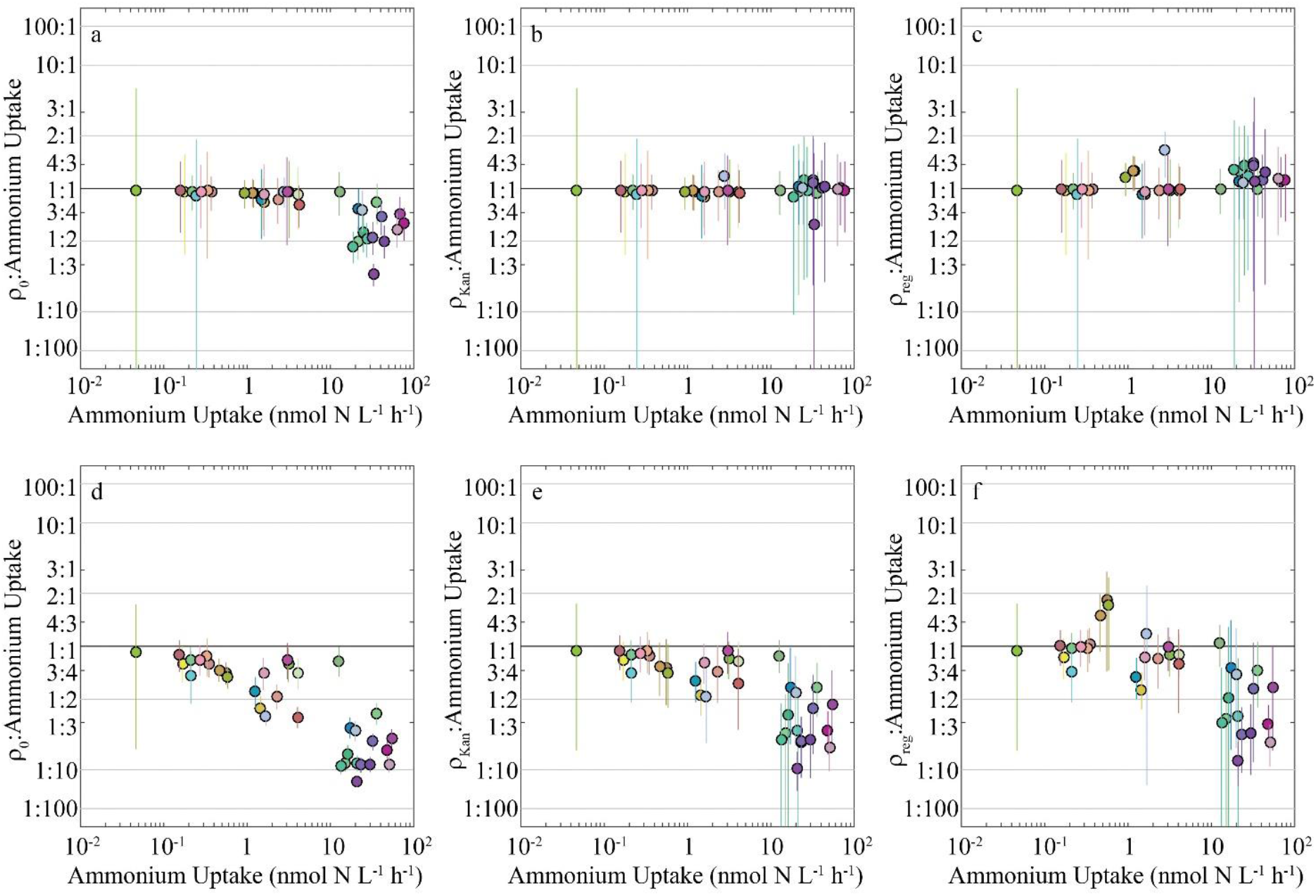
Comparison of ammonium uptake results for 4-h (a – c) and 24-h (d-f) ammonium uptake measurements from steady-state NEMURO+^15^N model, calculated using ρ_0_ (a, d, Eq. 1), pKan (b, e, Eq. 3), and ρ_reg_ (c, f, Eq. 14). Y-axis is the ratio of calculated uptake to “actual” uptake computed inside simulated bottles in the NEMURO+^15^N model. Colors represent different parameter sets and are identical to colors in Fig. 5.

### Results of simulated incubations compared to simulated in situ conditions

The above results compared nutrient uptake rates predicted based on bottle incubations to the actual nutrient uptake in those simulated bottles. However, investigators are often interested in determining *in situ* nutrient uptake rates rather than uptake rates within incubation bottles. I thus compared nutrient uptake estimates calculated from simulated bottle incubations (using ρ_0_, ρ_kan_, ρ_reg_, ρ_0,is_, ρ_kan,is_, and ρ_reg,is_) to nutrient uptake rates in the model at the depths from which the bottles were sampled. The *in situ* correction in Eq. 5, 6, and 14 leads to a downward revision of nutrient uptake estimates relative to ρ_0_, ρ_kan_, and ρ_reg_, especially at low nutrient concentrations when I used a minimum simulated nutrient spike of 10 nmol L^-1^ that exceeded 10% of ambient nutrients.

When comparing nitrate uptake rates from 4-h incubations using ρ_0_ and ρ_0,is_ to simulated *in situ* nutrient uptake the patterns were fairly complex (Fig. 8a,d). At low uptake rates ρ_0_ tended to overestimate *in situ* nitrate uptake, but was reasonably accurate at high uptake rates. ρ_0,is_, was fairly accurate at low uptake rates, but typically underestimated nitrate uptake at high uptake rates. These contrasting patterns were driven by differing causes. True nitrate uptake in the simulated bottles was typically higher than true nitrate uptake in the simulated ocean, due to the tracer addition. Thus when ρ_0_ was accurate at estimating nitrate uptake in the bottle, the correction applied by ρ_0,is_ led to more accurate estimation of *in situ* nutrient uptake. However, when ρ_0_ substantially underestimated true nitrate uptake in the bottle, it was often closer to the actual values *in situ*. This led the correction applied by ρ_0,is_ to underestimate true *in situ* nitrate uptake. Similar patterns were seen when comparing ρ_kan_ and ρ_kan,is_ (Fig. 8b,e), although the greater uncertainty estimates for ρ_kan_ relative to ρ_0_ (and ρ_kan,is_ relative to ρ_0,is_) more accurately reflected uncertainty in uptake rate measurements. For ρ_reg_ and ρ_reg,is_ the greater agreement between calculated and true nutrient uptake within the bottles led to generally greater agreement between ?reg,is and true *in situ* uptake than between ρ_re_g and true *in situ* uptake (Fig. 8c,f).

**Fig. 8.**
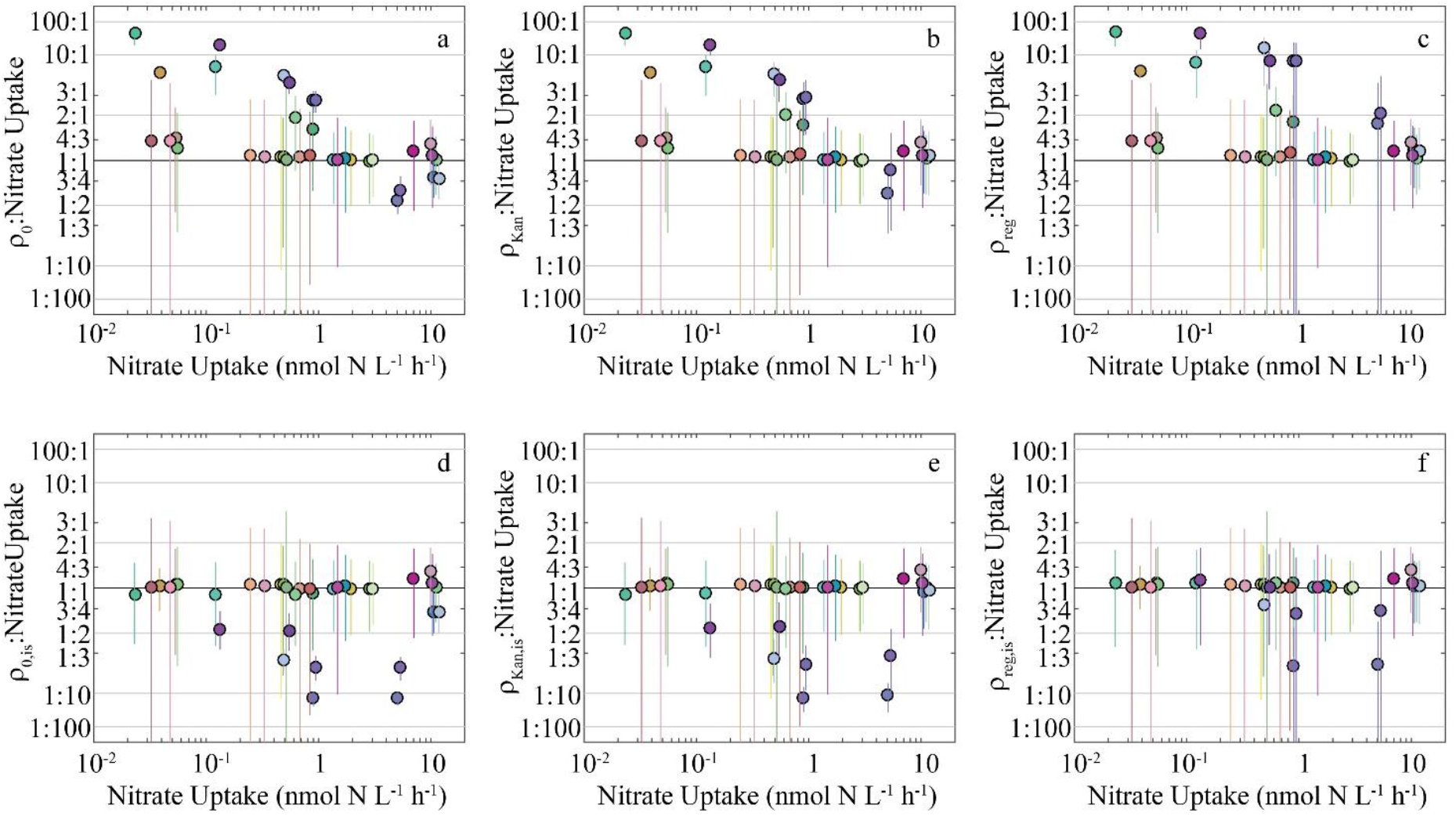
Comparison of simulated calculated nitrate uptake to simulated natural uptake rates. Calculated nitrate uptake rate is computed using ρ_0_ (a), ρ_kan_ (b), ρ_reg_ (c), ρ_0,is_ (d), ρ_kan,is_ (e), ρ_reg,is_ (f). Y-axis is the ratio of calculated uptake after 4-h simulated incubations to “actual” uptake computed in the 1-D NEMURO+^15^N model. Colors represent different parameter sets and are identical to colors in Fig. 5.

For simulations comparing *in situ* ammonium uptake to estimates from ρ_0_ and ρ_0,is_ the results were similar to those for nitrate uptake. ρ_0,is_ was an improvement when uptake rates were low, but led to slightly worse results when uptake was high (Fig. 9a,d). By contrast ρ_kan,is_ was typically a very good estimate of *in situ* uptake rates (Fig. 9e). The uncertainty estimates only failed to bracket the true value during two simulations and the worst disagreements led to only a roughly 1:2 ratio of estimated to true uptake rates. ρ_reg_ and ρ_reg,is_ showed similar performance (Fig. 9c,f). ρ_reg_ was an overestimate at low ammonium uptake rates. ρ_reg,is_ typically showed good agreement with uncertainty estimates only failing to bracket the true values for one simulation.

**Fig. 9.**
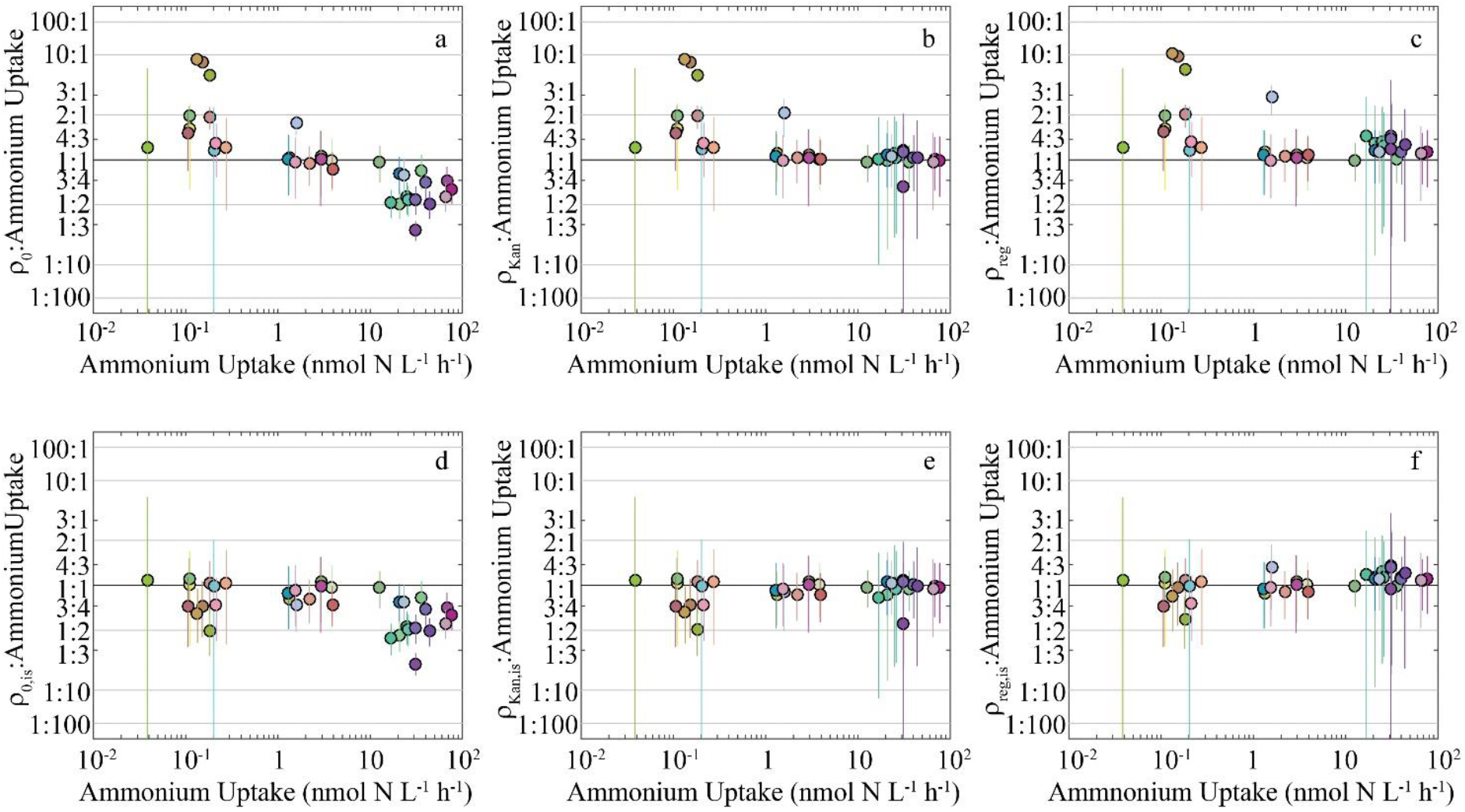
Comparison of simulated calculated ammonium uptake to simulated natural uptake rates. Calculated ammonium uptake rate is computed using ρ_0_ (a), ρ_kan_ (b), ρ_reg_ (c), ρ_0,is_ (d), ρ_kan,is_ (e), ρ_reg,is_ (f). Y-axis is the ratio of calculated uptake after 4-h simulated incubations to “actual” uptake computed in the 1-D NEMURO+^15^N model. Colors represent different parameter sets and are identical to colors in Fig. 5.

## APPENDIX 2: Assessment using diel uptake experiments

### Field methods

To investigate whether nutrient uptake calculation results based on model situations were applicable to field measurements, I conducted diel NO_3_^-^ and NH_4_^+^ uptake experiments in the Costa Rica Dome (CRD) and the oligotrophic central Gulf of Mexico (GoM). The CRD is an open ocean upwelling ecosystem with relatively high NO_3_^-^ concentrations, abundant populations of picophytoplankton (particularly *Synechococcus)*, and high biomass of mesozooplankton and higher trophic levels (Selph et al. 2016; Stukel et al. 2018a). The central Gulf of Mexico is an oligotrophic region with very low nutrient concentrations, deep chlorophyll maxima, and an important role for the nitrogen fixing cyanobacterium *Trichodesmium* (Gomez et al. 2018; Mulholland et al. 2006; Muller-Karger et al. 2015).

In both regions, I conducted diel uptake experiments, consisting of six consecutive 4-h incubations coincident with replicated 24-h incubations. Samples were incubated in 2.2-L (CRD) or 2.7-L (GoM) polycarbonate bottles, gently filled using silicon tubing. All 8 – 10 bottles for an experiment were filled from Niskin bottles from the same depth on the same cast. 24-h bottles and a single 4-h bottle were initially spiked with ^15^N-labeled NO_3_^-^ or NH_4_^+^. I added 100 nmol L^-1 15^NO_3_^-^ (final concentration) in the CRD and 8-10 nmol ^15^NO_3_^-^ or 5 – 10 nmol ^15^NH_4_^+^ in the GoM. These concentrations were chosen based on expectations for 10% of *in situ* nutrient concentrations, although subsequent measurements showed that in the GoM, NO_3_^-^ and NH_4_^+^ concentrations were occasionally as low as 10 and 54 nmol L^-1^, respectively. All bottles were placed in a deckboard incubator shaded with blue acrylic to match *in situ* light levels and cooled with flow-through surface seawater. Four hours later, the first 4-h sample was removed from the incubator and immediately filtered through a pre-combusted GF/F filter and the second 4-h sample was spiked. This process was repeated every four hours until twenty four hours had passed. At this point the final 4-h sample and the two or three 24-h samples were simultaneously removed and filtered. Isotope dilution could not be measured on the samples, because of the low nutrient concentration. Although the six 4-h incubations form a time-series that is comparable to conditions experienced in the 24-h incubations, I cannot exclude the possibility that incubation artifacts varied between bottles, particularly when tracer additions were relatively large compared to ambient nutrient concentration. Samples were frozen a-80°C immediately after filtration. On land they were dried and analyzed for δ^15^N and PON at the UC Davis Stable Isotope Facility. This approach yielded a series of 4-h samples covering the same period of time as the 24-h samples. An unbiased equation for estimating nutrient uptake should thus generate identical estimates for nutrient uptake if calculated using the 24-h incubations or the sum of uptake calculated during the series of 4-h samples.

When computing ρ_Kan_ and ρ_reg_, I assumed that a = 0 for NO_3_^-^ uptake in the CRD, because independent nitrification rate measurements showed that nitrification (i.e. recycling of NO_3_^-^) was negligible (Buchwald et al. 2015). In the GoM, I assumed that a = 1 for NO_3_^-^ and NH_4_^+^ uptake, because a deep nutricline (> 100 m) and very low euphotic zone nutrient concentrations suggested that upwelling could not support measured NO_3_^-^ and NH_4_^+^ uptake rates. The CRD data was published in Stukel et al. (2016). The GoM data is derived from two NOAA-funded cruises in May 2017 and May 2018.

### Results of diel incubation experiments

To determine if these results derived from simulations were reflected in actual nutrient uptake incubations, I conducted a series of diel uptake experiments in the CRD and GoM. Experiments included 24-h incubations in combination with 4-h incubations throughout the same period for the same phytoplankton community incubated at identical conditions. Results showed a strong diel cycle in nitrate uptake in both the CRD and GoM (Fig. 10). Nitrate uptake was substantially greater during daylight hours. Results also showed relatively good agreement between nitrate uptake calculated from 24-h or from the series of 4-h incubations when using ρ_reg,is_. Agreement was particularly good for the CRD samples (Fig. 10a-c), when nutrient recycling and isotope dilution were minimal as a result of reasonably high nitrate concentrations and negligible euphotic zone nitrification (Buchwald et al. 2015; Stukel et al. 2016). Greater discrepancy was found in the GoM; estimates from 24-h incubations occasionally exceeded and occasionally were less than 4-h incubations.

**Fig. 10.**
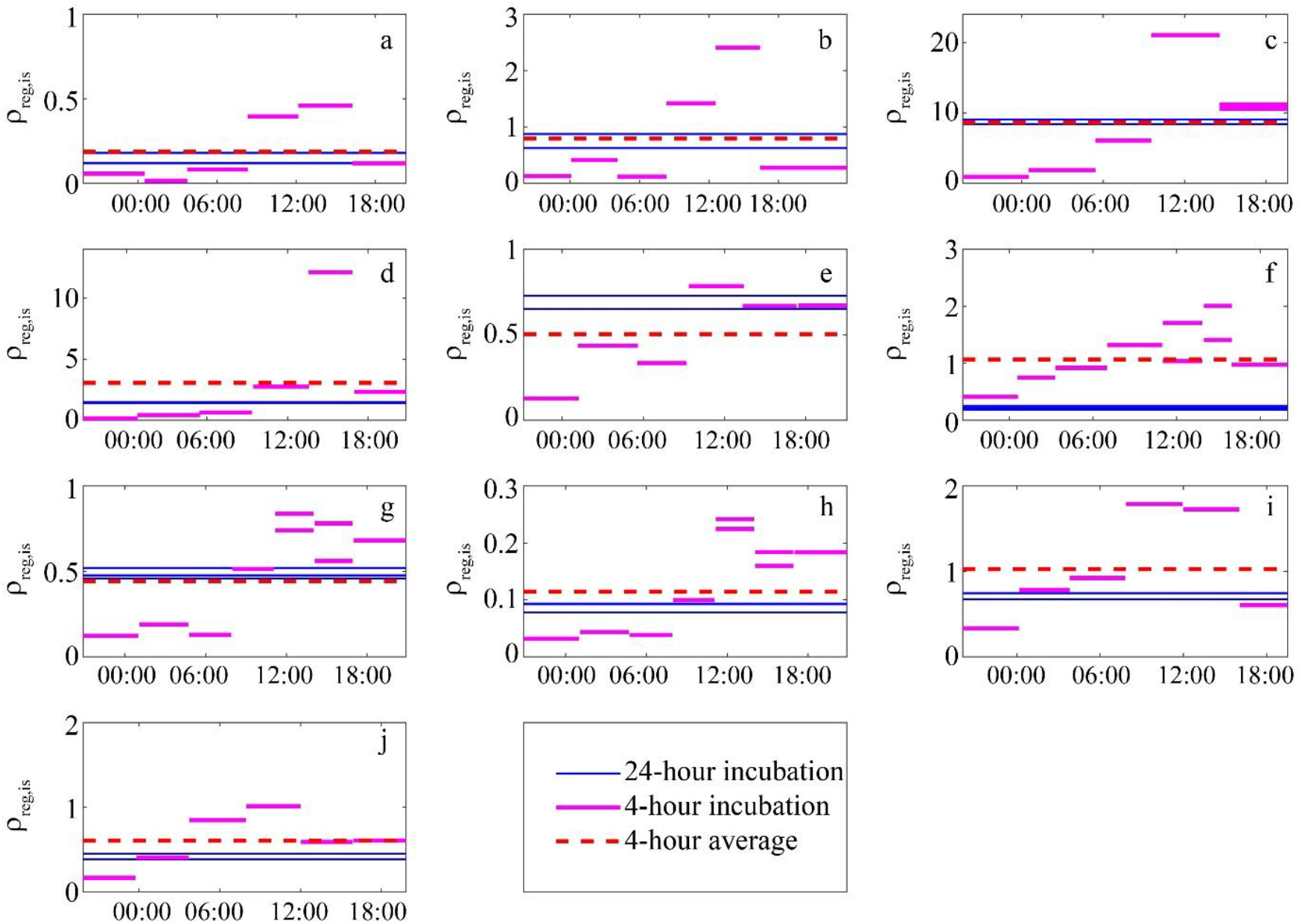
Diel nitrate uptake experiments computed using ρ_reg,is_ (nmol N L^-1^ h^-1^). a-c are from CRD. d-j are from GoM.

Results were, however, strikingly different for ammonium uptake measurements in the GoM (Fig. 11). 4-h incubation estimates showed no distinct diel periodicity and consistently exceeded estimates from 24-h incubations by a factor of 4.3-6.2 when computed with ρ_reg_. Comparisons between 4-h and 24-h ammonium uptake rates computed with ρ_kan_ and ρ_0_ were slightly worse, with ratios ranging from 4.9-6.3 and 5.1-6.3, respectively. The uncertainty estimates also shrank substantially with ρ_0_, yielding a false sense of certainty in the incorrect data.

**Fig. 11.**
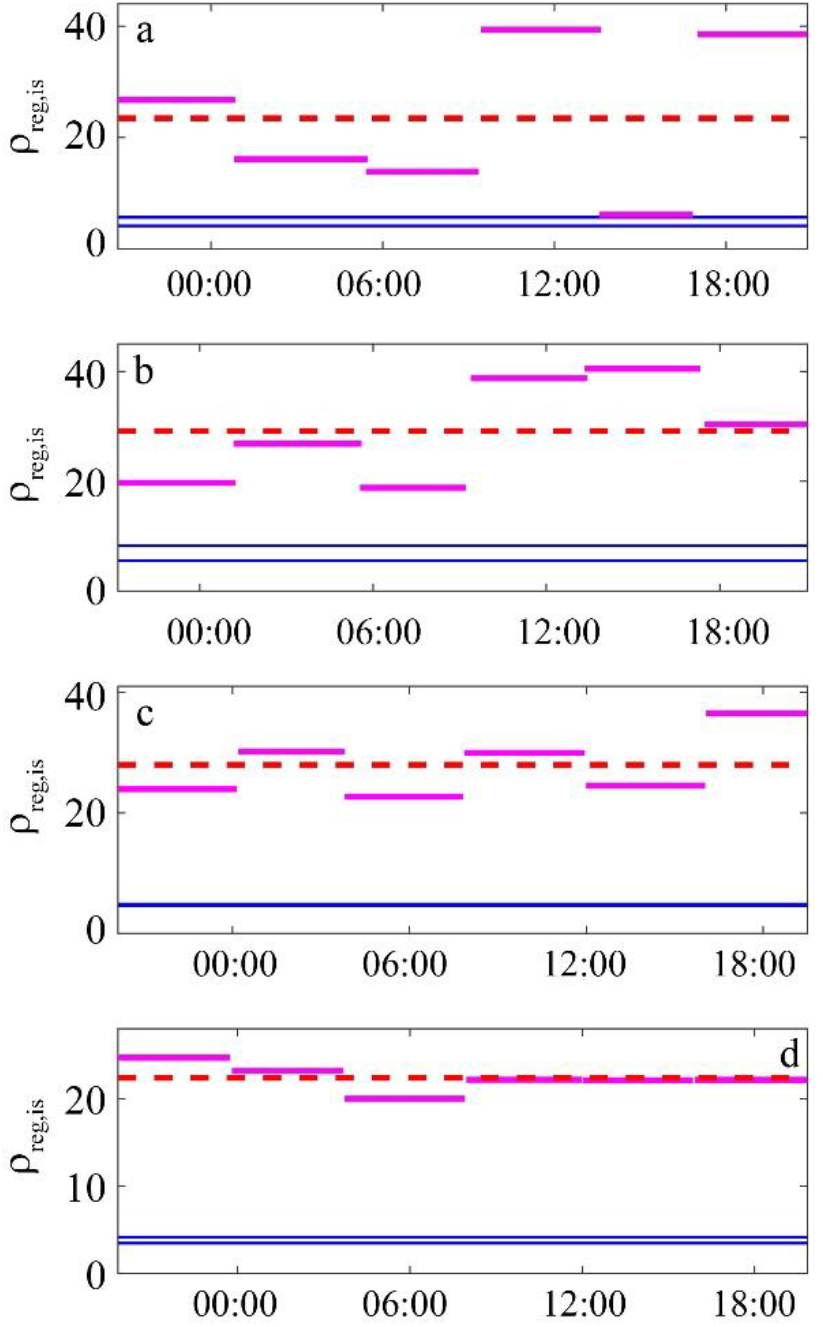
Diel ammonium uptake experiments from the GoM computed using ρ_reg,is_ (nmol N L^-1^ h^-1^). Symbols are same as Fig. 10.

When comparing across all measurements (ammonium uptake and nitrate uptake), the geometric mean of the ratio of 4-h nutrient uptake to 24-h nutrient uptake was 2.3 for ρ_0_ and 2.0 for both ρ_kan_ and ρ_reg_ (Fig. 3). Although all equations failed to deliver unbiased estimates for ammonium uptake in the GoM, nitrate uptake estimates agreed to within the errorbars for 9 out of 10 experiments with ρ_kan_ and ρ_reg_, but only 8 out of 10 for ρ_0_ (using symmetric confidence limits, Fig. 3a-c).

The large mismatch between 4-h and 24-h ammonium uptake measurements deserves greater investigation. Notably, when using more accurate asymmetric confidence limits (Fig. 3d-f), ρ_reg_ clearly outperforms ρ_0_ and ρ_kan_, because the uncertainty limits for two of the data points reflect a large degree of uncertainty in the upper limit of nitrate uptake. The cause of the apparent discrepancy between mean estimates of ammonium uptake based on 4- or 24-h incubations is easily seen in the data. I_P_(T) was similar after the 4-h and 24-h incubations, suggesting that the isotopes had reached equilibrium in the particulate pool after only 4 hours. However, this equilibrium value did not reach the asymptote predicted by Eq. 15 as would be expected based on the equations used to derive ρ_reg_. Rather, it reached an equilibrium that reflected the presence of unlabeled nitrate (in addition to ammonium and particulate nitrogen). More specifically, the final value for I_P_(T) was approximately equal to:

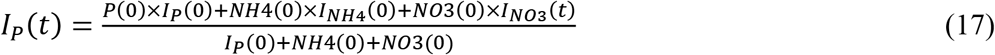

This suggests that when phytoplankton are deriving an essential element from two nutrient sources and substantial nutrient regeneration is occurring within the incubation experiments a modified equation may be necessary to account for these issues.

## ACKNOWLEDGMENTS

I am very grateful to my colleagues in the CRD FluZiE and NOAA Restore Act Bluefin Tuna Project who assisted in sample collection, especially Thomas Kelly, Karen Selph, Michael Landry, and Angela Knapp. The NEMURO+^15^Ncode used in this manuscript can be downloaded from GitHub at: https://github.com/stukel-lab/NEMURON15-CalcNUp. The Matlab scripts and Excel spreadsheets used to calculate uptake rates and related uncertainty are all available on GitHub at: https://github.com/stukel-lab/NUpCalculations and as online supplemental files with this manuscript. Funding for this project came from National Science Foundation Biological Oceanography grant #1851347 and a National Oceanic and Atmospheric Administration’s RESTORE Program Grant (Project Title: Effects of nitrogen sources and plankton food-web dynamics on habitat quality for the larvae of Atlantic bluefin tuna in the Gulf of Mexico) under federal funding opportunity NOAA-NOS-NCCOS-2017-2004875. https://restoreactscienceprogram.noaa.gov/funded-projects/bluefin-tuna-larvae.

## Notes

### Competing Interest Statement

The authors have declared no competing interest.

