## Supplemental Text for "Investigating equations for measuring dissolved inorganic nutrient uptake in oligotrophic conditions"

Michael R. Stukel[1,2]

[1] Dept. of Earth, Ocean & Atmospheric Science, Florida State University, Tallahassee, FL, USA

[2] Center for Ocean-Atmospheric Prediction Studies, Florida State University, Tallahassee, FL, USA

### Supplemental Text

Since Eqs. 3 and 13 ($\rho_{Kan}$ and $\rho_{reg}$) assume that the nutrient concentrations are at steady state in the incubation bottle, it is possible that evaluating the equations at steady state may unfairly favor $\rho_{Kan}$ and $\rho_{reg}$. I thus also conducted non-steady-state simulations. These simulations mimicked the behavior that might be expected if differing rates of cloud cover or storm induced mixing drive phytoplankton variability through changes in surface irradiance, mixed layer depth, and mixed layer vertical eddy diffusivity (Supp. Fig. 1), while using each of the parameter sets in Fig. 5. Each non-steady state simulation was initialized from the previous steady-state solution for that respective parameter set. It was then run for two years using the temporally-varying forcing time-series. During the second year of the run, I "sampled" the model every two weeks to calculate $NO_3Up$, $NH_4Up$, $\rho_0$, $\rho_{kan}$, $\rho_{reg}$, $\rho_{0,IS}$, $\rho_{kan,IS}$ and $\rho_{reg,IS}$. Variability resulting from these simulations is shown in Supp. Fig. 2.

Results were largely similar to those found under steady-state conditions (Supp. Fig. 3). With respect to 4-h nitrate uptake experiments, $\rho_0$ and $\rho_{kan}$ had similar and relatively mild negative biases that grew stronger at higher nitrate uptake rates (Supp. Fig. 3a,b). $\rho_{kan}$ showed less misfit than $\rho_0$ and larger uncertainty ranges leading to a better agreement with the data. In contrast, $\rho_{reg}$ was biased slightly high, but (as for $\rho_{kan}$) its uncertainty estimates typically bracketed the true value (Supp. Fig. 3c). For 4-h ammonium uptake rate experiments, $\rho_0$ again exhibited a consistent negative bias and misfit with the data again grew stronger with increasing ammonium uptake rates (Supp. Fig. 3d). $\rho_{kan}$ and $\rho_{reg}$ showed no substantial bias for most simulations, but with a few parameter sets, they showed slight overestimates (Supp. Fig. 3e,f).

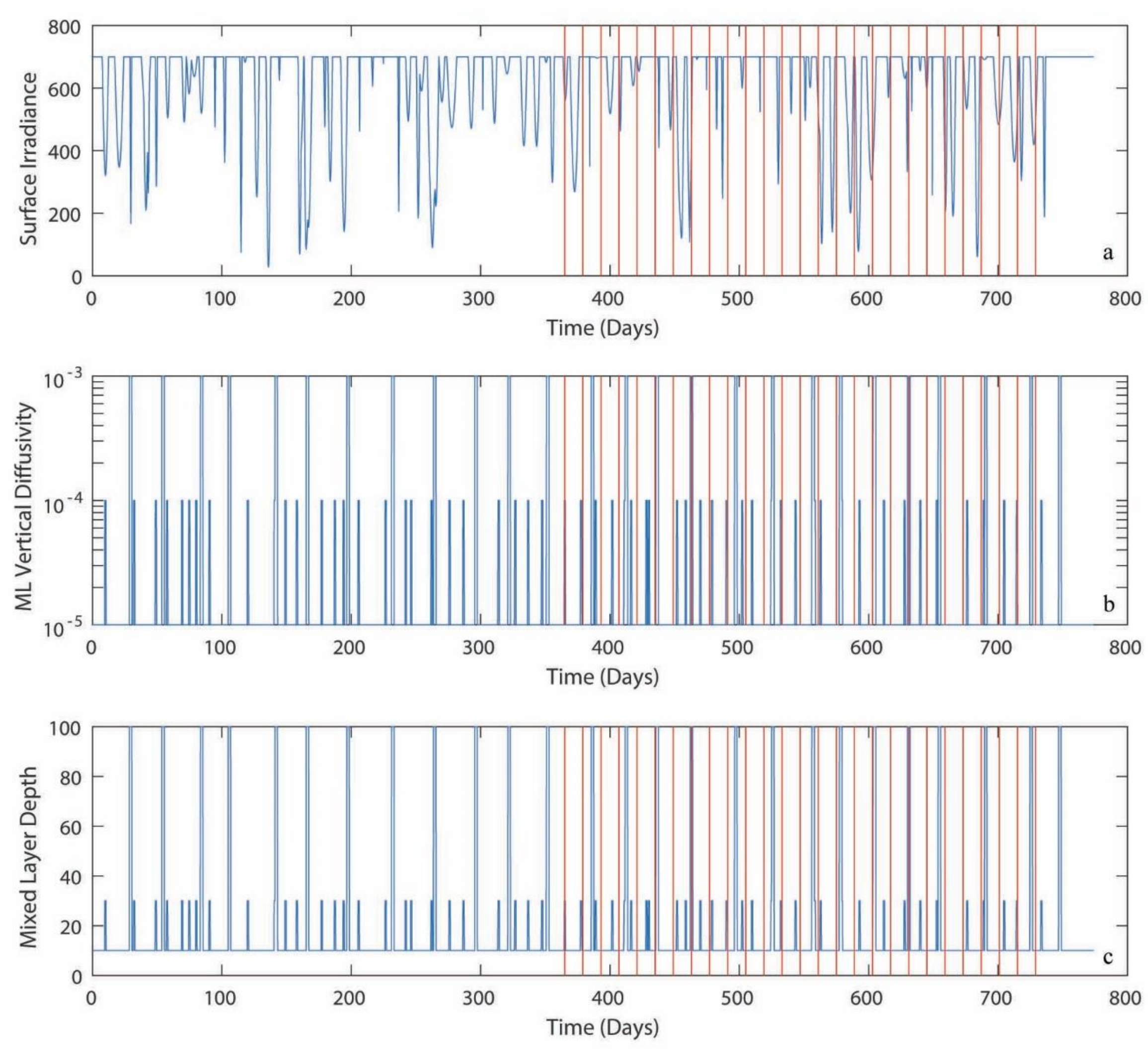


Supp. Fig. 1 – Time-series of surface irradiance (a), mixed layer vertical diffusivity (b), and mixed layer depth (c) used in non-steady-state simulations.  Red vertical lines mark sampling points.

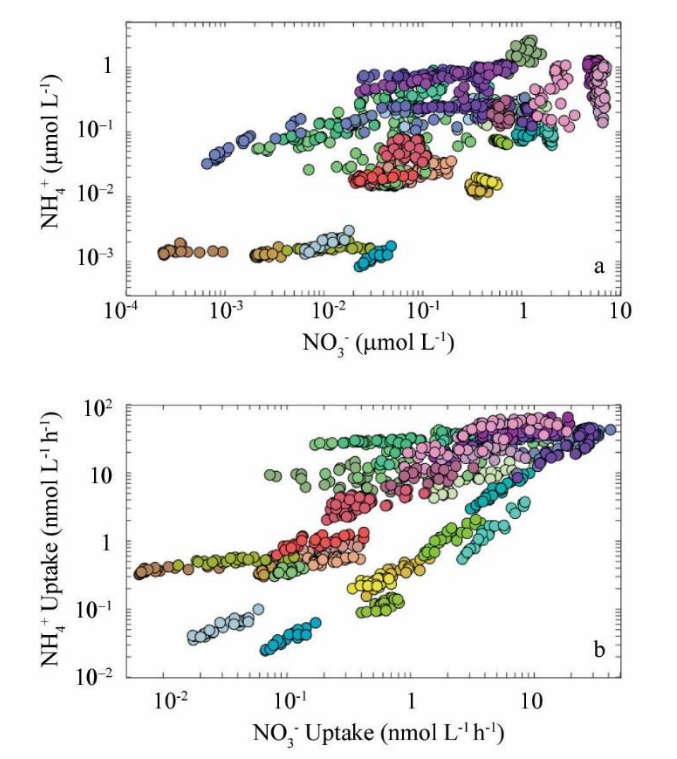


Supp. Fig. 2 – Nutrient concentrations (a) and nutrient uptake rates (b) at "sampling time points" for non steady-state simulations. Colors represent different parameter sets and are identical to colors in Fig. 3.

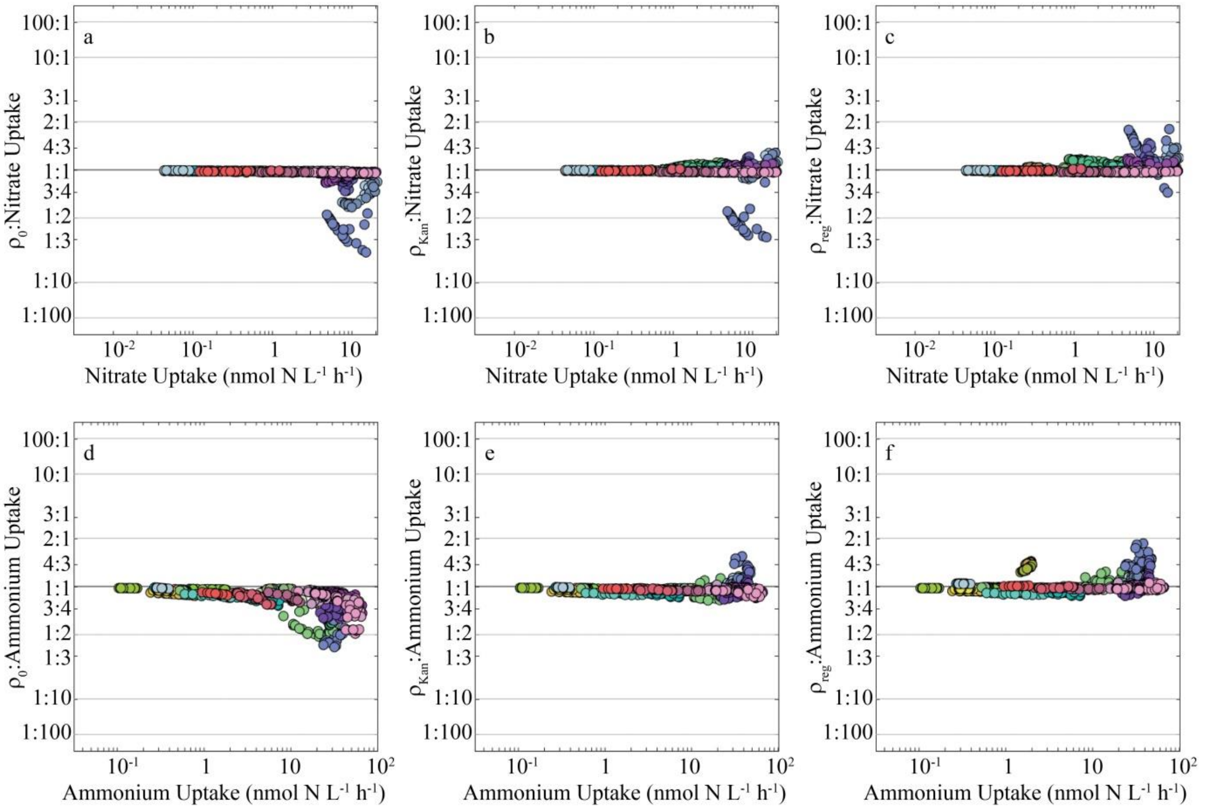


Supp. Fig. 3 – Comparison of nitrate uptake (a – c) and ammonium uptake (d – f) results for 4-h nutrient uptake measurements from non-steady-state NEMURO+$^{15}$N model, calculated using $\rho_0$ (a, d, Eq. 1), $\rho_{Kan}$ (b, e, Eq. 3), and $\rho_{reg}$ (c, f, Eq. 14). Y-axis is the ratio of calculated uptake to "actual" uptake computed inside simulated bottles in the NEMURO+$^{15}$N model. Colors represent different parameter sets and are identical to colors in Fig. 3.
