## Supplementary Appendix S1 for "Investigating equations for measuring dissolved inorganic nutrient uptake in oligotrophic conditions"

**Supplementary Appendix S1 – Derivation of $\rho_{reg}$ (nutrient uptake with regeneration and temporally-varying isotope dilution).**

If we assume (as was assumed for the derivations of $\rho_0$ and $\rho_{kan}$) that nutrient uptake ($\rho$) is constant throughout the duration of a nutrient uptake experiment, we can define a series of differential equations defining the rate of change of the substrate (S), the particulate organic matter in the incubation (P) and the total isotope-label in the substrate ($^{15}S$) and particulate organic matter ($^{15}P$):

$$\frac{\partial S(t)}{\partial t} = (a-1)\rho \qquad \text{(A1)}$$

$$\frac{\partial P(t)}{\partial t} = (1-a)\rho \qquad \text{(A2)}$$

$$\frac{\partial\, ^{15}S(t)}{\partial t} = a\rho \times I_P(t) - \rho \times I_S(t) \qquad \text{(A3)}$$

$$\frac{\partial\, ^{15}P(t)}{\partial t} = \rho \times I_S(t) - a\rho \times I_P(t) \qquad \text{(A4)}$$

where a is the fraction of nutrient uptake that gets recycled in the incubation (as in Kanda et al., 1987), IP(t) is the isotope ratio of P at time t and IS(t) is the isotope ratio of S at time t. It follows that:

$$\frac{\partial I_S(t)}{\partial t} = \frac{a\rho \times I_P(t) - \rho \times I_S(t)}{S(t)} \qquad \text{(A5)}$$

$$\frac{\partial I_P(t)}{\partial t} = \frac{\rho \times I_S(t) - a\rho \times I_P(t)}{P(t)} \qquad \text{(A6)}$$

Ideally, we should solve the above system of equations to determine an unbiased estimate of $\rho$. Unfortunately, while the above set of differential equations has a closed form solution, the solution cannot be solved for $\rho$. However, we can make the simplifying assumption that substrate concentration and particulate organic matter concentration remain approximately constant during the incubation ($S(t) \approx S(0)$ and $P(t) \approx P(0)$). I note that this approximation will be exactly true if a = 1. It should also be reasonable anytime that $\rho$ is constant throughout the incubation. I also note that S(0) is equal to Namb + Nspk from Eq. 2. Given this assumption, and conservation of $^{15}N$ in the incubation, we can show that:

$$I_S(t) = \frac{I_P(0) \times P(0) + I_S(0) \times S(0) - I_P(t) \times P(0)}{S(0)} \qquad \text{(A7)}$$

Substituting A7 into A6 and rearranging gives us:

$$\frac{\partial I_P(t)}{\partial t} = \rho\left(\frac{I_P(0)\times P(0)+I_S(0)\times S(0)}{P(0)\times S(t)} - \frac{P(0)+a\times S(t)}{P(0)\times S(t)} I_P(t)\right) \qquad \text{(A8)}$$

After rearranging:

$$\int \frac{\partial I_P(t)}{\left(\frac{I_P(0)\times P(0)+I_S(0)\times S(0)}{P(0)\times S(0)} - \frac{P(0)+a\times S(0)}{P(0)\times S(0)} I_P(t)\right)} = \int \rho \partial t \qquad \text{(A9)}$$

So:

$$-ln\left(\left|\frac{P(0)+a\times S(0)}{P(0)\times S(0)} I_P(t) - \frac{I_P(0)\times P(0)+I_S(0)\times S(0)}{P(0)\times S(0)}\right|\right) = \frac{P(0)+a\times S(0)}{P(0)\times S(0)} \rho t + C \qquad \text{(A10)}$$

Where C is a constant, which, after solving at time t=0, we can show is equal to:

$$C = -ln\left(\frac{I_S(0)-a\times I_P(0)}{P(0)}\right) \qquad \text{(A11)}$$

Therefore:

$$\rho = \left(ln\left(\frac{I_S(0)-a\times I_P(0)}{P(0)}\right) - ln\left(\frac{I_P(0)\times P(0)+I_S(0)\times S(0)}{P(0)\times S(0)} - \frac{P(0)+a\times S(0)}{P(0)\times S(0)} I_P(t)\right)\right)\left(\frac{P(0)\times S(0)}{P(0)+a\times S(0)}\right)\frac{1}{t} \qquad \text{(A12)}$$

Evaluated at the end of the incubation (t = T), this defines ρ in terms of variables measured in a typical nutrient uptake experiment:

$$\rho = \left(ln\left(\frac{I_S(0)-a\times I_P(0)}{P}\right) - ln\left(\frac{I_P(0)-I_P(t)}{[N_{spk}+N_{amb}]} + \frac{I_S(0)-a\times I_P(t)}{P}\right)\right)\left(\frac{P\times[N_{spk}+N_{amb}]}{P+a\times[N_{spk}+N_{amb}]}\right)\frac{1}{T} \qquad \text{(A13)}$$

Eq. A13 should be an unbiased estimate of nutrient uptake rates if S and P remain constant throughout the incubation. This will always be true when nutrient regeneration in the bottle is complete (a=1) and the labeled nutrient is the only form of that element being utilized by organisms during the experiment. Constancy of S and P are also implied by the assumption of constant ρ throughout the incubation experiments made in the derivations of Dugdale and Goering (1967) and Kanda et al. (1987). However, it is a potentially biased estimate when $a \neq 1$ and the concentrations of nutrients change during the incubation.
