## Supplementary Appendix S2 for "Investigating equations for measuring dissolved inorganic nutrient uptake in oligotrophic conditions"

### Supplementary Appendix S2 – Nutrient uptake uncertainty derivations.

All of these uncertainty equations have been implemented into Excel spreadsheets and Matlab scripts that are available as supplementary material with this manuscript.

#### Supplementary Appendix S2.1 – Uncertainty equations for $\rho_0$

The uncertainty in nutrient uptake as calculated by Eq. 1 can be computed as:

$$\sigma_{\rho_0} = \sqrt{\left(\frac{\partial\rho_0}{\partial P}\right)^2 \sigma_P{}^2 + \left(\frac{\partial\rho_0}{\partial T}\right)^2 \sigma_T{}^2 + \left(\frac{\partial\rho_0}{\partial I_P(0)}\right)^2 \sigma_{I_P(0)}{}^2 + \left(\frac{\partial\rho_0}{\partial I_P(T)}\right)^2 \sigma_{I_P(T)}{}^2 + \left(\frac{\partial\rho_0}{\partial I_{spk}}\right)^2 \sigma_{I_{spk}}{}^2 + \left(\frac{\partial\rho_0}{\partial I_{amb}}\right)^2 \sigma_{I_{amb}}{}^2 + \left(\frac{\partial\rho_0}{\partial [N]_{spk}}\right)^2 \sigma_{[N]_{spk}}{}^2 + \left(\frac{\partial\rho_0}{\partial [N]_{amb}}\right)^2 \sigma_{[N]_{amb}}{}^2} \qquad \text{(B1)}$$

where $\partial\rho_0/\partial x$ is the partial derivative of $\rho_0$ with respect to parameter x. These partial derivatives are found by differentiating the equation:

$$\rho_0 = \frac{P}{T} \times \frac{I_P(T) - I_P(0)}{\frac{I_{spk}[N]_{spk} + I_{amb}[N]_{amb}}{[N]_{spk} + [N]_{amb}} - I_P(0)} \qquad \text{(B2)}$$

which is found by substituting Eq. 2 into Eq. 1. The partial derivatives are:

$$\frac{\partial\rho_0}{\partial P} = \frac{\rho_0}{P} \qquad \text{(B3)}$$

$$\frac{\partial\rho_0}{\partial T} = \frac{-\rho_0}{T} \qquad \text{(B4)}$$

$$\frac{\partial\rho_0}{\partial I_P(T)} = \frac{\rho_0}{I_P(T) - I_P(0)} \qquad \text{(B5)}$$

$$\frac{\partial \rho_0}{\partial I_P(0)} = \frac{\rho_0}{I_{S,ex}} - \frac{P}{T}\left(\frac{\rho_0}{I_{S,ex} \times \rho_0}\right) \tag{B6}$$

Where $I_{s,ex} = I_S(0) - I_P(0)$

$$\frac{\partial \rho_0}{\partial [N]_{spk}} = \frac{-\rho_0}{I_{S,ex}} \times \frac{I_{spk} - I_S(0)}{[N]_{tot}} \tag{B7}$$

$$\frac{\partial \rho_0}{\partial [N]_{amb}} = \frac{-\rho_0}{I_{S,ex}} \times \frac{I_{amb} - I_S(0)}{[N]_{tot}} \tag{B8}$$

$$\frac{\partial \rho_0}{\partial I_{amb}} = \frac{-\rho_0}{I_{S,ex}} \times \frac{[N]_{amb}}{[N]_{tot}} \tag{B9}$$

$$\frac{\partial \rho_0}{\partial I_{spk}} = \frac{-\rho_0}{I_{S,ex}} \times \frac{[N]_{spk}}{[N]_{tot}} \tag{B10}$$

Note that for all reasonable combinations of parameters and uncertainties that I tested, the terms associated with Eq. A6 and A9 (uncertainty in the initial isotopic ratio of PON and uncertainty in the isotopic ratio of the ambient nutrient pool) can be neglected with less than a 1% decrease in $\sigma_{\rho 0}$, thus:

$$\sigma_{\rho_0} \approx \sqrt{\left(\frac{\partial \rho_0}{\partial P}\right)^2 {\sigma_P}^2 + \left(\frac{\partial \rho_0}{\partial T}\right)^2 {\sigma_T}^2 + \left(\frac{\partial \rho_0}{\partial I_P(T)}\right)^2 {\sigma_{I_P(T)}}^2 + \left(\frac{\partial \rho_0}{\partial I_{spk}}\right)^2 {\sigma_{I_{spk}}}^2 + \left(\frac{\partial \rho_0}{\partial [N]_{spk}}\right)^2 {\sigma_{[N]_{spk}}}^2 + \left(\frac{\partial \rho_0}{\partial [N]_{amb}}\right)^2 {\sigma_{[N]_{amb}}}^2} \tag{B11}$$

$$\sigma_{\rho_0} \approx \rho_0 \sqrt{\left(\frac{1}{P}\right)^2 {\sigma_P}^2 + \left(\frac{-1}{T}\right)^2 {\sigma_T}^2 + \left(\frac{1}{I_P(T) - I_P(0)}\right)^2 {\sigma_{I_P(T)}}^2 + \left(\frac{-1}{I_{S,ex}} \times \frac{[N]_{spk}}{[N]_{tot}}\right)^2 {\sigma_{I_{spk}}}^2 + \left(\frac{-1}{I_{S,ex}} \times \frac{I_{spk} - I_S(0)}{[N]_{tot}}\right)^2 {\sigma_{[N]_{spk}}}^2 + \left(\frac{-1}{I_{S,ex}} \times \frac{I_{amb} - I_S(0)}{[N]_{tot}}\right)^2 {\sigma_{[N]_{amb}}}^2} \tag{B12}$$

#### Supplementary Appendix S2.2 – Uncertainty equations for $\rho_{kan}$ (nutrient uptake with regeneration)

When nutrient regeneration within the incubation bottle is suspected, more accurate nutrient uptake estimates can be computed using Eqs. 4 and 5:

$$\rho_{kan} = \rho_0 \frac{-1+(1-b)^{1-a}}{(a-1)b} \tag{3}$$

$$b = \frac{\rho_0 \times T}{[N]_{amb}+[N]_{spk}} \tag{4}$$

The uncertainty in nutrient uptake calculated by Eq. 4 can therefore be calculated from:

$$\sigma_{\rho_{kan}} = \sqrt{\left(\frac{\partial \rho_{kan}}{\partial P}\right)^2 \sigma_P{}^2 + \left(\frac{\partial \rho_{kan}}{\partial T}\right)^2 \sigma_T{}^2 + \left(\frac{\partial \rho_{kan}}{\partial I_P(0)}\right)^2 \sigma_{I_P(0)}{}^2 + \left(\frac{\partial \rho_{kan}}{\partial I_P(T)}\right)^2 \sigma_{I_P(T)}{}^2 + \left(\frac{\partial \rho_{kan}}{\partial I_{spk}}\right)^2 \sigma_{I_{spk}}{}^2 + \left(\frac{\partial \rho_{kan}}{\partial I_{amb}}\right)^2 \sigma_{I_{amb}}{}^2 + \left(\frac{\partial \rho_{kan}}{\partial [N]_{spk}}\right)^2 \sigma_{[N]_{spk}}{}^2 + \left(\frac{\partial \rho_{kan}}{\partial [N]_{amb}}\right)^2 \sigma_{[N]_{amb}}{}^2 + \left(\frac{\partial \rho_{kan}}{\partial a}\right)^2 \sigma_a{}^2} \tag{B13}$$

The simplest way to differentiate these equations is to rewrite Eq. 4 as:

$$\rho_{kan} = \rho_0 \times g(\rho_0, a, b) \times h(\rho_0, a, b) \tag{B14}$$

where:

$$g(\rho_0, a, b) = (-1 + (1-b)^{1-a}) \tag{B15}$$

$$h(\rho_0, a, b) = \frac{1}{(a-1)b} \tag{B16}$$

Since $\rho_0$ (and hence b) is independent of a (the ratio of nutrient regeneration to "true" uptake rates in the incubation bottle), $\partial\rho_{kan}/\partial a$ can be derived relatively simply by differentiating g and h with respect to a and applying the multiplication rule for derivatives to Eq. A14:

$$\frac{\partial \rho_{kan}}{\partial a} = \frac{\partial \rho_0}{\partial a} g(\rho_0, a, b) h(\rho_0, a, b) + \frac{\partial g}{\partial a} \rho_0 h(\rho_0, a, b) + \frac{\partial h}{\partial a} \rho_0 g(\rho_0, a, b) \tag{B17}$$

$$\frac{\partial \rho_{kan}}{\partial a} = -\ln(1-b)(1-b)^{(1-a)} \times \rho_0 \times h(\rho_0, a, b) + \frac{-\rho_0 \times g(\rho_0, a, b)}{b(a-1)^2} \quad \text{(B18)}$$

$\rho_0$ (and b) are functions of all other variables. Thus for any of these variables (temporarily denoted as x) Eq. A14 must be evaluated as:

$$\frac{\partial \boldsymbol{\rho_{kan}}}{\partial x} = \frac{\partial \rho_0}{\partial x} g(\rho_0, a, b) h(\rho_0, a, b) + \frac{\partial g}{\partial x} \rho_0(x) h(\rho_0, a, b) + \frac{\partial h}{\partial b} \rho_0(x) g(\rho_0, a, b) \quad \text{(B19)}$$

where:

$$\frac{\partial g}{\partial x} = \frac{\partial g}{\partial b} \frac{\partial b}{\partial \rho_0} \frac{\partial \rho_0}{\partial x} \quad \text{(B20)}$$

$$\frac{\partial h}{\partial x} = \frac{\partial h}{\partial b} \frac{\partial b}{\partial \rho_0} \frac{\partial \rho_0}{\partial x} \quad \text{(B21)}$$

for variables, P, $I_{spk}$, $I_{amb}$, $I_P(T)$, and $I_P(0)$, which only appear in b through $\rho_0$. For variables T, $N_{amb}$, and $N_{spk}$, we need to use the equations:

$$\frac{\partial g}{\partial x} = \frac{\partial g}{\partial b} \left( \frac{\partial \left[ \frac{T}{N_{amb} + N_{spk}} \right]}{\partial x} \rho_0 + \frac{T}{N_{amb} + N_{spk}} \frac{\partial \rho_0}{\partial x} \right) \quad \text{(B22)}$$

$$\frac{\partial h}{\partial x} = \frac{\partial h}{\partial b} \left( \frac{\partial \left[ \frac{T}{N_{amb} + N_{spk}} \right]}{\partial x} \rho_0 + \frac{T}{N_{amb} + N_{spk}} \frac{\partial \rho_0}{\partial x} \right) \quad \text{(B23)}$$

We can thus derive:

$$\frac{\partial g}{\partial P} = \frac{a-1}{(1-b)^a} \times \frac{T}{[N]_{amb}+[N]_{spk}} \frac{\partial \rho_0}{\partial P} \tag{B24}$$

$$\frac{\partial g}{\partial T} = 0 \tag{B25}$$

$$\frac{\partial g}{\partial I_P(T)} = \frac{a-1}{(1-b)^a} \times \frac{T}{[N]_{amb}+[N]_{spk}} \frac{\partial \rho_0}{\partial I_P(T)} \tag{B26}$$

$$\frac{\partial g}{\partial I_P(0)} = \frac{a-1}{(1-b)^a} \times \frac{T}{[N]_{amb}+[N]_{spk}} \frac{\partial \rho_0}{\partial I_P(0)} \tag{B27}$$

$$\frac{\partial g}{\partial [N]_{amb}} = \frac{a-1}{(1-b)^a} \left( \frac{-b}{[N]_{amb}+[N]_{spk}} + \frac{T}{[N]_{amb}+[N]_{spk}} \frac{\partial \rho_0}{\partial [N]_{amb}} \right) \tag{B28}$$

$$\frac{\partial g}{\partial [N]_{spk}} = \frac{a-1}{(1-b)^a} \left( \frac{-b}{[N]_{amb}+[N]_{spk}} + \frac{T}{[N]_{amb}+[N]_{spk}} \frac{\partial \rho_0}{\partial [N]_{spk}} \right) \tag{B29}$$

$$\frac{\partial g}{\partial I_{spk}} = \frac{a-1}{(1-b)^a} \times \frac{T}{[N]_{amb}+[N]_{spk}} \frac{\partial \rho_0}{\partial I_{spk}} \tag{B30}$$

$$\frac{\partial g}{\partial I_{amb}} = \frac{a-1}{(1-b)^a} \times \frac{T}{[N]_{amb}+[N]_{spk}} \frac{\partial \rho_0}{\partial I_{amb}} \tag{B31}$$

$$\frac{\partial h}{\partial P} = \frac{-1}{(a-1)b^2} \times \frac{T}{[N]_{amb}+[N]_{spk}} \frac{\partial \rho_0}{\partial P} \tag{B32}$$

$$\frac{\partial h}{\partial T} = 0 \tag{B33}$$

$$\frac{\partial h}{\partial I_P(T)} = \frac{-1}{(a-1)b^2} \times \frac{T}{[N]_{amb}+[N]_{spk}} \frac{\partial \rho_0}{\partial I_P(T)} \tag{B34}$$

$$\frac{\partial h}{\partial I_P(0)} = \frac{-1}{(a-1)b^2} \times \frac{T}{[N]_{amb}+[N]_{spk}} \frac{\partial \rho_0}{\partial I_P(0)} \tag{B35}$$

$$\frac{\partial h}{\partial [N]_{amb}} = \frac{-1}{(a-1)b^2} \times \left(\frac{-b}{[N]_{amb}+[N]_{spk}} + \frac{T}{[N]_{amb}+[N]_{spk}} \frac{\partial \rho_0}{\partial [N]_{amb}}\right) \tag{B36}$$

$$\frac{\partial h}{\partial [N]_{spk}} = \frac{-1}{(a-1)b^2} \times \left(\frac{-b}{[N]_{amb}+[N]_{spk}} + \frac{T}{[N]_{amb}+[N]_{spk}} \frac{\partial \rho_0}{\partial [N]_{spk}}\right) \tag{B37}$$

$$\frac{\partial h}{\partial I_{spk}} = \frac{-1}{(a-1)b^2} \times \frac{T}{[N]_{amb}+[N]_{spk}} \frac{\partial \rho_0}{\partial I_{spk}} \tag{B38}$$

$$\frac{\partial h}{\partial I_{amb}} = \frac{-1}{(a-1)b^2} \times \frac{T}{[N]_{amb}+[N]_{spk}} \frac{\partial \rho_0}{\partial I_{amb}} \tag{B39}$$

Eqs. A20 – A39 and A3 – A10 can thus be inserted into Eq. A15 and A16. These equations can be combined with Eq. A18 and the terms can be inserted into Eq. A13 to quantify the uncertainty in nutrient uptake if isotope dilution is occurring.

#### Supplementary Appendix S2.3 – Uncertainty equations for $\rho_{0,is}$

When the added nutrients from the isotopically-labeled spike are expected to have substantially modified nutrient uptake rates in the incubation bottle relative to nutrient uptake rates *in situ*, the in situ uptake rates can be computed from the incubation uptake rates and knowledge of the half-saturation rate of the ambient phytoplankton community (KS) using the equation:

$$\rho_{0,IS} = \rho_0 \left(\frac{[N]_{amb}}{[N]_{spk}+[N]_{amb}}\right)\left(\frac{[N]_{spk}+[N]_{amb}+K_S}{[N]_{amb}+K_S}\right) \tag{5}$$

Uncertainty in $K_S$ is not expected to be symmetric. It is more realistic to surmise that if $K_S$ is equal to $10^{-1}$ µmol $L^{-1}$, then $K_S$ might range from $10^{-2}$ to $10^{0}$ µmol $L^{-1}$. I therefore replace $K_S$ in Eq. 6 with:

$$K_S = 10^{L10_{KS}} \tag{B40}$$

where $L10_{KS} = \log_{10}(K_S)$. To compute the uncertainty in Eq. 6, I then use the equation:

$$\sigma_{\rho_{0,is}} = \sqrt{\left(\frac{\partial\rho_{0,is}}{\partial P}\right)^2 {\sigma_P}^2 + \left(\frac{\partial\rho_{0,is}}{\partial T}\right)^2 {\sigma_T}^2 + \left(\frac{\partial\rho_{0,is}}{\partial I_P(0)}\right)^2 {\sigma_{I_P(0)}}^2 + \left(\frac{\partial\rho_{0,is}}{\partial I_P(T)}\right)^2 {\sigma_{I_P(T)}}^2 + \left(\frac{\partial\rho_{0,is}}{\partial I_{spk}}\right)^2 {\sigma_{I_{spk}}}^2 + \left(\frac{\partial\rho_{0,is}}{\partial I_{amb}}\right)^2 {\sigma_{I_{amb}}}^2 + \left(\frac{\partial\rho_{0,is}}{\partial [N]_{spk}}\right)^2 {\sigma_{[N]_{spk}}}^2 + \left(\frac{\partial\rho_{0,is}}{\partial [N]_{amb}}\right)^2 {\sigma_{[N]_{amb}}}^2 + \left(\frac{\partial\rho_{0,is}}{\partial L10_{KS}}\right)^2 {\sigma_{L10_{KS}}}^2} \tag{B41}$$

To differentiate Eq. 6, I start by defining:

$$y([N]_{amb}, [N]_{spk}) = \frac{[N]_{amb}}{[N]_{spk}+[N]_{amb}} \tag{B42}$$

$$z([N]_{amb}, [N]_{spk}, K_S) = \frac{[N]_{spk}+[N]_{amb}+K_S}{[N]_{amb}+K_S} \tag{B43}$$

Since y and z are not functions of P, T, $I_P(0)$, $I_P(T)$, $I_{amb}$, and $I_{spk}$:

$$\frac{\partial\rho_{0,is}}{\theta P} = \left(\frac{[N]_{amb}}{[N]_{spk}+[N]_{amb}}\right)\left(\frac{[N]_{spk}+[N]_{amb}+K_S}{[N]_{amb}+K_S}\right)\frac{\partial\rho_0}{\theta P} \tag{B44}$$

$$\frac{\partial\rho_{0,is}}{\theta T} = \left(\frac{[N]_{amb}}{[N]_{spk}+[N]_{amb}}\right)\left(\frac{[N]_{spk}+[N]_{amb}+K_S}{[N]_{amb}+K_S}\right)\frac{\partial\rho_0}{\theta T} \tag{B45}$$

$$\frac{\partial\rho_{0,is}}{\theta I_P(0)} = \left(\frac{[N]_{amb}}{[N]_{spk}+[N]_{amb}}\right)\left(\frac{[N]_{spk}+[N]_{amb}+K_S}{[N]_{amb}+K_S}\right)\frac{\partial\rho_0}{\theta I_P(0)} \tag{B46}$$

$$\frac{\partial\rho_{0,is}}{\theta I_P(T)} = \left(\frac{[N]_{amb}}{[N]_{spk}+[N]_{amb}}\right)\left(\frac{[N]_{spk}+[N]_{amb}+K_S}{[N]_{amb}+K_S}\right)\frac{\partial\rho_0}{\theta I_P(T)} \tag{B47}$$

$$\frac{\partial \rho_{0,is}}{\theta I_{spk}} = \left(\frac{[N]_{amb}}{[N]_{spk}+[N]_{amb}}\right)\left(\frac{[N]_{spk}+[N]_{amb}+K_S}{[N]_{amb}+K_S}\right)\frac{\partial \rho_0}{\theta I_{spk}} \tag{B48}$$

$$\frac{\partial \rho_{0,is}}{\theta I_{amb}} = \left(\frac{[N]_{amb}}{[N]_{spk}+[N]_{amb}}\right)\left(\frac{[N]_{spk}+[N]_{amb}+K_S}{[N]_{amb}+K_S}\right)\frac{\partial \rho_0}{\theta I_{amb}} \tag{B49}$$

Since, $\rho_0$ and $y([N]_{amb},[N]_{spk})$ are not functions of $K_S$:

$$\frac{\partial \rho_{0,is}}{\partial L10_{KS}} = \rho_0\left(\frac{[N]_{amb}}{[N]_{spk}+[N]_{amb}}\right)\frac{\partial z}{\partial L10_{KS}} \tag{B50}$$

$$\frac{\partial \rho_{0,is}}{\partial L10_{KS}} = \rho_0\left(\frac{[N]_{amb}}{[N]_{spk}+[N]_{amb}}\right)\left(\frac{-\ln(10)\times[N]_{spk}\times K_S}{([N]_{amb}+K_S)^2}\right) \tag{B51}$$

The derivative of Eq. 6 with respect to $N_{amb}$ and $N_{spk}$ can be found as:

$$\frac{\partial \rho_{0,IS}}{\partial N_{amb}} = \frac{\partial \rho_0}{\partial N_{amb}}\times y\times z + \frac{\partial y}{\partial N_{amb}}\times \rho_0\times z + \frac{\partial z}{\partial N_{amb}}\times \rho_0\times y \tag{B52}$$

$$\frac{\partial \rho_{0,IS}}{\partial N_{spk}} = \frac{\partial \rho_0}{\partial N_{spk}}\times y\times z + \frac{\partial y}{\partial N_{spk}}\times \rho_0\times z + \frac{\partial z}{\partial N_{spk}}\times \rho_0\times y \tag{B53}$$

where:

$$\frac{\partial y}{\partial N_{amb}} = \frac{[N]_{spk}}{\left([N]_{spk}+[N]_{amb}\right)^2} \tag{B54}$$

$$\frac{\partial z}{\partial N_{amb}} = \frac{-[N]_{spk}}{(K_S+[N]_{amb})^2} \tag{B55}$$

$$\frac{\partial y}{\partial N_{spk}} = \frac{-[N]_{amb}}{\left([N]_{spk}+[N]_{amb}\right)^2} \tag{B56}$$

$$\frac{\partial z}{\partial N_{spk}} = \frac{1}{K_S+[N]_{amb}} \tag{B57}$$

#### Supplementary Appendix S2.4 – Uncertainty equations for $\rho_{kan,is}$

When isotope dilution and modified nutrient uptake rates resulting from the added tracer spike are both suspected to be quantitatively important, nutrient uptake should be computed from Eq. 7:

$$\rho_{kan,IS} = \rho_{kan}\left(\frac{[N]_{amb}}{[N]_{spk}+[N]_{amb}}\right)\left(\frac{[N]_{spk}+[N]_{amb}+K_S}{[N]_{amb}+K_S}\right) \tag{6}$$

Uncertainty in Eq. 7 should be quantified using the following equation:

$$\sigma_{\rho_{kan,is}} = \sqrt{\left(\frac{\partial\rho_{kan,is}}{\partial P}\right)^2 \sigma_P{}^2 + \left(\frac{\partial\rho_{kan,is}}{\partial T}\right)^2 \sigma_T{}^2 + \left(\frac{\partial\rho_{kan,is}}{\partial I_P(0)}\right)^2 \sigma_{I_P(0)}{}^2 + \left(\frac{\partial\rho_{kan,is}}{\partial I_P(T)}\right)^2 \sigma_{I_P(T)}{}^2 + \left(\frac{\partial\rho_{kan,is}}{\partial I_{spk}}\right)^2 \sigma_{I_{spk}}{}^2 + \left(\frac{\partial\rho_{kan,is}}{\partial I_{amb}}\right)^2 \sigma_{I_{amb}}{}^2 + \left(\frac{\partial\rho_{kan,is}}{\partial [N]_{spk}}\right)^2 \sigma_{[N]_{spk}}{}^2 + \left(\frac{\partial\rho_{kan,is}}{\partial [N]_{amb}}\right)^2 \sigma_{[N]_{amb}}{}^2 + \left(\frac{\partial\rho_{kan,is}}{\partial a}\right)^2 \sigma_{K_S}{}^2 + \left(\frac{\partial\rho_{kan,is}}{\partial L10_{KS}}\right)^2 \sigma_{L10_{KS}}{}^2} \tag{B58}$$

Using derivations nearly identical to those for Eq. 6, it can be shown that:

$$\frac{\partial\rho_{kan,is}}{\partial P} = \left(\frac{[N]_{amb}}{[N]_{spk}+[N]_{amb}}\right)\left(\frac{[N]_{spk}+[N]_{amb}+K_S}{[N]_{amb}+K_S}\right)\frac{\partial\rho_{kan}}{\partial P} \tag{B59}$$

$$\frac{\partial\rho_{kan,is}}{\partial T} = \left(\frac{[N]_{amb}}{[N]_{spk}+[N]_{amb}}\right)\left(\frac{[N]_{spk}+[N]_{amb}+K_S}{[N]_{amb}+K_S}\right)\frac{\partial\rho_{kan}}{\partial T} \tag{B60}$$

$$\frac{\partial\rho_{kan,is}}{\partial I_P(0)} = \left(\frac{[N]_{amb}}{[N]_{spk}+[N]_{amb}}\right)\left(\frac{[N]_{spk}+[N]_{amb}+K_S}{[N]_{amb}+K_S}\right)\frac{\partial\rho_{kan}}{\partial I_P(0)} \tag{B61}$$

$$\frac{\partial\rho_{kan,is}}{\partial I_P(T)} = \left(\frac{[N]_{amb}}{[N]_{spk}+[N]_{amb}}\right)\left(\frac{[N]_{spk}+[N]_{amb}+K_S}{[N]_{amb}+K_S}\right)\frac{\partial\rho_{kan}}{\partial I_P(T)} \tag{B62}$$

$$\frac{\partial\rho_{kan,is}}{\partial I_{spk}} = \left(\frac{[N]_{amb}}{[N]_{spk}+[N]_{amb}}\right)\left(\frac{[N]_{spk}+[N]_{amb}+K_S}{[N]_{amb}+K_S}\right)\frac{\partial\rho_{kan}}{\partial I_{spk}} \tag{B63}$$

$$\frac{\partial \rho_{kan,is}}{\partial I_{amb}} = \left(\frac{[N]_{amb}}{[N]_{spk}+[N]_{amb}}\right)\left(\frac{[N]_{spk}+[N]_{amb}+K_S}{[N]_{amb}+K_S}\right)\frac{\partial \rho_{kan}}{\partial I_{amb}} \tag{B64}$$

$$\frac{\partial \rho_{kan,is}}{\partial a} = \left(\frac{[N]_{amb}}{[N]_{spk}+[N]_{amb}}\right)\left(\frac{[N]_{spk}+[N]_{amb}+K_S}{[N]_{amb}+K_S}\right)\frac{\partial \rho_{kan}}{\partial a} \tag{B65}$$

$$\frac{\partial \rho_{kan,is}}{\partial L10_{KS}} = \rho_{kan}\left(\frac{[N]_{amb}}{[N]_{spk}+[N]_{amb}}\right)\left(\frac{-\ln(10)\times[N]_{spk}\times K_S}{([N]_{amb}+K_S)^2}\right) \tag{B66}$$

$$\frac{\partial \rho_{kan,IS}}{\partial N_{amb}} = \frac{\partial \rho_{kan}}{\partial N_{amb}} \times y \times z + \frac{\partial y}{\partial N_{amb}} \times \rho_{kan} \times z + \frac{\partial z}{\partial N_{amb}} \times \rho_{kan} \times y \tag{B67}$$

$$\frac{\partial \rho_{kan,IS}}{\partial N_{spk}} = \frac{\partial \rho_{kan}}{\partial N_{spk}} \times y \times z + \frac{\partial y}{\partial N_{spk}} \times \rho_{kan} \times z + \frac{\partial z}{\partial N_{spk}} \times \rho_{kan} \times y \tag{B68}$$

and:

$$\frac{\partial y}{\partial N_{amb}} = \frac{[N]_{spk}}{\left([N]_{spk}+[N]_{amb}\right)^2} \tag{B69}$$

$$\frac{\partial z}{\partial N_{amb}} = \frac{-[N]_{spk}}{(K_S+[N]_{amb})^2} \tag{B70}$$

$$\frac{\partial y}{\partial N_{spk}} = \frac{-[N]_{amb}}{\left([N]_{spk}+[N]_{amb}\right)^2} \tag{B71}$$

$$\frac{\partial z}{\partial N_{spk}} = \frac{1}{K_S+[N]_{amb}} \tag{B72}$$

#### Supplementary Appendix S2.5 – Uncertainty equations for $\rho_{reg}$ (nutrient uptake with regeneration and temporally-varying isotope dilution)

When substantial nutrient regeneration is occurring, $\rho_{kan}$ may not be appropriate because it assumes that all of the regenerated nutrients will have isotopic ratios equal to the isotopic ratio of natural POM (i.e., it assumes that labeled nitrogen taken up during the experiment cannot be recycled) and it also assumes that $\Delta I_P(t)$ is constant in time. If we relax these assumptions, but instead assume that the PON concentration and substrate concentration are constant in time (which will be true if nutrient regeneration is complete), we can quantify nutrient uptake as:

$$\rho_{reg} = \left( ln\left(\frac{I_S(0)-a\times I_P(0)}{P}\right) - ln\left(\frac{I_P(0)-I_P(t)}{[N_{spk}+N_{amb}]} + \frac{I_S(0)-a\times I_P(t)}{P}\right)\right)\left(\frac{P\times[N_{spk}+N_{amb}]}{P+a\times[N_{spk}+N_{amb}]}\right)\frac{1}{T} \quad (13)$$

Uncertainty in Eq. 13 should be quantified using the equation:

$$\sigma_{\rho_{reg}} = \sqrt{\left(\frac{\partial\rho_{reg}}{\partial P}\right)^2 {\sigma_P}^2 + \left(\frac{\partial\rho_{reg}}{\partial T}\right)^2 {\sigma_T}^2 + \left(\frac{\partial\rho_{reg}}{\partial I_P(0)}\right)^2 {\sigma_{I_P(0)}}^2 + \left(\frac{\partial\rho_{reg}}{\partial I_P(T)}\right)^2 {\sigma_{I_P(T)}}^2 + \left(\frac{\partial\rho_{reg}}{\partial I_{spk}}\right)^2 {\sigma_{I_{spk}}}^2 + \left(\frac{\partial\rho_{reg}}{\partial I_{amb}}\right)^2 {\sigma_{I_{amb}}}^2 + \left(\frac{\partial\rho_{reg}}{\partial [N]_{spk}}\right)^2 {\sigma_{[N]_{spk}}}^2 + \left(\frac{\partial\rho_{reg}}{\partial [N]_{amb}}\right)^2 {\sigma_{[N]_{amb}}}^2 + \left(\frac{\partial\rho_{reg}}{\partial a}\right)^2 {\sigma_a}^2} \quad (B73)$$

where:

$$\frac{\partial\rho_{reg}}{\partial P} = \left(\frac{1}{\left(\frac{I_P(0)-I_P(t)}{[N_{spk}+N_{amb}]}\right)P+\left(I_S(0)-a\times I_P(t)\right)}\right)\left(\frac{I_P(t)-I_P(0)}{P+a\times[N_{spk}+N_{amb}]}\right)\frac{1}{T} + \frac{\rho_{reg}}{P}\left(\frac{[N_{spk}+N_{amb}]}{P+a[N_{spk}+N_{amb}]}\right) \quad (B74)$$

$$\frac{\partial\rho_{reg}}{\partial T} = \left(\frac{-1}{T}\right)\rho_{reg} \quad (B75)$$

$$\frac{\partial\rho_{reg}}{\partial I_P(0)} = \left(\frac{a}{a\times I_P(0)-I_S(0)} - \frac{P}{\left(I_P(0)-I_P(t)\right)\times P+\left([N_{spk}+N_{amb}]\right)\left(I_S(0)-a\times I_P(t)\right)}\right)\left(\frac{P\times[N_{spk}+N_{amb}]}{P+a\times[N_{spk}+N_{amb}]}\right)\frac{1}{T} \quad (B76)$$

$$\frac{\partial\rho_{reg}}{\partial I_P(T)} = \frac{-P\times[N_{spk}+N_{amb}]}{\left(P+a[N_{spk}+N_{amb}]\right)I_P(t)-[N_{spk}+N_{amb}]I_S(0)-I_P(0)P}\times\frac{1}{T}$$

$$\frac{\partial\rho_{reg}}{\partial I_{spk}} = \left(\frac{1}{I_S(0)-a\times I_P(0)} - \frac{[N_{spk}+N_{amb}]}{\left(I_S(0)-a\times I_P(t)\right)\left([N]_{spk}+[N]_{amb}\right)+\left(I_P(0)-I_P(t)\right)P}\right)\left(\frac{P\times[N]_{spk}}{P+a\times[N_{spk}+N_{amb}]}\right)\frac{1}{T} \quad (B77)$$

$$\frac{\partial\rho_{reg}}{\partial I_{amb}} = \left(\frac{1}{I_S(0)-a\times I_P(0)} - \frac{[N_{spk}+N_{amb}]}{\left(I_S(0)-a\times I_P(t)\right)[N_{spk}+N_{amb}]+P\left(I_P(0)-I_P(t)\right)}\right)\left(\frac{P\times[N]_{amb}}{P+a\times[N_{spk}+N_{amb}]}\right)\frac{1}{T} \quad (B78)$$

For calculating derivatives with respect to $N_{spk}$ and $N_{amb}$, I define:

$$f = ln\left(\frac{I_S(0) - a \times I_P(0)}{P}\right) \tag{B79}$$

$$g = ln\left(\frac{I_P(0) - I_P(t)}{[N_{spk} + N_{amb}]} + \frac{I_S(0) - a \times I_P(t)}{P}\right) \tag{B80}$$

$$h = \left(\frac{P \times [N_{spk} + N_{amb}]}{P + a \times [N_{spk} + N_{amb}]}\right) \tag{B81}$$

Therefore:

$$\rho_{reg} = (f - g)h\frac{1}{T} \tag{B82}$$

$$\frac{\partial \rho_{reg}}{\partial N_{spk}} = \left(\frac{\partial f}{\partial N_{spk}} - \frac{\partial g}{\partial N_{spk}}\right)h\frac{1}{T} + (f - g)\frac{1}{T}\frac{\partial h}{\partial N_{spk}} \tag{B83}$$

$$\frac{\partial f}{\partial [N]_{spk}} = \frac{\left(I_{amb} - I_{spk}\right)[N]_{amb}}{\left([N]_{spk} + [N]_{amb}\right)\left(\left(aI_P(0) - I_{spk}\right)[N]_{spk} + \left(aI_P(0) - I_{amb}\right)[N]_{amb}\right)} \tag{B84}$$

$$\frac{\partial g}{\partial [N]_{spk}} = -\frac{PI_P(t) + \left(I_{spk} - I_{amb}\right)[N]_{amb} - PI_P(0)}{\left([N]_{spk} + [N]_{amb}\right)\left(\left(aI_P(t) - I_{spk}\right)[N]_{spk} + \left(a[N]_{amb} + P\right)I_P(t) - I_{amb}[N]_{amb} - PI_P(0)\right)} \tag{B85}$$

$$\frac{\partial h}{\partial [N]_{spk}} = \frac{P^2}{\left(a[N]_{spk} + a[N]_{amb} + P\right)^2} \tag{B86}$$

$$\frac{\partial \rho_{reg}}{\partial N_{amb}} = \left(\frac{\partial f}{\partial N_{amb}} - \frac{\partial g}{\partial N_{amb}}\right)h\frac{1}{T} + (f - g)\frac{1}{T}\frac{\partial h}{\partial N_{amb}} \tag{B87}$$

$$\frac{\partial f}{\partial [N]_{amb}} = \frac{-\left(I_{amb} - I_{spk}\right)[N]_{spk}}{\left([N]_{spk} + [N]_{amb}\right)\left(\left(aI_P(0) - I_{spk}\right)[N]_{spk} + \left(aI_P(0) - I_{amb}\right)[N]_{amb}\right)} \tag{B88}$$

$$\frac{\partial g}{\partial [N]_{amb}} = -\frac{P \times I_P(t) + \left(I_{amb} - I_{spk}\right)[N]_{spk} - P \times I_P(0)}{\left([N]_{spk} + [N]_{amb}\right)\left(\left(a \times I_P(t) - I_{amb}\right)[N]_{amb} + \left(a[N]_{spk} + P\right)I_P(t) - I_{spk}[N]_{spk} - P \times I_P(0)\right)} \tag{B89}$$

$$\frac{\partial h}{\partial [N]_{amb}} = \frac{P^2}{\left(a[N]_{spk}+a[N]_{amb}+P\right)^2} \tag{B90}$$

$$\frac{\partial \rho_{reg}}{\partial a} = \left(\frac{I_P(0)}{a\times I_P(0)-I_S(0)} - \frac{I_P(T)\times[N_{spk}+N_{amb}]}{\left(a\times I_P(T)-I_S(0)\right)[N_{spk}+N_{amb}]-P\left(I_P(0)-I_P(t)\right)}\right)\left(\frac{P\times[N_{spk}+N_{amb}]}{P+a\times[N_{spk}+N_{amb}]}\right)\frac{1}{T} + \left(ln\left(\frac{I_S(0)-a\times I_P(0)}{P}\right) - ln\left(\frac{I_P(0)-I_P(t)}{[N_{spk}+N_{amb}]} + \frac{I_S(0)-a\times I_P(t)}{P}\right)\right)\frac{-P\left([N_{spk}+N_{amb}]\right)^2}{\left(\left([N_{spk}+N_{amb}]\right)a+P\right)^2}\frac{1}{T} \tag{B91}$$

#### Supplementary Appendix S2.6 – Uncertainty equations for $\rho_{reg,is}$

When substantial isotope regeneration and modified nutrient uptake rates resulting from the added tracer spike are both suspected to be quantitatively important, nutrient uptake should be computed from Eq. 14:

$$\rho_{reg,IS} = \rho_{reg}\left(\frac{[N]_{amb}}{[N]_{spk}+[N]_{amb}}\right)\left(\frac{[N]_{spk}+[N]_{amb}+K_S}{[N]_{amb}+K_S}\right) \tag{14}$$

Uncertainty in Eq. 14 should be quantified using the following equation:

$$\sigma_{\rho_{reg,is}} = \sqrt{\left(\frac{\partial\rho_{reg,is}}{\partial P}\right)^2 {\sigma_P}^2 + \left(\frac{\partial\rho_{reg,is}}{\partial T}\right)^2 {\sigma_T}^2 + \left(\frac{\partial\rho_{reg,is}}{\partial I_P(0)}\right)^2 {\sigma_{I_P(0)}}^2 + \left(\frac{\partial\rho_{reg,is}}{\partial I_P(T)}\right)^2 {\sigma_{I_P(T)}}^2 + \left(\frac{\partial\rho_{reg,is}}{\partial I_{spk}}\right)^2 {\sigma_{I_{spk}}}^2 + \left(\frac{\partial\rho_{reg,is}}{\partial I_{amb}}\right)^2 {\sigma_{I_{amb}}}^2 + \left(\frac{\partial\rho_{reg,is}}{\partial [N]_{spk}}\right)^2 {\sigma_{[N]_{spk}}}^2 + \left(\frac{\partial\rho_{reg,is}}{\partial [N]_{amb}}\right)^2 {\sigma_{[N]_{amb}}}^2 + \left(\frac{\partial\rho_{reg,is}}{\partial a}\right)^2 {\sigma_{K_S}}^2 + \left(\frac{\partial\rho_{reg,is}}{\partial L10_{KS}}\right)^2 {\sigma_{L10_{KS}}}^2} \tag{B92}$$

Using derivations nearly identical to those for Eq. 6, it can be shown that:

$$\frac{\partial \rho_{reg,is}}{\partial P} = \left(\frac{[N]_{amb}}{[N]_{spk}+[N]_{amb}}\right)\left(\frac{[N]_{spk}+[N]_{amb}+K_S}{[N]_{amb}+K_S}\right)\frac{\partial \rho_{reg}}{\partial P} \tag{B93}$$

$$\frac{\partial \rho_{reg,is}}{\partial T} = \left(\frac{[N]_{amb}}{[N]_{spk}+[N]_{amb}}\right)\left(\frac{[N]_{spk}+[N]_{amb}+K_S}{[N]_{amb}+K_S}\right)\frac{\partial \rho_{reg}}{\partial T} \tag{B94}$$

$$\frac{\partial \rho_{reg,is}}{\partial I_P(0)} = \left(\frac{[N]_{amb}}{[N]_{spk}+[N]_{amb}}\right)\left(\frac{[N]_{spk}+[N]_{amb}+K_S}{[N]_{amb}+K_S}\right)\frac{\partial \rho_{reg}}{\partial I_P(0)} \tag{B95}$$

$$\frac{\partial \rho_{reg,is}}{\partial I_P(T)} = \left(\frac{[N]_{amb}}{[N]_{spk}+[N]_{amb}}\right)\left(\frac{[N]_{spk}+[N]_{amb}+K_S}{[N]_{amb}+K_S}\right)\frac{\partial \rho_{reg}}{\partial I_P(T)} \tag{B96}$$

$$\frac{\partial \rho_{reg,is}}{\partial I_{spk}} = \left(\frac{[N]_{amb}}{[N]_{spk}+[N]_{amb}}\right)\left(\frac{[N]_{spk}+[N]_{amb}+K_S}{[N]_{amb}+K_S}\right)\frac{\partial \rho_{reg}}{\partial I_{spk}} \tag{B97}$$

$$\frac{\partial \rho_{reg,is}}{\partial I_{amb}} = \left(\frac{[N]_{amb}}{[N]_{spk}+[N]_{amb}}\right)\left(\frac{[N]_{spk}+[N]_{amb}+K_S}{[N]_{amb}+K_S}\right)\frac{\partial \rho_{reg}}{\partial I_{amb}} \tag{B98}$$

$$\frac{\partial \rho_{reg,is}}{\partial a} = \left(\frac{[N]_{amb}}{[N]_{spk}+[N]_{amb}}\right)\left(\frac{[N]_{spk}+[N]_{amb}+K_S}{[N]_{amb}+K_S}\right)\frac{\partial \rho_{reg}}{\partial a} \tag{B99}$$

$$\frac{\partial \rho_{reg,is}}{\partial L10_{KS}} = \rho_{reg}\left(\frac{[N]_{amb}}{[N]_{spk}+[N]_{amb}}\right)\left(\frac{-\ln(10)\times[N]_{spk}\times K_S}{([N]_{amb}+K_S)^2}\right) \tag{B100}$$

$$\frac{\partial \rho_{reg,IS}}{\partial N_{amb}} = \frac{\partial \rho_{reg}}{\partial N_{amb}} \times y \times z + \frac{\partial y}{\partial N_{amb}} \times \rho_{reg} \times z + \frac{\partial z}{\partial N_{amb}} \times \rho_{reg} \times y \tag{B101}$$

$$\frac{\partial \rho_{reg,IS}}{\partial N_{spk}} = \frac{\partial \rho_{reg}}{\partial N_{spk}} \times y \times z + \frac{\partial y}{\partial N_{spk}} \times \rho_{reg} \times z + \frac{\partial z}{\partial N_{spk}} \times \rho_{reg} \times y \tag{B102}$$

and:

$$\frac{\partial y}{\partial N_{amb}} = \frac{[N]_{spk}}{\left([N]_{spk}+[N]_{amb}\right)^2} \tag{B103}$$

$$\frac{\partial z}{\partial N_{amb}} = \frac{-[N]_{spk}}{(K_S+[N]_{amb})^2} \tag{B104}$$

$$\frac{\partial y}{\partial N_{spk}} = \frac{-[N]_{amb}}{\left([N]_{spk}+[N]_{amb}\right)^2} \tag{B105}$$

$$\frac{\partial z}{\partial N_{spk}} = \frac{1}{K_S+[N]_{amb}} \tag{B106}$$

#### Supplementary Appendix S2.7 – Propagation of uncertainty with paired measurements

Most frequently scientists compute uncertainty in $\rho_0$ by conducting incubations in duplicate or triplicate and then computing the standard deviation (SD) or standard error of the mean (SE = SD/sqrt($N_{inc}$)) of the paired measurements where $N_{inc}$ is the number of incubations conducted. They then use this standard deviation as the uncertainty in the measurements. This approach is inaccurate, however, if the error in any of the input arguments (T, P, $I_P(T)$, $I_P(0)$, $I_{spk}$, $I_{amb}$, $[N]_{spk}$, $[N]_{amb}$ is expected to be correlated. Correlated errors are highly likely to occur in situations where a single value is used for each incubation. For instance, it is rare for the ambient nutrient concentration ($[N]_{amb}$) to be measured independently in each incubation bottle. Instead, it is commonly measured on a separate sample drawn from the same environmental sampling bottle. If this single measurement of $[N]_{amb}$ is applied to each incubation, the resultant nutrient uptake rates ($\rho_0$) computed from Eq. 1 (or Eq. A2) should not be considered independent. By contrast, in a typical set of triplicate uptake incubations, P an d $I_P(T)$ are measured at the end in each experimental bottle. These variables can thus be considered independent. To accurately quantify uncertainty in such incubations when SD or SE are used instead of the uncertainty in each individual parameter, I begin by defining $X_1, \ldots, X_{Nmeas}$ as the variables that are measured independently for each incubation and $Y_1, \ldots, Y_{Nassume}$ as the variables that are not measured independently for each incubation bottle, where $N_{meas}$ is the number of variables independently measured in each incubation bottle and $N_{assume}$ is the number of variables that are assumed to be identical in all incubation bottles. The goal of most incubation experiments is to estimate the true nutrient uptake rate *in situ* ($\rho$). However, in practice, we instead find the arithmetic mean of nutrient uptake in several incubation bottles, using Eqs. 1, 4, 6, or 7. For instance:

$$\overline{\rho_0} = \frac{1}{N_{inc}} \sum_{i=1}^{N_{inc}} \rho_{0,i} \tag{B107}$$

where $N_{inc}$ is the number of incubations conducted and $\rho_{0,i}$ is the computed uptake rate in the $i^{th}$ incubation. Since $\rho_{0,i}$ is a function of $X_{1,i}, \ldots X_{Nmease,i}$ and $Y_1, \ldots, Y_{Nmeas}$, it follows that $\overline{\rho_0}$ must also be a function of X and Y. However, when constant values are assumed for $Y_1, \ldots, Y_{Nmeas}$, the sample standard deviation (SD) and sample standard error of the mean (SE = SD/sqrt($N_{inc}$)) will depend on $\sigma_X$, but not $\sigma_Y$. Using constant values for Y for calculating $\rho_{0,i}$ is equivalent to assuming $\sigma_Y = 0$. I will thus define:

$$\sigma'_{\overline{\rho_0}} = \frac{1}{N_{inc}} \sqrt{\sum_i^{N_{inc}} \sum_k^{N_{meas}} \left(\frac{\partial \rho_{0,i}}{\partial X_{i,k}}\right)^2 {\sigma_{X_{i,k}}}^2} \qquad \text{(B108)}$$

which is the uncertainty in Eq. A73 if all variables Y have zero uncertainty and measurement error in all parameters X is assumed to be uncorrelated. Since these are the same assumptions inherent to calculating the sample standard deviation (SD) or sample standard error of the mean (SE), it follows that SE = $\sigma'_{\overline{\rho_0}}$. However, since both SE and $\sigma'_{\overline{\rho_0}}$ neglect uncertainty in Y, it is clear that they will be biased estimators for the true uncertainty in ρ. Instead, the true uncertainty in ρ can be calculated as:

$$\sigma_{\overline{\rho_0}} = \frac{1}{N_{inc}} \sqrt{\sum_i^{N_{inc}} \sum_k^{N_{meas}} \left(\frac{\partial \rho_{0,i}}{\partial X_{i,k}}\right)^2 {\sigma_{X_{i,k}}}^2 + \sum_i^{N_{inc}} \sum_k^{N_{meas}} \left(\frac{\partial \rho_{0,i}}{\partial Y_{i,k}}\right)^2 {\sigma_{Y_{i,k}}}^2 + \sum_i^{N_{inc}} \sum_{j(j\neq 1)}^{N_{inc}} \sum_{k=1}^{N_{meas}} \left(\frac{\partial \rho_{0,i}}{\partial Y_{i,k}}\right) \left(\frac{\partial \rho_{0,j}}{\partial Y_{j,k}}\right) \sigma_{Y_{i,k}, Y_{j,k}}} \qquad \text{(B109)}$$

Since $Y_{i,k} = Y_{j,k}$, the covariance ($\sigma_{Y_{i,k}, Y_{j,k}}$) is equal to ${\sigma_{Y_{i,k}}}^2$. If I make the simplifying assumption that:

$$\left(\frac{\partial \rho_{0,i}}{\partial Y_{i,k}}\right) = \left(\frac{\partial \rho_{0,j}}{\partial Y_{j,k}}\right) \qquad \text{(B110)}$$

I can simplify Eq. A75 to:

$$\sigma_{\overline{\rho_0}} \approx \frac{1}{N_{inc}} \sqrt{\sum_i^{N_{inc}} \sum_k^{N_{meas}} \left(\frac{\partial \rho_{0,i}}{\partial X_{i,k}}\right)^2 {\sigma_{X_{i,k}}}^2 + {N_{inc}}^2 \sum_k^{N_{meas}} \left(\frac{\partial \rho_{0,i}}{\partial Y_{i,k}}\right)^2 {\sigma_{Y_{i,k}}}^2} \qquad \text{(B111)}$$

and if I substitute in Eq. A74, remembering that SE = $\sigma'_{\overline{\rho_0}}$, I get an estimate for the uncertainty resulting from combining the measured standard error with uncertainty in the variables $Y_1, \ldots Y_{Nassume}$ that were applied to all incubations:

$$\sigma_{\overline{\rho_0}} \approx \sqrt{SE^2 + \sum_k^{N_{meas}} \left(\frac{\partial \rho_{0,i}}{\partial Y_{i,k}}\right)^2 \sigma_{Y_{i,k}}{}^2} \tag{B112}$$

I will assume that uncertainty in $[N]_{spk}$, arises from variability from one spike to another, rather than from inaccurate calibration of the pipet that affects all spikes. In such a case $[N]_{spk}$ is also independent across the different incubations (note that although in reality errors in $[N]_{spk}$ should be considered weakly correlated, $\sigma_{Nspk}$ is only a minor contributor to $\sigma_{\rho 0}$, so this assumption introduces little error to the final estimate). $I_P(0)$, $I_{spk}$, and $I_{amb}$ are usually assumed to take the same value in all incubations and hence cannot be considered independent. However, $\sigma_{IP(0)}$, $\sigma_{Ispk}$, and $\sigma_{Iamb}$ are all minor contributors to $\sigma_{\rho 0}$, so I will neglect them here. That leaves $[N]_{amb}$ and T as the correlated variables most likely to impact our estimates of *in situ* nutrient uptake measured using duplicate or triplicate incubations. Although it might seem that T should be uncorrelated between experiments (since incubation start time, filtration start time, and filtration end time can be measured independently) the greatest uncertainty in T actually arises from uncertainty in what time should be used for the termination of the incubation (e.g., beginning of filtration, midpoint of filtration, or end of filtration). In the example I have illustrated here, uncertainty in ρ0 can be calculated as:

$$\sigma_{\overline{\rho_0}} \approx \sqrt{SE_{\rho_0}{}^2 + \left(\frac{\partial \rho_0}{\partial T}\right)^2 \sigma_T{}^2 + \left(\frac{\partial \rho_0}{\partial N_{amb}}\right)^2 \sigma_{N_{amb}}{}^2} \tag{B113}$$

Where $SE_{\rho 0}$ is the sample standard error calculated from multiple replicate incubations using Eq. 1 and a constant value for $N_{amb}$ and T in all incubations and $\partial\rho_0/\partial T$ and $\partial\rho_0/\partial N_{amb}$ are given by Eqs. B4 and B8. Using the same arguments as advanced above, it is easy to show that when correcting for isotope dilution or increased uptake in the incubation bottle relative to in situ using Eqs. 3, 13, 5, 6, or 14 respectively, uncertainty can be calculated as:

$$\sigma_{\overline{\rho_{kan}}} \approx \sqrt{SE_{\rho_{kan}}{}^2 + \left(\frac{\partial \rho_{kan}}{\partial T}\right)^2 \sigma_T{}^2 + \left(\frac{\partial \rho_{kan}}{\partial N_{amb}}\right)^2 \sigma_{N_{amb}}{}^2 + \left(\frac{\partial \rho_{kan}}{\partial a}\right)^2 \sigma_a{}^2} \tag{B114}$$

$$\sigma_{\overline{\rho_{reg}}} \approx \sqrt{SE_{\rho_{reg}}{}^2 + \left(\frac{\partial \rho_{reg}}{\partial T}\right)^2 \sigma_T{}^2 + \left(\frac{\partial \rho_{reg}}{\partial N_{amb}}\right)^2 \sigma_{N_{amb}}{}^2 + \left(\frac{\partial \rho_{reg}}{\partial a}\right)^2 \sigma_a{}^2} \tag{B115}$$

$$\sigma_{\overline{\rho_{0,is}}} \approx \sqrt{SE_{\rho_{0,is}}{}^2 + \left(\frac{\partial \rho_{0,is}}{\partial T}\right)^2 \sigma_T{}^2 + \left(\frac{\partial \rho_{0,is}}{\partial N_{amb}}\right)^2 \sigma_{N_{amb}}{}^2 + \left(\frac{\partial \rho_{0,is}}{\partial L10_{KS}}\right)^2 \sigma_{L10_{KS}}{}^2} \tag{B116}$$

$$\sigma_{\overline{\rho_{kan,is}}} \approx \sqrt{SE_{\rho_{kan,is}}{}^2 + \left(\frac{\partial \rho_{kan,is}}{\partial T}\right)^2 \sigma_T{}^2 + \left(\frac{\partial \rho_{kan,is}}{\partial N_{amb}}\right)^2 \sigma_{N_{amb}}{}^2 + \left(\frac{\partial \rho_{kan,is}}{\partial a}\right)^2 \sigma_a{}^2 + \left(\frac{\partial \rho_{kan,is}}{\partial L10_{KS}}\right)^2 \sigma_{L10_{KS}}{}^2} \tag{B117}$$

$$\sigma_{\overline{\rho_{reg,is}}} \approx \sqrt{SE_{\rho_{reg,is}}{}^2 + \left(\frac{\partial \rho_{reg,is}}{\partial T}\right)^2 \sigma_T{}^2 + \left(\frac{\partial \rho_{reg,is}}{\partial N_{amb}}\right)^2 \sigma_{N_{amb}}{}^2 + \left(\frac{\partial \rho_{reg,is}}{\partial a}\right)^2 \sigma_a{}^2 + \left(\frac{\partial \rho_{reg,is}}{\partial L10_{KS}}\right)^2 \sigma_{L10_{KS}}{}^2} \quad \text{(B118)}$$

where $\partial\rho_{kan}/\partial T$, $\partial\rho_{kan}/\partial N_{amb}$, and $\partial\rho_{kan}/\partial a$ can be calculated from Eqs. B17, B19, B20, B21, B28, and B36; $\partial\rho_{reg}/\partial T$, $\partial\rho_{reg}/\partial N_{amb}$, and $\partial\rho_{reg}/\partial a$ can be calculated from Eqs. B75, B87, B91; $\partial\rho_{0,is}/\partial T$, $\partial\rho_{0,is}/\partial N_{amb}$, and $\partial\rho_{0,is}/\partial L10_{KS}$ can be calculated from Eqs. B45, B51, and B52; $\partial\rho_{kan,is}/\partial T$, $\partial\rho_{kan,is}/\partial N_{amb}$, $\partial\rho_{kan,is}/\partial L10_{KS}$, and $\partial\rho_{kan,is}/\partial a$ can be calculated from Eqs. B60, B65, B66, and B67; and $\partial\rho_{reg,is}/\partial T$, $\partial\rho_{reg,is}/\partial N_{amb}$, $\partial\rho_{reg,is}/\partial L10_{KS}$, and $\partial\rho_{reg,is}/\partial a$ can be calculated from Eqs. B94, B101, B100, and B99.
