## Supplementary material for "Investigating equations for measuring dissolved inorganic nutrient uptake in oligotrophic conditions": Calculation Files for Nutrient Uptake Eqs: 1 - File List and Explanations - Read Me.pdf

This folder contains a series of files that can be used to calculate nutrient uptake rates ($\rho_0$, $\rho_{kan}$, $\rho_{reg}$, $\rho_{0,is}$, $\rho_{kan,is}$, or $\rho_{reg,is}$) as well as uncertainty in nutrient uptake rates based on equations in Stukel et al. (submitted). There are two types of files included in this folders: Matlab scripts (.m files) and Microsoft Excel spreadsheets (.xlsx).

Each Matlab script quantifies nutrient uptake rates and uncertainty for a specific equation. Scripts that do not end in MC quantify symmetric confidence limits (e.g. nutrient uptake = $\rho \pm \sigma$). Scripts that end in MC use a Monte Carlo approach to quantify asymmetric confidence limits (e.g. nutrient uptake = $\rho$, but may vary from $\rho_{low}$ to $\rho_{high}$). Note that for both types of files, uncertainty outputs depend on uncertainty inputs. Specifically, if uncertainty in input parameters is given as standard deviation, the uncertainty in nutrient uptake rate should be treated as the standard deviation. If uncertainty in input parameters is given as 95% confidence limits, the uncertainty in nutrient uptake rate should be treated 95% confidence limits.

The three Excel spreadsheets each quantify nutrient uptake rates and corresponding uncertainty for all 6 equations. All Excel spreadsheets report symmetric confidence limits. The spreadsheet **'N Uptake Uncertainty Calculations.xlsx'** was designed to be used for investigation of the results from each of the 6 equations.

The spreadsheet **'Nutrient Uptake Log Sheet.xlsx'** was designed to be used as a log sheet that can be used to quantify nutrient uptake rates and uncertainties for multiple experiments. It should give the exact same answers as spreadsheet 'N Uptake Uncertainty Calculations.xlsx'.

The spreadsheet **'Nutrient Uptake Log Sheet - replicates.xlsx'** was designed to be used as a log sheet for quantifying nutrient uptake rates and corresponding uncertainty when replicate measurements are made. Please note that it is currently set up for quadruplicate measurements. When used with triplicate or duplicate measurements it may be necessary (depending on the version of Excel being used) to modify the spreadsheet slightly by deleting cells T9:LC9 (for triplicate measurements) or T8:LC9 (for duplicate measurements).
