## Supplementary material for "Investigating equations for measuring dissolved inorganic nutrient uptake in oligotrophic conditions": Calculation Files for Nutrient Uptake Eqs: ComputeRho0.m

```
function [rho0,sig_rho0] = ...
    ComputeRho0(T,P,IpT,Ip0,Ispk,Iamb,Nspk,Namb,sig_T,sig_P,sig_IpT,sig_Ip0,sig_Ispk,sig_Iamb,sig_Nspk,sig_Namb)

% This code calculates nutrient uptake rate (rho0), based on the Equations 
% of Dugdale and Wilkerson (1986), which is written as Eq. 1 in Stukel et 
% al. (submitted) and uncertainty estimates (symmetric) for rho0 based on
% equations in Stukel et al. (submitted).  

% Input parameters are:
% T = duration of incubation
% P = particulate concentration at end of incubation
% IpT = Isotope ratio of particulate pool at end of incubation
% Ip0 = Isotope ratio of particulate pool at beginning of incubation
% Ispk = Isotope ratio of tracer spike
% Iamb = Isotope ratio of ambient nutrient pool
% Nspk = Concentration of tracer spike
% Namb = Concentration of ambient nutrient pool
% sig_T = uncertainty in duration of incubation
% sig_P = uncertainty in particulate concentration at end of incubation
% sig_IpT = uncertainty in Isotope ratio of particulate pool at end of incubation
% sig_Ip0 = uncertainty in Isotope ratio of particulate pool at beginning of incubation
% sig_Ispk = uncertainty in Isotope ratio of tracer spike
% sig_Iamb = uncertainty in Isotope ratio of ambient nutrient pool
% sig_Nspk = uncertainty in Concentration of tracer spike
% sig_Namb = uncertainty in Concentration of ambient nutrient pool

Is0 = (Ispk.*Nspk+Iamb.*Namb)/(Nspk+Namb);

rho0 = P./T.*(IpT-Ip0)./(Is0-Ip0);

drho0dT = -rho0./T;
drho0dP = rho0./P;
drho0dIpT = rho0./(IpT-Ip0);
drho0dIp0 = rho0./(Is0 - Ip0) - P./T./(Is0 - Ip0);
drho0dIspk = -rho0./(Is0 - Ip0) .* Nspk./(Nspk+Namb);
drho0dIamb = -rho0./(Is0 - Ip0) .* Namb./(Nspk+Namb);
drho0dNspk = -rho0./(Is0 - Ip0) .* (Ispk - Is0)./(Nspk+Namb);
drho0dNamb = -rho0./(Is0 - Ip0) .* (Iamb - Is0)./(Nspk+Namb);

sig_rho02 = drho0dT.^2    .*   sig_T.^2 + ...
            drho0dP.^2    .*   sig_P.^2 + ...
            drho0dIpT.^2  .*   sig_IpT.^2 + ...
            drho0dIp0.^2  .*   sig_Ip0.^2 + ...
            drho0dIspk.^2 .*   sig_Ispk.^2 + ...
            drho0dIamb.^2 .*   sig_Iamb.^2 + ...
            drho0dNspk.^2 .*   sig_Nspk.^2 + ...
            drho0dNamb.^2 .*   sig_Namb.^2;
        
sig_rho0 = sig_rho02.^0.5;
```
