## Supplementary material for "Investigating equations for measuring dissolved inorganic nutrient uptake in oligotrophic conditions": Calculation Files for Nutrient Uptake Eqs: ComputeRho0is.m

```
function [rho0is,sig_rho0is] = ...
    ComputeRho0is(T,P,IpT,Ip0,Ispk,Iamb,Nspk,Namb,L10KS,sig_T,sig_P,sig_IpT,sig_Ip0,sig_Ispk,sig_Iamb,sig_Nspk,sig_Namb,sig_L10KS)

% This code calculates nutrient uptake rate (rho0,is), based on the Equations 
% of Dugdale and Wilkerson (1986), with a modification to account for 
% increased nutrient uptake in the incubation bottle (relative to 
% in situ conditions) resulting from the nutrient spike.  This is written 
% as Eq. 5 in Stukel et al. (submitted). This code also calculates 
% uncertainty estimates (symmetric) for rho0,is based on equations in Stukel 
% et al. (submitted).  

% Input parameters are:
% T = duration of incubation
% P = particulate concentration at end of incubation
% IpT = Isotope ratio of particulate pool at end of incubation
% Ip0 = Isotope ratio of particulate pool at beginning of incubation
% Ispk = Isotope ratio of tracer spike
% Iamb = Isotope ratio of ambient nutrient pool
% Nspk = Concentration of tracer spike
% Namb = Concentration of ambient nutrient pool
% L10KS = log-base 10 transformation of the half-saturation constant% sig_T = uncertainty in duration of incubation
% sig_P = uncertainty in particulate concentration at end of incubation
% sig_IpT = uncertainty in Isotope ratio of particulate pool at end of incubation
% sig_Ip0 = uncertainty in Isotope ratio of particulate pool at beginning of incubation
% sig_Ispk = uncertainty in Isotope ratio of tracer spike
% sig_Iamb = uncertainty in Isotope ratio of ambient nutrient pool
% sig_Nspk = uncertainty in Concentration of tracer spike
% sig_Namb = uncertainty in Concentration of ambient nutrient pool
% sig_L10KS = uncertainty in log-base 10 transformation of the half-saturation constant

Ks = 10.^L10KS;

Is0 = (Ispk.*Nspk+Iamb.*Namb)/(Nspk+Namb);

rho0 = P./T.*(IpT-Ip0)./(Is0-Ip0);

drho0dT = -rho0./T;
drho0dP = rho0./P;
drho0dIpT = rho0./(IpT-Ip0);
drho0dIp0 = rho0./(Is0 - Ip0) - P./T./(Is0 - Ip0);
drho0dIspk = -rho0./(Is0 - Ip0) .* Nspk./(Nspk+Namb);
drho0dIamb = -rho0./(Is0 - Ip0) .* Namb./(Nspk+Namb);
drho0dNspk = -rho0./(Is0 - Ip0) .* (Ispk - Is0)./(Nspk+Namb);
drho0dNamb = -rho0./(Is0 - Ip0) .* (Iamb - Is0)./(Nspk+Namb);

% sig_rho02 = drho0dT.^2    .*   sig_T.^2 + ...
%             drho0dP.^2    .*   sig_P.^2 + ...
%             drho0dIpT.^2  .*   sig_IpT.^2 + ...
%             drho0dIp0.^2  .*   sig_Ip0.^2 + ...
%             drho0dIspk.^2 .*   sig_Ispk.^2 + ...
%             drho0dIamb.^2 .*   sig_Iamb.^2 + ...
%             drho0dNspk.^2 .*   sig_Nspk.^2 + ...
%             drho0dNamb.^2 .*   sig_Namb.^2;
%         
% sig_rho0 = sig_rho02.^0.5;

y = Namb./(Nspk+Namb);
z = (Nspk + Namb + Ks)./(Namb + Ks);

rho0is = rho0 .* y .* z;

drho0isdP = y .* z .* drho0dP;
drho0isdT = y .* z .* drho0dT;
drho0isdIp0 = y .* z .* drho0dIp0;
drho0isdIpT = y .* z .* drho0dIpT;
drho0isdIspk = y .* z .* drho0dIspk;
drho0isdIamb = y .* z .* drho0dIamb;

drho0isdL10KS = rho0 .* y .* ( -log(10).*Nspk.*Ks ./ ((Namb+Ks).^2) );

dydNamb = Nspk ./ ((Nspk+Namb).^2);
dzdNamb = -Nspk ./ ((Ks+Namb).^2);
dydNspk = -Namb ./ ((Nspk+Namb).^2);
dzdNspk = 1/(Ks+Namb);

drho0isdNamb = drho0dNamb .* y .* z    +    dydNamb .* rho0 .* z    +    dzdNamb .* rho0 .* y;
drho0isdNspk = drho0dNspk .* y .* z    +    dydNspk .* rho0 .* z    +    dzdNspk .* rho0 .* y;

sig_rho0is2 = drho0isdT.^2    .*   sig_T.^2 + ...
            drho0isdP.^2    .*   sig_P.^2 + ...
            drho0isdIpT.^2  .*   sig_IpT.^2 + ...
            drho0isdIp0.^2  .*   sig_Ip0.^2 + ...
            drho0isdIspk.^2 .*   sig_Ispk.^2 + ...
            drho0isdIamb.^2 .*   sig_Iamb.^2 + ...
            drho0isdNspk.^2 .*   sig_Nspk.^2 + ...
            drho0isdNamb.^2 .*   sig_Namb.^2 + ...
            drho0isdL10KS.^2.*   sig_L10KS.^2;
        

sig_rho0is = sig_rho0is2.^0.5;        
```
