## Supplementary material for "Investigating equations for measuring dissolved inorganic nutrient uptake in oligotrophic conditions": Calculation Files for Nutrient Uptake Eqs: ComputeRho0isMC.m

```
function [rho0is,conf_rho0is] = ...
    ComputeRho0isMC(T,P,IpT,Ip0,Ispk,Iamb,Nspk,Namb,L10KS,sig_T,sig_P,sig_IpT,sig_Ip0,sig_Ispk,sig_Iamb,sig_Nspk,sig_Namb,sig_L10KS)

% This code calculates nutrient uptake rate (rho0,is), based on the Equations 
% of Dugdale and Wilkerson (1986), with a modification to account for 
% increased nutrient uptake in the incubation bottle (relative to 
% in situ conditions) resulting from the nutrient spike.  This is written 
% as Eq. 5 in Stukel et al. (submitted). This code also calculates 
% asymmetric confidence limits for rho0,is using a Monte Carlo approach.  

% Input parameters are:
% T = duration of incubation
% P = particulate concentration at end of incubation
% IpT = Isotope ratio of particulate pool at end of incubation
% Ip0 = Isotope ratio of particulate pool at beginning of incubation
% Ispk = Isotope ratio of tracer spike
% Iamb = Isotope ratio of ambient nutrient pool
% Nspk = Concentration of tracer spike
% Namb = Concentration of ambient nutrient pool
% L10KS = log-base 10 transformation of the half-saturation constant
% sig_T = uncertainty in duration of incubation
% sig_P = uncertainty in particulate concentration at end of incubation
% sig_IpT = uncertainty in Isotope ratio of particulate pool at end of incubation
% sig_Ip0 = uncertainty in Isotope ratio of particulate pool at beginning of incubation
% sig_Ispk = uncertainty in Isotope ratio of tracer spike
% sig_Iamb = uncertainty in Isotope ratio of ambient nutrient pool
% sig_Nspk = uncertainty in Concentration of tracer spike
% sig_Namb = uncertainty in Concentration of ambient nutrient pool
% sig_L10KS = uncertainty in log-base 10 transformation of the half-saturation constant


[rho0is] = ComputeRho0is(T,P,IpT,Ip0,Ispk,Iamb,Nspk,Namb,L10KS,sig_T,sig_P,sig_IpT,sig_Ip0,sig_Ispk,sig_Iamb,sig_Nspk,sig_Namb,sig_L10KS);

N = 1000; %Number of simulations to conduct

for i=1:N
    T2 = T + randn*sig_T;
    P2 = P + randn*sig_P;
    IpT2 = IpT + randn*sig_IpT;
    Ip02 = Ip0 + randn*sig_Ip0;
    Ispk2 = Ispk + randn*sig_Ispk;
    Iamb2 = Iamb + randn*sig_Iamb;
    Nspk2 = Nspk + randn*sig_Nspk;
    Namb2 = Namb + randn*sig_Namb;
    L10KS2 = L10KS + randn*sig_L10KS;
    Is0 = (Ispk2.*Nspk2+Iamb2.*Namb2)./(Nspk2+Namb2);
    if T2<=0 | Ispk2<=Iamb2
        rho_temp(i)=Inf;
    elseif P2<=0 | IpT2<=Ip02 | Namb2<=0
        rho_temp(i)=0;
    elseif Is0 <= Ip02
        rho_temp(i)=Inf;
    else
        temp=ComputeRho0is(T2,P2,IpT2,Ip02,Ispk2,Iamb2,Nspk2,Namb2,L10KS2,sig_T,sig_P,sig_IpT,sig_Ip0,sig_Ispk,sig_Iamb,sig_Nspk,sig_Namb,sig_L10KS);
        rho_temp(i)=temp;
    end
end
rho_temp=sort(rho_temp);
conf_rho0is = [rho_temp(round(N*0.15865)) rho_temp(round(N*0.84135))];
```
