## Supplementary material for "Investigating equations for measuring dissolved inorganic nutrient uptake in oligotrophic conditions": Calculation Files for Nutrient Uptake Eqs: ComputeRhoKanda.m

```
function [rhokan,sig_rhokan] = ...
    ComputeRhoKanda(T,P,IpT,Ip0,Ispk,Iamb,Nspk,Namb,a,sig_T,sig_P,sig_IpT,sig_Ip0,sig_Ispk,sig_Iamb,sig_Nspk,sig_Namb,sig_a)

% This code calculates nutrient uptake rate (rhokan), based on the Equations 
% of Kanda et al. (1987), which is written as Eq. 3 in Stukel et 
% al. (submitted) and uncertainty estimates (symmetric) for rhokan based on
% equations in Stukel et al. (submitted).  

% Input parameters are:
% T = duration of incubation
% P = particulate concentration at end of incubation
% IpT = Isotope ratio of particulate pool at end of incubation
% Ip0 = Isotope ratio of particulate pool at beginning of incubation
% Ispk = Isotope ratio of tracer spike
% Iamb = Isotope ratio of ambient nutrient pool
% Nspk = Concentration of tracer spike
% Namb = Concentration of ambient nutrient pool
% a = ratio of nutrient regeneration to nutrient uptake
% sig_T = uncertainty in duration of incubation
% sig_P = uncertainty in particulate concentration at end of incubation
% sig_IpT = uncertainty in Isotope ratio of particulate pool at end of incubation
% sig_Ip0 = uncertainty in Isotope ratio of particulate pool at beginning of incubation
% sig_Ispk = uncertainty in Isotope ratio of tracer spike
% sig_Iamb = uncertainty in Isotope ratio of ambient nutrient pool
% sig_Nspk = uncertainty in Concentration of tracer spike
% sig_Namb = uncertainty in Concentration of ambient nutrient pool
% sig_a = uncertainty in ratio of nutrient regeneration to nutrient uptake

Is0 = (Ispk.*Nspk+Iamb.*Namb)./(Nspk+Namb);

rho0 = P./T.*(IpT-Ip0)./(Is0-Ip0);

b = rho0.*T./(Namb+Nspk);

if a==1
    %if a is equal to zero, g_x and h_x are both equal to zero, but g_x*h_x
    %reduces to -1/b*ln(1-b).  To include uncertainty in a in the
    %derivation if a is assumed to be equal to one, I use this value for
    %g_x*h_x when calculating rhokan, but change a to 0.999 to calculate
    %sig_rhokan
    a = 0.99999;
end

g_x = -1 + (1-b).^(1-a);
h_x = 1./((a-1).*b);
rhokan = rho0.*g_x.*h_x;

drho0dT = -rho0./T;
drho0dP = rho0./P;
drho0dIpT = rho0./(IpT-Ip0);
drho0dIp0 = rho0./(Is0 - Ip0) - P./T./(Is0 - Ip0);
drho0dIspk = -rho0./(Is0 - Ip0) .* Nspk./(Nspk+Namb);
drho0dIamb = -rho0./(Is0 - Ip0) .* Namb./(Nspk+Namb);
drho0dNspk = -rho0./(Is0 - Ip0) .* (Ispk - Is0)./(Nspk+Namb);
drho0dNamb = -rho0./(Is0 - Ip0) .* (Iamb - Is0)./(Nspk+Namb);


drhokanda = -log(1-b) .* (1-b).^(1-a) .* rho0 .* h_x    -   rho0 .* g_x ./ (b .* (a-1).^2);

dgdP = (a-1)./(1-b).^a .* T./(Namb+Nspk) .* drho0dP;
dgdT = 0;
dgdIpT = (a-1)./(1-b).^a .* T./(Namb+Nspk) .* drho0dIpT;
dgdIp0 = (a-1)./(1-b).^a .* T./(Namb+Nspk) .* drho0dIp0;
dgdNamb = (a-1)./(1-b).^a .* (-b./(Namb+Nspk) + T./(Namb+Nspk) .* drho0dNamb);
dgdNspk = (a-1)./(1-b).^a .* (-b./(Namb+Nspk) + T./(Namb+Nspk) .* drho0dNspk);
dgdIspk = (a-1)./(1-b).^a .* T./(Namb+Nspk) .* drho0dIspk;
dgdIamb = (a-1)./(1-b).^a .* T./(Namb+Nspk) .* drho0dIamb;

dhdP = -1./((a-1).*b.^2) .* T./(Namb+Nspk) .* drho0dP;
dhdT = 0;
dhdIpT = -1./((a-1).*b.^2) .* T./(Namb+Nspk) .* drho0dIpT;
dhdIp0 = -1./((a-1).*b.^2) .* T./(Namb+Nspk) .* drho0dIp0;
dhdNamb = -1./((a-1).*b.^2) .* (-b./(Namb+Nspk) + T./(Namb+Nspk) .* drho0dNamb);
dhdNspk = -1./((a-1).*b.^2) .* (-b./(Namb+Nspk) + T./(Namb+Nspk) .* drho0dNspk);
dhdIspk = -1./((a-1).*b.^2) .* T./(Namb+Nspk) .* drho0dIspk;
dhdIamb = -1./((a-1).*b.^2) .* T./(Namb+Nspk) .* drho0dIamb;

drhokandP = g_x.*h_x.*drho0dP + rho0.*h_x.*dgdP + rho0.*g_x.*dhdP ;
drhokandT = g_x.*h_x.*drho0dT + rho0.*h_x.*dgdT + rho0.*g_x.*dhdT ;
drhokandIpT = g_x.*h_x.*drho0dIpT + rho0.*h_x.*dgdIpT + rho0.*g_x.*dhdIpT ;
drhokandIp0 = g_x.*h_x.*drho0dIp0 + rho0.*h_x.*dgdIp0 + rho0.*g_x.*dhdIp0 ;
drhokandNamb = g_x.*h_x.*drho0dNamb + rho0.*h_x.*dgdNamb + rho0.*g_x.*dhdNamb ;
drhokandNspk = g_x.*h_x.*drho0dNspk + rho0.*h_x.*dgdNspk + rho0.*g_x.*dhdNspk ;
drhokandIspk = g_x.*h_x.*drho0dIspk + rho0.*h_x.*dgdIspk + rho0.*g_x.*dhdIspk ;
drhokandIamb = g_x.*h_x.*drho0dIamb + rho0.*h_x.*dgdIamb + rho0.*g_x.*dhdIamb ;


sig_rhokan2 = drhokanda.^2    .*   sig_a.^2 + ...
            drhokandT.^2    .*   sig_T.^2 + ...
            drhokandP.^2    .*   sig_P.^2 + ...
            drhokandIpT.^2  .*   sig_IpT.^2 + ...
            drhokandIp0.^2  .*   sig_Ip0.^2 + ...
            drhokandIspk.^2 .*   sig_Ispk.^2 + ...
            drhokandIamb.^2 .*   sig_Iamb.^2 + ...
            drhokandNspk.^2 .*   sig_Nspk.^2 + ...
            drhokandNamb.^2 .*   sig_Namb.^2;
        
sig_rhokan = sig_rhokan2.^0.5;

if a==1
    %if a is equal to zero, g_x and h_x are both equal to zero, but g_x*h_x
    %reduces to -1/b*ln(1-b).  To include uncertainty in a in the
    %derivation if a is assumed to be equal to one, I use this value for
    %g_x*h_x when calculating rhokan, but change a to 0.999 to calculate
    %sig_rhokan
    rhokan = rho0.*(-1./b).*log(1-b);
else
    rhokan = rho0.*g_x.*h_x;
end
```
