## Supplementary material for "Investigating equations for measuring dissolved inorganic nutrient uptake in oligotrophic conditions": Calculation Files for Nutrient Uptake Eqs: ComputeRhoReg.m

```
function [rhoreg,sig_rhoreg] = ...
    ComputeRhoReg(T,P,IpT,Ip0,Ispk,Iamb,Nspk,Namb,a,sig_T,sig_P,sig_IpT,sig_Ip0,sig_Ispk,sig_Iamb,sig_Nspk,sig_Namb,sig_a)

% This code calculates nutrient uptake rate (rhoreg), assuming nutrient 
% regeneration within the incubation bottle, using Eq. 13 in Stukel et 
% al. (submitted) and uncertainty estimates (symmetric) for rhoreg based on
% equations in Stukel et al. (submitted).  

% Input parameters are:
% T = duration of incubation
% P = particulate concentration at end of incubation
% IpT = Isotope ratio of particulate pool at end of incubation
% Ip0 = Isotope ratio of particulate pool at beginning of incubation
% Ispk = Isotope ratio of tracer spike
% Iamb = Isotope ratio of ambient nutrient pool
% Nspk = Concentration of tracer spike
% Namb = Concentration of ambient nutrient pool
% a = ratio of nutrient regeneration to nutrient uptake
% sig_T = uncertainty in duration of incubation
% sig_P = uncertainty in particulate concentration at end of incubation
% sig_IpT = uncertainty in Isotope ratio of particulate pool at end of incubation
% sig_Ip0 = uncertainty in Isotope ratio of particulate pool at beginning of incubation
% sig_Ispk = uncertainty in Isotope ratio of tracer spike
% sig_Iamb = uncertainty in Isotope ratio of ambient nutrient pool
% sig_Nspk = uncertainty in Concentration of tracer spike
% sig_Namb = uncertainty in Concentration of ambient nutrient pool
% sig_a = uncertainty in ratio of nutrient regeneration to nutrient uptake

Is0 = (Ispk.*Nspk+Iamb.*Namb)./(Nspk+Namb);
S = Nspk + Namb;

%rhoreg = P/T*S/(P+a*S) * ( log((Is0-a*Ip0)/P) - log((Ip0*P+Is0*S)/(P*S) - (P+a*S)/(P*S)*IpT   ) );
                                                
f = log( (Is0 - a*Ip0)/P);
g = log( (Ip0 - IpT)/S + (Is0-a*IpT)/P);
h = P*S/(P+a*S);

rhoreg = (f - g)*h / T;

 
% if abs(rhoreg2-rhoreg)>0.0000001
%     rhoreg2-rhoreg;
%     'Something is wrong'
%     breakitty
% end

dfdp = -1/P;
dgdp = -(Is0 - a*IpT) / (P * ( ( (Ip0-IpT)/S)*P + (Is0 - a*IpT)));
dhdp = a * S^2 / (P + a*S)^2;
drhodp = (dfdp - dgdp)*h/T + (f-g)*dhdp/T;

% drhodp = S*P*( (Is0-a*IpT)/( ( (Is0-a*IpT)/P + (Ip0-IpT)/S ) * P^2 ) - 1/P ) / (T*(P+a*S)) + ...
%     S * ( log((Is0-a*Ip0)/P) - log( (Is0-a*IpT)/P + (Ip0-IpT)/S ) ) / (T*(P+a*S)) - ...
%     S*P * ( log((Is0-a*Ip0)/P) - log( (Is0-a*IpT)/P + (Ip0-IpT)/S ) ) / (T*(P+a*S)^2);

drhodT = -1/T * rhoreg;

dfdIp0 = a / (a*Ip0-Is0);
dgdIp0 = P / ( (Ip0-IpT)*P + S*(Is0-a*IpT));
drhodIp0 = (dfdIp0 - dgdIp0)*h/T;

dgdIpT = (P + a*S) / ((P + a*S)*IpT - S*Is0 - Ip0*P);
drhodIpT = -dgdIpT*h/T;

dfdIspk = 1 / (Is0 - a*Ip0) * Nspk/S;
dgdIspk = Nspk / ((Is0 - a*IpT)*S + (Ip0-IpT)*P);
drhodIspk = (dfdIspk - dgdIspk)*h/T;

dfdIamb = 1 / (Is0 - a*Ip0) * Namb/S;
dgdIamb = 1 / (Is0 + P*((Ip0-IpT)/S) - a*IpT) * Namb/S;
drhodIamb = (dfdIamb - dgdIamb)*h/T;

dfdNspk = (Iamb-Ispk)*Namb / (S*((a*Ip0-Ispk)*Nspk + (a*Ip0-Iamb)*Namb));
dgdNspk = -(P*IpT+(Ispk-Iamb)*Namb-P*Ip0) / ( S*((a*IpT-Ispk)*Nspk + (a*Namb+P)*IpT - Iamb*Namb - P*Ip0) );
dhdNspk = P^2 / (a*Nspk + a*Namb + P)^2;
drhodNspk = (dfdNspk - dgdNspk)*h/T + (f-g)*dhdNspk/T;

dfdNamb = -(Iamb-Ispk)*Nspk / (S*((a*Ip0-Ispk)*Nspk + (a*Ip0-Iamb)*Namb));
dgdNamb = -(P*IpT+(Iamb-Ispk)*Nspk-P*Ip0) / ( S*((a*IpT-Iamb)*Namb + (a*Nspk+P)*IpT - Ispk*Nspk - P*Ip0) );
dhdNamb = P^2 / (a*Nspk + a*Namb + P)^2;
drhodNamb = (dfdNamb - dgdNamb)*h/T + (f-g)*dhdNamb/T;

dfda = Ip0 / (a*Ip0 - Is0);
dgda = IpT / (a*IpT - Is0 - P*(Ip0-IpT)/S);
dhda = -P*S^2/ ((S*a+P)^2);
drhoda = (dfda - dgda)*h/T + (f-g)*dhda/T;


% drhoda = P*S * ( IpT/(P * ((Is0-IpT*a)/P + (Ip0-IpT)/S)) - Ip0/(Is0-Ip0*a) ) / (T*(S*a+P)) - ...
%     P*S^2 * ( log((Is0-a*Ip0)/P) - log( (Ip0-IpT)/S + (Is0-a*IpT)/P ) ) / (T * (S*a+P)^2);

% drhoda = P*S * ( IpT/(P * ((Is0-IpT*a)/P + (Ip0-IpT)/S)) - Ip0/(Is0-Ip0*a) ) / (T*(S*a+P)) + ...
%     (f-g)/T * dhda;


sig_rhoreg = sqrt( drhodp^2*sig_P^2 + drhodT^2*sig_T^2 + drhodIp0^2*sig_Ip0^2 + drhodIpT^2*sig_IpT^2 + ...
    drhodIspk^2*sig_Ispk^2 + drhodIamb^2*sig_Iamb^2 + drhodNspk^2*sig_Nspk^2 + drhodNamb^2*sig_Namb^2 + drhoda^2*sig_a^2);
```
