## Supplementary material for "Investigating equations for measuring dissolved inorganic nutrient uptake in oligotrophic conditions": Calculation Files for Nutrient Uptake Eqs: ComputeRhoRegis.m

```
function [rhoregis,sig_rhoregis] = ...
    ComputeRhoRegis(T,P,IpT,Ip0,Ispk,Iamb,Nspk,Namb,a,L10KS,sig_T,sig_P,sig_IpT,sig_Ip0,sig_Ispk,sig_Iamb,sig_Nspk,sig_Namb,sig_a,sig_L10KS)

% This code calculates nutrient uptake rate (rhoreg,is), assuming nutrient 
% regeneration within the incubation bottle, , with a modification to account for 
% increased nutrient uptake in the incubation bottle (relative to 
% in situ conditions) resulting from the nutrient spike.  This is written 
% as Eq. 14 in Stukel et al. (submitted). This code also calculates 
% asymmetric confidence limits for rhokan,is using a Monte Carlo approach. 

% Input parameters are:
% T = duration of incubation
% P = particulate concentration at end of incubation
% IpT = Isotope ratio of particulate pool at end of incubation
% Ip0 = Isotope ratio of particulate pool at beginning of incubation
% Ispk = Isotope ratio of tracer spike
% Iamb = Isotope ratio of ambient nutrient pool
% Nspk = Concentration of tracer spike
% Namb = Concentration of ambient nutrient pool
% a = ratio of nutrient regeneration to nutrient uptake
% L10KS = log-base 10 transformation of the half-saturation constant
% sig_T = uncertainty in duration of incubation
% sig_P = uncertainty in particulate concentration at end of incubation
% sig_IpT = uncertainty in Isotope ratio of particulate pool at end of incubation
% sig_Ip0 = uncertainty in Isotope ratio of particulate pool at beginning of incubation
% sig_Ispk = uncertainty in Isotope ratio of tracer spike
% sig_Iamb = uncertainty in Isotope ratio of ambient nutrient pool
% sig_Nspk = uncertainty in Concentration of tracer spike
% sig_Namb = uncertainty in Concentration of ambient nutrient pool
% sig_a = uncertainty in ratio of nutrient regeneration to nutrient uptake
% sig_L10KS = uncertainty in log-base 10 transformation of the half-saturation constant

Is0 = (Ispk.*Nspk+Iamb.*Namb)./(Nspk+Namb);
S = Nspk + Namb;
Ks = 10^L10KS;
                                                
f = log( (Is0 - a*Ip0)/P);
g = log( (Ip0 - IpT)/S + (Is0-a*IpT)/P);
h = P*S/(P+a*S);
y = Namb/(Nspk+Namb);
z = (Nspk+Namb+Ks)/(Namb+Ks);


rhoreg = (f - g)*h / T;
rhoregis = (f - g)*h / T * y * z;

 
% if abs(rhoreg2-rhoreg)>0.0000001
%     rhoreg2-rhoreg;
%     'Something is wrong'
%     breakitty
% end

dfdp = -1/P;
dgdp = -(Is0 - a*IpT) / (P * ( ( (Ip0-IpT)/S)*P + (Is0 - a*IpT)));
dhdp = a * S^2 / (P + a*S)^2;
drhodp = (dfdp - dgdp)*h/T + (f-g)*dhdp/T;
drhodp = drhodp*y*z;

% drhodp = S*P*( (Is0-a*IpT)/( ( (Is0-a*IpT)/P + (Ip0-IpT)/S ) * P^2 ) - 1/P ) / (T*(P+a*S)) + ...
%     S * ( log((Is0-a*Ip0)/P) - log( (Is0-a*IpT)/P + (Ip0-IpT)/S ) ) / (T*(P+a*S)) - ...
%     S*P * ( log((Is0-a*Ip0)/P) - log( (Is0-a*IpT)/P + (Ip0-IpT)/S ) ) / (T*(P+a*S)^2);

drhodT = -1/T * rhoreg;
drhodT = drhodT*y*z;

dfdIp0 = a / (a*Ip0-Is0);
dgdIp0 = P / ( (Ip0-IpT)*P + S*(Is0-a*IpT));
drhodIp0 = (dfdIp0 - dgdIp0)*h/T;
drhodIp0 = drhodIp0*y*z;

dgdIpT = (P + a*S) / ((P + a*S)*IpT - S*Is0 - Ip0*P);
drhodIpT = -dgdIpT*h/T;
drhodIpT = drhodIpT*y*z;

dfdIspk = 1 / (Is0 - a*Ip0) * Nspk/S;
dgdIspk = Nspk / ((Is0 - a*IpT)*S + (Ip0-IpT)*P);
drhodIspk = (dfdIspk - dgdIspk)*h/T;
drhodIspk = drhodIspk*y*z;

dfdIamb = 1 / (Is0 - a*Ip0) * Namb/S;
dgdIamb = 1 / (Is0 + P*((Ip0-IpT)/S) - a*IpT) * Namb/S;
drhodIamb = (dfdIamb - dgdIamb)*h/T;
drhodIamb = drhodIamb*y*z;

dfdNspk = (Iamb-Ispk)*Namb / (S*((a*Ip0-Ispk)*Nspk + (a*Ip0-Iamb)*Namb));
dgdNspk = -(P*IpT+(Ispk-Iamb)*Namb-P*Ip0) / ( S*((a*IpT-Ispk)*Nspk + (a*Namb+P)*IpT - Iamb*Namb - P*Ip0) );
dhdNspk = P^2 / (a*Nspk + a*Namb + P)^2;
drhodNspk = (dfdNspk - dgdNspk)*h/T + (f-g)*dhdNspk/T;
dydNspk = -Namb/(Nspk+Namb)^2;
dzdNspk = 1/(Ks+Namb);
drhodNspk = drhodNspk*y*z + dydNspk*rhoreg*z + dzdNspk*rhoreg*y;


dfdNamb = -(Iamb-Ispk)*Nspk / (S*((a*Ip0-Ispk)*Nspk + (a*Ip0-Iamb)*Namb));
dgdNamb = -(P*IpT+(Iamb-Ispk)*Nspk-P*Ip0) / ( S*((a*IpT-Iamb)*Namb + (a*Nspk+P)*IpT - Ispk*Nspk - P*Ip0) );
dhdNamb = P^2 / (a*Nspk + a*Namb + P)^2;
drhodNamb = (dfdNamb - dgdNamb)*h/T + (f-g)*dhdNamb/T;
dydNamb = Nspk / (Nspk+Namb)^2;
dzdNamb = -Nspk / (Ks+Namb)^2;
drhodNamb = drhodNamb*y*z + dydNamb*rhoreg*z + dzdNamb*rhoreg*y;

dfda = Ip0 / (a*Ip0 - Is0);
dgda = IpT / (a*IpT - Is0 - P*(Ip0-IpT)/S);
dhda = -P*S^2/ ((S*a+P)^2);
drhoda = (dfda - dgda)*h/T + (f-g)*dhda/T;
drhoda = drhoda*y*z;

drhodL10KS = rhoreg * (Namb/(Nspk+Namb)) * ( (-log(10)*Nspk*Ks)/(Namb+Ks)^2);

sig_rhoregis = sqrt( drhodp^2*sig_P^2 + drhodT^2*sig_T^2 + drhodIp0^2*sig_Ip0^2 + drhodIpT^2*sig_IpT^2 + ...
    drhodIspk^2*sig_Ispk^2 + drhodIamb^2*sig_Iamb^2 + drhodNspk^2*sig_Nspk^2 + drhodNamb^2*sig_Namb^2 + ...
    drhoda^2*sig_a^2 + drhodL10KS^2*sig_L10KS^2);
```
