## Supplementary material for "Investigating equations for measuring dissolved inorganic nutrient uptake in oligotrophic conditions": NEMURO+15N Files: 1 - Read Me - NEMURON15.pdf

This is code for the version of NEMURO+$^{15}$N that was used to investigate nutrient uptake rate equations in Stukel et al. (submitted, L&O Methods). For additional details about the model, see Kishi et al. (2007), Stukel et al. (2018, PLOS ONE), and Stukel et al. (submitted, L&O Methods). To run the model, all that you should need to do is copy all of the files into a single folder and then run the file 'RunNEMURON15.m' from that folder using Matlab. This code was developed using Matlab R2016a. While I believe that it is backwards compatible, I do not guarantee it.

Please not that model duration is set at line 8 and layer thickness can be set at line 10. The model is currently configured for a one-dimensional model with 2-m layer thickness to a depth of 300 m. The variable ParamSet can be changed to use different sets of parameters in NEMURO, to investigate model sensitivity to these parameters.

For questions and comments, please contact Mike Stukel.
