## Supplementary material for "Investigating equations for measuring dissolved inorganic nutrient uptake in oligotrophic conditions": NEMURO+15N Files: LightMod.m

```
function[lightbottom]=LightMod(PhyN,lightin,alpha1,alpha2,layerthickness)

% alpha1 = 0.04;    %m-1
% alpha2 = 0.04;    %umol N L-1 m-1

kappa = alpha1 + alpha2*PhyN;

lightmid = lightin .* exp(-kappa.*layerthickness/2);
lightbottom = lightin .* exp(-kappa.*layerthickness);
lightave = -lightin./layerthickness.*(exp(-kappa.*layerthickness)./kappa-1./kappa);
```
