## Supplementary material for "Investigating equations for measuring dissolved inorganic nutrient uptake in oligotrophic conditions": NEMURO+15N Files: Mixing.m

```
function[tracernew,tracer15Nnew]=Mixing(tracer,tracer15N,LayerThickness,dt,z,numlevs,DeepNO3,DeepSi,Deep15NNO3,DeepNH4,Deep15NNH4,Kzmixed,Kzdeep,MLD)

Kz=ones(size(z))*Kzdeep;
Kz(find(z<MLD))=Kzmixed;

Kz(find(z<MLD*2 &z>MLD))=Kzdeep+((MLD*2-z(find(z<MLD*2 & z>MLD)))/MLD).^2.*(Kzmixed-Kzdeep);
Kz=Kz*24*60*60;
deltaz=LayerThickness;

deep=zeros(1,11);
deep(1,6)=DeepNO3;
deep(1,7)=DeepNH4;
deep(1,10)=DeepSi;

deep15N=zeros(1,9);
deep15N(6)=DeepNO3*Deep15NNO3;
deep15N(7)=DeepNH4*Deep15NNH4;

tracernew=tracer;
tracer15Nnew=tracer15N;

tracer(numlevs+1,1:11)=deep;
tracer15N(numlevs+1,:)=deep15N;

deriv=(tracer(1:end-1,:)-tracer(2:end,:))/deltaz;
Flux=zeros(size(tracernew));
for i=1:11
    Flux(:,i)=-deriv(:,i).*Kz'*dt;   
end

tracernew(1:end,:)=tracernew(1:end,:)+Flux(1:end,:);
tracernew(2:end,:)=tracernew(2:end,:)-Flux(1:end-1,:);


deriv15N=(tracer15N(1:end-1,:)-tracer15N(2:end,:))/deltaz;
Flux15N=zeros(size(tracer15Nnew));
for i=1:9
    Flux15N(:,i)=-deriv15N(:,i).*Kz'*dt;   
end

tracer15Nnew(1:end,:)=tracer15Nnew(1:end,:)+Flux15N(1:end,:);
tracer15Nnew(2:end,:)=tracer15Nnew(2:end,:)-Flux15N(1:end-1,:);


% deriv15N=(tracer15N(1:end-1,:)-tracer15N(2:end,:))/deltaz;
% Flux15N=zeros(size(tracer15Nnew));
% for i=1:9
%     Flux15N(:,i)=-deriv15N(:,i).*Kz'*dt;   
% end
% 
% tracer15Nnew(1:end,:)=tracer15Nnew(1:end,:)+Flux15N(1:end,:);
% tracer15Nnew(2:end,:)=tracer15Nnew(2:end,:)-Flux15N(2:end,:);
```
