## Supplementary material for "Investigating equations for measuring dissolved inorganic nutrient uptake in oligotrophic conditions": NEMURO+15N Files: NEMURO_params15N.m

```
function[param15N]=NEMURO_params15N(input)


%Nitrogen Assimilation
param15N(1)=-5;           %Eps_NO3up
param15N(2)=-10;          %Eps_NH4up

%Excretion
param15N(3)=-5;           %Eps_ZsExc
param15N(4)=-5;           %Eps_ZlExc
param15N(5)=-5;           %Eps_ZpExc

%Egestion
param15N(6)=-2;           %Eps_ZsEg
param15N(7)=-2;           %Eps_ZlEg
param15N(8)=-2;           %Eps_ZpEg

%Nitrification
param15N(9)=-14;          %Eps_Nit

%Remineralization
param15N(10)=-1;          %Eps_P2D
param15N(11)=-1;          %Eps_P2N
param15N(12)=-1;          %Eps_D2N

%Nitrogen Isotopic Ratio of N2
param15N(13)=0.0036765;           %RN2
```
