## Supplementary material for "Investigating equations for measuring dissolved inorganic nutrient uptake in oligotrophic conditions": NEMURO+15N Files: NEMURO_params.m

```
function[param]=NEMURO_params(input)

if strcmp(input,'Li')
    
    %Parameters for Underwater Light Attenuation
    param(:,1)=0.04;       %Alpha1
    param(:,2)=0.04;       %Alpha2
    
    %Parameters for small phytoplankton (PS)
    param(:,3)=0.4;        %VmaxS
    param(:,4)=1;          %KNO3S
    param(:,5)=0.1;        %KNH4S
    param(:,6)=1.5;        %PsiS
    param(:,7)=0.0693;     %kGppS
    %param(:,8)=104.7;      %IoptS
    param(:,9)=0.03;       %ResPS0
    param(:,10)=0.0519;    %kResPS
    param(:,11)=0.135;     %gamS
    param(:,12)=0.0585;    %MorPS0
    param(:,13)=0.0693;    %kMorPS
    
    %Parameters for large phytoplankton (PL)
    param(:,14)=1.0;       %VmaxL
    param(:,15)=3;         %KNO3L
    param(:,16)=0.3;       %KNH4L
    param(:,17)=4;         %KSiL
    param(:,18)=4.6;       %PsiL
    param(:,19)=0.0693;    %kGppL
    %param(:,20)=104.7;     %IoptL
    param(:,21)=0.03;      %ResPL0
    param(:,22)=0.0519;    %kResPL
    param(:,23)=0.135;     %gamL
    param(:,24)=0.029;     %MorPL0
    param(:,25)=0.0693;    %kMorPL
    
    %Parameters for small zooplankton (ZS)
    param(:,26)=0.4;       %GrmaxSPS
    param(:,27)=0.0693;    %kGraS
    param(:,28)=1.4;       %lamS
    param(:,29)=0.04;      %PS2ZSstar
    param(:,30)=0.0585;    %MorZS0
    param(:,31)=0.0693;    %kMorZS
    param(:,32)=0.7;       %AlphaZS
    param(:,33)=0.3;       %BetaZS
    
    %Parameters for large zooplankton (ZL)
    param(:,34)=0.1;       %GrmaxLPS
    param(:,35)=0.4;       %GrmaxLPL
    param(:,36)=0.4;       %GrmaxLZS
    param(:,37)=0.0693;    %kGraL
    param(:,38)=1.4;       %lamL
    param(:,39)=0.04;      %PS2ZLstar
    param(:,40)=0.04;      %PL2ZLstar
    param(:,41)=0.04;      %ZS2ZLstar
    param(:,42)=0.0585;    %MorZL0
    param(:,43)=0.0693;    %kMorZL
    param(:,44)=0.7;       %AlphaZL
    param(:,45)=0.3;       %BetaZL
    
    %Parameters for predatory zooplankton (ZP)
    param(:,46)=0.2;       %GrmaxPPL
    param(:,47)=0.2;       %GrmaxPZS
    param(:,48)=0.2;       %GrmaxPZL
    param(:,49)=0.0693;    %kGraP
    param(:,50)=1.4;       %lamP
    param(:,51)=0.04;      %PL2ZPstar
    param(:,52)=0.04;      %ZS2ZPstar
    param(:,53)=0.04;      %ZL2ZPstar
    param(:,54)=4.605;     %PsiPL
    param(:,55)=3.01;      %PsiZS
    param(:,56)=0.0585;    %MorZP0
    param(:,57)=0.0693;    %kMorZP
    param(:,58)=0.7;       %AlphaZP
    param(:,59)=0.3;       %BetaZP
    
    %Other Parameters (decompostion, etc.)
    param(:,60)=0.03;      %Nit0
    param(:,61)=0.0693;    %kNit
    param(:,62)=0.1;       %VP2N0
    param(:,63)=0.0693;    %kP2N
    param(:,64)=0.01;      %VP2D0
    param(:,65)=0.0693;    %kP2D
    param(:,66)=0.02;      %VD2N0
    param(:,67)=0.0693;    %kD2N
    param(:,68)=0.1;       %VP2Si0
    param(:,69)=0.0693;    %kP2Si
    param(:,70)=0;         %RSiNPS
    param(:,71)=1;         %RSiNPL
    param(:,72)=0;         %RSiNZS
    param(:,73)=0;         %RSiNZL
    param(:,74)=40;       %setVP
    param(:,75)=40;       %setVO
    
    %Parameters for the Light Parameterization from Li et al. (2010)
    param(:,76)=0.0030;     %AlphaPS
    param(:,77)=0.0061;     %AlphaPL
    param(:,78)=0.0017;     %BetaPS
    param(:,79)=0.0002;     %BetaPL
    
elseif strcmp(input,'Fortran')
    
    %Parameters for Underwater Light Attenuation
    param(:,1)=0.04;       %Alpha1
    param(:,2)=0.04;       %Alpha2
    
    %Parameters for small phytoplankton (PS)
    param(:,3)=0.4;        %VmaxS
    param(:,4)=1;          %KNO3S
    param(:,5)=0.1;        %KNH4S
    param(:,6)=1.5;        %PsiS
    param(:,7)=0.0693;     %kGppS
    %param(:,8)=104.7;      %IoptS
    param(:,9)=0.03;       %ResPS0
    param(:,10)=0.0519;    %kResPS
    param(:,11)=0.135;     %gamS
    param(:,12)=0.0585;    %MorPS0
    param(:,13)=0.0693;    %kMorPS
    
    %Parameters for large phytoplankton (PL)
    param(:,14)=0.8;       %VmaxL
    param(:,15)=3;         %KNO3L
    param(:,16)=0.3;       %KNH4L
    param(:,17)=6;         %KSiL
    param(:,18)=1.5;       %PsiL
    param(:,19)=0.0693;    %kGppL
    %param(:,20)=104.7;     %IoptL
    param(:,21)=0.03;      %ResPL0
    param(:,22)=0.0519;    %kResPL
    param(:,23)=0.135;     %gamL
    param(:,24)=0.029;     %MorPL0
    param(:,25)=0.0693;    %kMorPL
    
    %Parameters for small zooplankton (ZS)
    param(:,26)=0.4;       %GrmaxSPS
    param(:,27)=0.0693;    %kGraS
    param(:,28)=1.4;       %lamS
    param(:,29)=0.04;      %PS2ZSstar
    param(:,30)=0.0585;    %MorZS0
    param(:,31)=0.0693;    %kMorZS
    param(:,32)=0.7;       %AlphaZS
    param(:,33)=0.3;       %BetaZS
    
    %Parameters for large zooplankton (ZL)
    param(:,34)=0.1;       %GrmaxLPS
    param(:,35)=0.4;       %GrmaxLPL
    param(:,36)=0.4;       %GrmaxLZS
    param(:,37)=0.0693;    %kGraL
    param(:,38)=1.4;       %lamL
    param(:,39)=0.04;      %PS2ZLstar
    param(:,40)=0.04;      %PL2ZLstar
    param(:,41)=0.04;      %ZS2ZLstar
    param(:,42)=0.0585;    %MorZL0
    param(:,43)=0.0693;    %kMorZL
    param(:,44)=0.7;       %AlphaZL
    param(:,45)=0.3;       %BetaZL
    
    %Parameters for predatory zooplankton (ZP)
    param(:,46)=0.2;       %GrmaxPPL
    param(:,47)=0.2;       %GrmaxPZS
    param(:,48)=0.2;       %GrmaxPZL
    param(:,49)=0.0693;    %kGraP
    param(:,50)=1.4;       %lamP
    param(:,51)=0.04;      %PL2ZPstar
    param(:,52)=0.04;      %ZS2ZPstar
    param(:,53)=0.04;      %ZL2ZPstar
    param(:,54)=4.605;     %PsiPL
    param(:,55)=3.01;      %PsiZS
    param(:,56)=0.0585;    %MorZP0
    param(:,57)=0.0693;    %kMorZP
    param(:,58)=0.7;       %AlphaZP
    param(:,59)=0.3;       %BetaZP
    
    %Other Parameters (decompostion, etc.)
    param(:,60)=0.03;      %Nit0
    param(:,61)=0.0693;    %kNit
    param(:,62)=0.1;       %VP2N0
    param(:,63)=0.0693;    %kP2N
    param(:,64)=0.01;      %VP2D0
    param(:,65)=0.0693;    %kP2D
    param(:,66)=0.02;      %VD2N0
    param(:,67)=0.0693;    %kD2N
    param(:,68)=0.1;       %VP2Si0
    param(:,69)=0.0693;    %kP2Si
    param(:,70)=0;         %RSiNPS
    param(:,71)=1;         %RSiNPL
    param(:,72)=0;         %RSiNZS
    param(:,73)=0;         %RSiNZL
    param(:,74)=40;       %setVP
    param(:,75)=40;       %setVO
    
    %Parameters for the Light Parameterization from Fortran nemuro.in
    param(:,76)=0.01;     %AlphaPS
    param(:,77)=0.01;     %AlphaPL
    param(:,78)=0.00045;     %BetaPS
    param(:,79)=0.00045;     %BetaPL
    
    
end
```
