## Supplementary material for "Investigating equations for measuring dissolved inorganic nutrient uptake in oligotrophic conditions": NEMURO+15N Files: NEMURON15.m

```
function[tracernew,tracer15Nnew]=NEMURON15_excrete(tracer,tracer15N,param,param15N,paramDOM,dt,inputs)

%In this version of the code, I set the phytoplankton to excrete DOM rather
%than respire a mixture of NH4 and NO3.  

if strcmp(inputs,'HTLout')
    Zoo2PON=0;
else
    Zoo2PON=1;
end


% dt=0.1;
% 
% [param]=NEMURO_params(0);
% [param15N]=NEMURO_params15N(0);
% 
% tracer(1,1:11)=0.1;
% tracer(1,12)=100;
% tracer(1,13)=12;
% 
% tracer15N=tracer(1,1:9)*0.0036765;

%This is the start of the true code

PS=tracer(:,1);
PL=tracer(:,2);
ZS=tracer(:,3);
ZL=tracer(:,4);
ZP=tracer(:,5);
NO3=tracer(:,6);
NH4=tracer(:,7);
PON=tracer(:,8);
DON=tracer(:,9);
Si=tracer(:,10);
Opal=tracer(:,11);
Irr=tracer(:,12);
TMP=tracer(:,13);

N15PS=tracer15N(:,1);
N15PL=tracer15N(:,2);
N15ZS=tracer15N(:,3);
N15ZL=tracer15N(:,4);
N15ZP=tracer15N(:,5);
N15NO3=tracer15N(:,6);
N15NH4=tracer15N(:,7);
N15PON=tracer15N(:,8);
N15DON=tracer15N(:,9);

%Parameters---------------------------------------------------------
%[param]=NEMURO_tracers(0);

%Parameters for Underwater Light Attenuation
Alpha1=param(:,1);       %
Alpha2=param(:,2);       %

%Parameters for small phytoplankton (PS)
VmaxS=param(:,3);        %
KNO3S=param(:,4);          %
KNH4S=param(:,5);        %
PsiS=param(:,6);        %
kGppS=param(:,7);     %
%IoptS=param(:,8);      %
ResPS0=param(:,9);       %
kResPS=param(:,10);    %
gamS=param(:,11);     %
MorPS0=param(:,12);    %
kMorPS=param(:,13);    %

%Parameters for large phytoplankton (PL)
VmaxL=param(:,14);       %
KNO3L=param(:,15);         %
KNH4L=param(:,16);       %
KSiL=param(:,17);         %
PsiL=param(:,18);       %
kGppL=param(:,19);    %
%IoptL=param(:,20);     %
ResPL0=param(:,21);      %
kResPL=param(:,22);    %
gamL=param(:,23);     %
MorPL0=param(:,24);     %
kMorPL=param(:,25);    %

%Parameters for small zooplankton (ZS)
GrmaxSPS=param(:,26);       %
kGraS=param(:,27);    %
lamS=param(:,28);       %
PS2ZSstar=param(:,29);      %
MorZS0=param(:,30);    %
kMorZS=param(:,31);    %
AlphaZS=param(:,32);       %
BetaZS=param(:,33);       %

%Parameters for large zooplankton (ZL)
GrmaxLPS=param(:,34);       %
GrmaxLPL=param(:,35);       %
GrmaxLZS=param(:,36);       %
kGraL=param(:,37);    %
lamL=param(:,38);       %
PS2ZLstar=param(:,39);      %
PL2ZLstar=param(:,40);      %
ZS2ZLstar=param(:,41);      %
MorZL0=param(:,42);    %
kMorZL=param(:,43);    %
AlphaZL=param(:,44);       %
BetaZL=param(:,45);       %

%Parameters for predatory zooplankton (ZP)
GrmaxPPL=param(:,46);       %
GrmaxPZS=param(:,47);       %
GrmaxPZL=param(:,48);       %
kGraP=param(:,49);    %
lamP=param(:,50);       %
PL2ZPstar=param(:,51);      %
ZS2ZPstar=param(:,52);      %
ZL2ZPstar=param(:,53);      %
PsiPL=param(:,54);     %
PsiZS=param(:,55);      %
MorZP0=param(:,56);    %
kMorZP=param(:,57);    %
AlphaZP=param(:,58);       %
BetaZP=param(:,59);       %

%Other Parameters (decompostion, etc.)
Nit0=param(:,60);      %
kNit=param(:,61);    %
VP2N0=param(:,62);       %
kP2N=param(:,63);    %
VP2D0=param(:,64);      %
kP2D=param(:,65);    %
VD2N0=param(:,66);      %
kD2N=param(:,67);    %
VP2Si0=param(:,68);       %
kP2Si=param(:,69);    %
RSiNPS=param(:,70);         %
RSiNPL=param(:,71);         %
RSiNZS=param(:,72);         %
RSiNZL=param(:,73);         %
setVP=param(:,74);       %
setVO=param(:,75);       %

%Parameters for the Light Parameterization from Li et al. (2010)
AlphaPS=param(:,76);     %
AlphaPL=param(:,77);     %
BetaPS=param(:,78);     %
BetaPL=param(:,79);     %

%15N Parameters
Eps_NO3up=param15N(1);          
Eps_NH4up=param15N(2);          %
Eps_ZsExc=param15N(3);           %
Eps_ZlExc=param15N(4);           %
Eps_ZpExc=param15N(5);           %
Eps_ZsEg=param15N(6);           %
Eps_ZlEg=param15N(7);           %
Eps_ZpEg=param15N(8);           %
Eps_Nit=param15N(9);          %
Eps_P2D=param15N(10);          %
Eps_P2N=param15N(11);          %
Eps_D2N=param15N(12);          %
RN2=param15N(13);           %

Alpha_NO3up=exp(Eps_NO3up/1000);
Alpha_NH4up=exp(Eps_NH4up/1000);
Alpha_ZsExc=exp(Eps_ZsExc/1000);
Alpha_ZlExc=exp(Eps_ZlExc/1000);
Alpha_ZpExc=exp(Eps_ZpExc/1000);
Alpha_ZsEg=exp(Eps_ZsEg/1000);
Alpha_ZlEg=exp(Eps_ZlEg/1000);
Alpha_ZpEg=exp(Eps_ZpEg/1000);
Alpha_Nit=exp(Eps_Nit/1000);
Alpha_P2D=exp(Eps_P2D/1000);
Alpha_P2N=exp(Eps_P2N/1000);
Alpha_D2N=exp(Eps_D2N/1000);


%Small phytoplankton photosynthesis
NitLimS = NO3./(NO3+KNO3S).*exp(-PsiS.*NH4);
AmmLimS = NH4./(NH4+KNH4S);
%LigLimS = Irr./IoptS.*exp(1-Irr./IoptS);  %Note that light limitation is handled differently in different versions of NEMURO.  Here I am using the Kishi et al. light parameterization
LigLimS = (1-exp(-AlphaPS.*Irr/VmaxS)).*exp(-BetaPS.*Irr/VmaxS);
GppPSn = VmaxS.*(NitLimS+AmmLimS).*exp(kGppS.*TMP).*LigLimS.*PS;
RnewS = NitLimS./(NitLimS+AmmLimS);

%Large phytoplankton photosynthesis
NitLimL = NO3./(NO3+KNO3L).*exp(-PsiL.*NH4);
AmmLimL = NH4./(NH4+KNH4L);
SiLimL  = Si./(Si + KSiL);
%LigLimL = Irr./IoptL.*exp(1-Irr./IoptL);  %Note that light limitation is handled differently in different versions of NEMURO.  Here I am using the Kishi et al. light parameterization
LigLimL = (1-exp(-AlphaPL.*Irr/VmaxL)).*exp(-BetaPL.*Irr/VmaxL);
GppPLn  = VmaxL.*min(NitLimL+AmmLimL,SiLimL).*exp(kGppL.*TMP).*LigLimL.*PL;
RnewL = NitLimL./(NitLimL+AmmLimL);

%Small Phyto Parameters
ResPSn = ResPS0*exp(kResPS.*TMP).*PS;
MorPSn = MorPS0*exp(kMorPS.*TMP).*PS.^2;
ExcPSn = gamS.*GppPSn;

%Large Phyto Parameters
ResPLn = ResPL0*exp(kResPL.*TMP).*PL;
MorPLn = MorPL0*exp(kMorPL.*TMP).*PL.^2;
ExcPLn = gamL.*GppPLn;

%Protozoan Grazing
GraPS2ZSn = max(0,GrmaxSPS*exp(kGraS.*TMP).*(1-exp(lamS.*(PS2ZSstar-PS))).*ZS);


%Mesozooplankton Grazing
GraPS2ZLn = max(0,GrmaxLPS*exp(kGraL.*TMP).*(1-exp(lamL.*(PS2ZLstar-PS))).*ZL);
GraPL2ZLn = max(0,GrmaxLPL*exp(kGraL.*TMP).*(1-exp(lamL.*(PL2ZLstar-PL))).*ZL);
GraZS2ZLn = max(0,GrmaxLZS*exp(kGraL.*TMP).*(1-exp(lamL.*(ZS2ZLstar-ZS))).*ZL);

%Predatory Zooplankton Grazing
GraPL2ZPn = max(0,GrmaxPPL.*exp(kGraP.*TMP).*(1-exp(lamP.*(PL2ZPstar-PL))).* ...
    exp(-PsiPL.*(ZL+ZS)).*ZP);
GraZS2ZPn = max(0,GrmaxPZS.*exp(kGraP.*TMP).*(1-exp(lamP.*(ZS2ZPstar-ZS))).* ...
    exp(-PsiZS.*ZL).*ZP);
GraZL2ZPn = max(0,GrmaxPZL.*exp(kGraP.*TMP).*(1-exp(lamP.*(ZL2ZPstar-ZL))).*ZP);

%Zooplankton Excretion & Egestion
ExcZSn = (AlphaZS - BetaZS)*GraPS2ZSn;
ExcZLn = (AlphaZL - BetaZL)*(GraPS2ZLn+GraPL2ZLn+GraZS2ZLn);
ExcZPn = (AlphaZP - BetaZP)*(GraPL2ZPn+GraZS2ZPn+GraZL2ZPn);
EgeZSn = (1-AlphaZS)*GraPS2ZSn;
EgeZLn = (1-AlphaZL)*(GraPS2ZLn+GraPL2ZLn+GraZS2ZLn);
EgeZPn = (1-AlphaZP)*(GraPL2ZPn+GraZS2ZPn+GraZL2ZPn);

%Mortality
MorZSn = MorZS0*exp(kMorZS.*TMP).*ZS.^2;
MorZLn = MorZL0*exp(kMorZL.*TMP).*ZL.^2;
MorZPn = MorZP0*exp(kMorZP.*TMP).*ZP.^2;

%Decomposition
DecPON2NH4 = VP2N0*exp(kP2N.*TMP).*PON;
DecPON2DON = VP2D0*exp(kP2D.*TMP).*PON;
DecDON2NH4 = VD2N0*exp(kD2N.*TMP).*DON;

%Nitrification
Nit = Nit0.*exp(kNit.*TMP).*NH4;

%Silicate
GppPLsi = GppPLn*RSiNPL;
ResPLsi = ResPLn*RSiNPL;
MorPLsi = MorPLn*RSiNPL;
ExcPLsi = ExcPLn*RSiNPL;
EgeZLsi = GraPL2ZLn*RSiNPL;
EgeZPsi = GraPL2ZPn*RSiNPL;
DecOpal2Si = VP2Si0*exp(kP2Si*TMP).*Opal;

RPS = N15PS./max(PS,10^-20);
RPL = N15PL./max(PL,10^-20);
RZS = N15ZS./max(ZS,10^-20);
RZL = N15ZL./max(ZL,10^-20);
RZP = N15ZP./max(ZP,10^-20);
RNO3 = N15NO3./max(NO3,10^-20);
RNH4 = N15NH4./max(NH4,10^-20);
RDON = N15DON./max(DON,10^-20);
RPON = N15PON./max(PON,10^-20);

dPSdt=0;
dPLdt=0;
dZSdt=0;
dZLdt=0;
dZPdt=0;
dNO3dt=0;
dNH4dt=0;
dPONdt=0;
dDONdt=0;
dSidt=0;
dOpaldt=0;

d15NPSdt=0;
d15NPLdt=0;
d15NZSdt=0;
d15NZLdt=0;
d15NZPdt=0;
d15NNO3dt=0;
d15NNH4dt=0;
d15NPONdt=0;
d15NDONdt=0;

dPSdt = dPSdt + GppPSn;
dNO3dt = dNO3dt - GppPSn.*RnewS;
dNH4dt = dNH4dt - GppPSn.*(1-RnewS);


 d15NPSdt = d15NPSdt + (GppPSn.*RnewS).*RNO3.*Alpha_NO3up;
 d15NNO3dt = d15NNO3dt - (GppPSn.*RnewS).*RNO3.*Alpha_NO3up;
 d15NPSdt = d15NPSdt + (GppPSn.*(1-RnewS) ).*RNH4.*Alpha_NH4up;
 d15NNH4dt = d15NNH4dt - (GppPSn.*(1-RnewS) ).*RNH4.*Alpha_NH4up;
 
dPSdt = dPSdt - ResPSn;
dDONdt = dDONdt + ResPSn;

 d15NPSdt = d15NPSdt - ResPSn.*RPS;
 d15NDONdt = d15NDONdt + ResPSn.*RPS;
 
dPLdt = dPLdt + GppPLn;
dNO3dt = dNO3dt - GppPLn.*RnewL;
dNH4dt = dNH4dt - GppPLn.*(1-RnewL);

 d15NPLdt = d15NPLdt + (GppPLn.*RnewL).*RNO3.*Alpha_NO3up;
 d15NNO3dt = d15NNO3dt - (GppPLn.*RnewL).*RNO3.*Alpha_NO3up;
 d15NPLdt = d15NPLdt + (GppPLn.*(1-RnewL)).*RNH4.*Alpha_NH4up;
 d15NNH4dt = d15NNH4dt - (GppPLn.*(1-RnewL)).*RNH4.*Alpha_NH4up;
 
dPLdt = dPLdt - ResPLn;
dDONdt = dDONdt + ResPLn;

 d15NPLdt = d15NPLdt - ResPLn.*RPL;
 d15NDONdt = d15NDONdt + ResPLn.*RPL;
 
dZSdt = dZSdt + GraPS2ZSn;  
dPSdt = dPSdt - GraPS2ZSn;

 d15NZSdt = d15NZSdt + GraPS2ZSn.*RPS;
 d15NPSdt = d15NPSdt - GraPS2ZSn.*RPS;

dZLdt = dZLdt + GraPS2ZLn + GraPL2ZLn + GraZS2ZLn;
dZSdt = dZSdt - GraZS2ZLn;
dPLdt = dPLdt - GraPL2ZLn;
dPSdt = dPSdt - GraPS2ZLn;
 
 d15NZLdt = d15NZLdt + GraPS2ZLn.*RPS + GraPL2ZLn.*RPL + GraZS2ZLn.*RZS;
 d15NZSdt = d15NZSdt - GraZS2ZLn.*RZS;
 d15NPLdt = d15NPLdt - GraPL2ZLn.*RPL;
 d15NPSdt = d15NPSdt - GraPS2ZLn.*RPS;

dZPdt = dZPdt + GraPL2ZPn + GraZS2ZPn + GraZL2ZPn;
dPLdt = dPLdt - GraPL2ZPn;
dZSdt = dZSdt - GraZS2ZPn;
dZLdt = dZLdt - GraZL2ZPn;

 d15NZPdt = d15NZPdt + GraPL2ZPn.*RPL + GraZS2ZPn.*RZS + GraZL2ZPn.*RZL;
 d15NPLdt = d15NPLdt - GraPL2ZPn.*RPL;
 d15NZSdt = d15NZSdt - GraZS2ZPn.*RZS;
 d15NZLdt = d15NZLdt - GraZL2ZPn.*RZL;

% EgeZSn=0
% EgeZLn=0
% EgeZPn=0 
 
dPSdt = dPSdt - ExcPSn;
dPLdt = dPLdt - ExcPLn;
dZSdt = dZSdt - ExcZSn;
dZLdt = dZLdt - ExcZLn;
dZPdt = dZPdt - ExcZPn;
dNH4dt = dNH4dt + ExcZSn + ExcZLn + ExcZPn;
dDONdt = dDONdt + paramDOM*ExcPSn + paramDOM*ExcPLn;

 d15NPSdt = d15NPSdt - ExcPSn.*RPS;
 d15NPLdt = d15NPLdt - ExcPLn.*RPL;
 d15NZSdt = d15NZSdt - ExcZSn.*RZS*Alpha_ZsExc;
 d15NZLdt = d15NZLdt - ExcZLn.*RZL*Alpha_ZlExc;
 d15NZPdt = d15NZPdt - ExcZPn.*RZP*Alpha_ZpExc;
 d15NNH4dt = d15NNH4dt + ExcZSn.*RZS*Alpha_ZsExc + ExcZLn.*RZL*Alpha_ZlExc + ExcZPn.*RZP*Alpha_ZpExc;
 d15NDONdt = d15NDONdt + paramDOM*ExcPSn.*RPS + paramDOM*ExcPLn.*RPL;

dZSdt = dZSdt - EgeZSn;
dZLdt = dZLdt - EgeZLn;
dZPdt = dZPdt - EgeZPn;
dPONdt = dPONdt + EgeZSn + EgeZLn + EgeZPn;

 d15NZSdt = d15NZSdt - EgeZSn.*RZS*Alpha_ZsEg;
 d15NZLdt = d15NZLdt - EgeZLn.*RZL*Alpha_ZlEg;
 d15NZPdt = d15NZPdt - EgeZPn.*RZP*Alpha_ZpEg;
 d15NPONdt = d15NPONdt + EgeZSn.*RZS*Alpha_ZsEg + EgeZLn.*RZL*Alpha_ZlEg + EgeZPn.*RZP*Alpha_ZpEg;

dPSdt = dPSdt - MorPSn;
dPLdt = dPLdt - MorPLn;
dZSdt = dZSdt - MorZSn;
dZLdt = dZLdt - MorZLn;
dZPdt = dZPdt - MorZPn;
if Zoo2PON==1
    dPONdt = dPONdt + MorPSn + MorPLn + MorZSn + MorZLn + MorZPn;
else
    dPONdt = dPONdt + MorPSn + MorPLn + MorZSn;
end

d15NPSdt = d15NPSdt - MorPSn.*RPS;
d15NPLdt = d15NPLdt - MorPLn.*RPL;
d15NZSdt = d15NZSdt - MorZSn.*RZS;
d15NZLdt = d15NZLdt - MorZLn.*RZL;
d15NZPdt = d15NZPdt - MorZPn.*RZP;
if Zoo2PON==1
    d15NPONdt = d15NPONdt + MorPSn.*RPS + MorPLn.*RPL + ...
        MorZSn.*RZS + MorZLn.*RZL + MorZPn.*RZP;
else
    d15NPONdt = d15NPONdt + MorPSn.*RPS + MorPLn.*RPL + ...
        MorZSn.*RZS;
end


dPONdt = dPONdt - DecPON2NH4 - DecPON2DON;
dNH4dt = dNH4dt + DecPON2NH4;
dDONdt = dDONdt + paramDOM*DecPON2DON;
dDONdt = dDONdt - DecDON2NH4;
dNH4dt = dNH4dt + DecDON2NH4;

 d15NPONdt = d15NPONdt - DecPON2NH4.*RPON*Alpha_P2N - DecPON2DON.*RPON*Alpha_P2D;
 d15NNH4dt = d15NNH4dt + DecPON2NH4.*RPON*Alpha_P2N;
 d15NDONdt = d15NDONdt + paramDOM*DecPON2DON.*RPON*Alpha_P2D;
 d15NDONdt = d15NDONdt - DecDON2NH4.*RDON*Alpha_D2N;
 d15NNH4dt = d15NNH4dt + DecDON2NH4.*RDON*Alpha_D2N;

dNO3dt = dNO3dt + Nit;
dNH4dt = dNH4dt - Nit;

 d15NNO3dt = d15NNO3dt + Nit.*RNH4*Alpha_Nit;
 d15NNH4dt = d15NNH4dt - Nit.*RNH4*Alpha_Nit;

dSidt = dSidt + ExcPLsi + DecOpal2Si - (GppPLsi - ResPLsi);
dOpaldt = dOpaldt + MorPLsi + EgeZLsi + EgeZPsi - DecOpal2Si;

% dPLsidt = GppPLsi - ResPLsi - MorPLsi - ExcPLsi - GraPL2ZLsi - GraPL2ZPsi;
% dZLsidt = GraPL2ZLsi - EgeZLsi;
% dZPsidt = GraPL2ZPsi - EgeZPsi;

tracernew(:,1)=tracer(:,1)+dPSdt*dt;
tracernew(:,2)=tracer(:,2)+dPLdt*dt;
tracernew(:,3)=tracer(:,3)+dZSdt*dt;
tracernew(:,4)=tracer(:,4)+dZLdt*dt;
tracernew(:,5)=tracer(:,5)+dZPdt*dt;
tracernew(:,6)=tracer(:,6)+dNO3dt*dt;
tracernew(:,7)=tracer(:,7)+dNH4dt*dt;
tracernew(:,8)=tracer(:,8)+dPONdt*dt;
tracernew(:,9)=tracer(:,9)+dDONdt*dt;
tracernew(:,10)=tracer(:,10)+dSidt*dt;
tracernew(:,11)=tracer(:,11)+dOpaldt*dt;
tracernew(:,12)=tracer(:,12);
tracernew(:,13)=tracer(:,13);

 tracer15Nnew(:,1)=tracer15N(:,1)+d15NPSdt*dt;
 tracer15Nnew(:,2)=tracer15N(:,2)+d15NPLdt*dt;
 tracer15Nnew(:,3)=tracer15N(:,3)+d15NZSdt*dt;
 tracer15Nnew(:,4)=tracer15N(:,4)+d15NZLdt*dt;
 tracer15Nnew(:,5)=tracer15N(:,5)+d15NZPdt*dt;
 tracer15Nnew(:,6)=tracer15N(:,6)+d15NNO3dt*dt;
 tracer15Nnew(:,7)=tracer15N(:,7)+d15NNH4dt*dt;
 tracer15Nnew(:,8)=tracer15N(:,8)+d15NPONdt*dt;
 tracer15Nnew(:,9)=tracer15N(:,9)+d15NDONdt*dt;

% %Code for checking conservation
% sum(tracernew(:,1:9))-sum(tracer(:,1:9))
% (tracernew(:,2)*RSiNPL+tracernew(:,10)+tracernew(:,11))-(tracer(:,2)*RSiNPL+tracer(:,10)+tracer(:,11))
% sum(tracer15Nnew(:,1:9))-sum(tracer15N(:,1:9))
```
