## Supplementary material for "Investigating equations for measuring dissolved inorganic nutrient uptake in oligotrophic conditions": NEMURO+15N Files: RunNEMURON15.m

```
clear all
close all

ParamSet=1

dt=1/24/60;
dtdiff=1/24/60/10;
duration=365*2;
SurfIrr = 708.4;   %From Li et al (2010)
LayerThickness=2;
numlevs=150;
DeepNO3=10;
DeepSi=10;
DeepNH4=0;
atm15N=0.0036765;
Deep15NNO3 = 5.7/1000*atm15N + atm15N;
Deep15NNH4 = atm15N;
Temperature=20;
Kzmixed = 10^-3;    %Vertical Diffusivity Coefficient (m2/s)
Kzdeep = 10^-5;    %Vertical Diffusivity Coefficient (m2/s)
MLD=20;

z=0:LayerThickness:(numlevs-1)*LayerThickness;

%Defining Parameters
[param]=NEMURO_params('Li');
[param15N]=NEMURO_params15N(0);
paramDOM=1;
RN2=param15N(13);
[param,Kzmixed,Kzdeep,MLD,Temperature,SurfIrr]=SetParams(param,ParamSet,Kzmixed,Kzdeep,MLD,Temperature,SurfIrr);

%Initializing State Variables
tracer(1,1:11)=0.1;
tracer(1,12)=100;
tracer(1,13)=Temperature;
tracer15N=tracer(1,1:9)*RN2;
for i=2:numlevs
    tracer(i,:)=tracer(1,:);
    tracer(i,13)=20;
    tracer15N(i,:)=tracer15N(1,:);
end
tracer(i,6)=DeepNO3;
tracer(i,10)=DeepSi;
Alpha1=param(:,1);
Alpha2=param(:,2);
tracer0=tracer;

tracer15N0=tracer15N;

for i=1:duration/dt
    tracer(:,13)=Temperature;
    Phy = tracer(:,1)+tracer(:,2);
    Irr = LightMod(Phy(1),SurfIrr,Alpha1,Alpha2,LayerThickness);
    tracer(1,12)=Irr;
    for j=2:numlevs
        Irr = LightMod(Phy(2),Irr,Alpha1,Alpha2,LayerThickness);
        tracer(j,12)=Irr;
    end
    [tracernew,tracer15Nnew]=NEMURON15(tracer,tracer15N,param,param15N,paramDOM,dt,'HTLin');
    if i*dt==round(i*dt)
        
        tracer0(:,:,i*dt+1)=tracernew;
        tracer15N0(:,:,i*dt+1)=tracer15Nnew;
        time(i*dt+1)=i*dt+1;
    end
    tracer=tracernew;
    tracer15N=tracer15Nnew;
    
    [tracernew,tracer15Nnew]=Sinking(tracer,tracer15N,LayerThickness,param,dt);
    tracer=tracernew;
    tracer15N=tracer15Nnew;
    for j=1:dt/dtdiff
        [tracernew,tracer15Nnew]=Mixing(tracer,tracer15N,LayerThickness,dtdiff,z,numlevs,DeepNO3,DeepSi,Deep15NNO3,DeepNH4,Deep15NNH4,Kzmixed,Kzdeep,MLD);
        tracer=tracernew;
        tracer15N=tracer15Nnew;
    end
end
```
