## Supplementary material for "Investigating equations for measuring dissolved inorganic nutrient uptake in oligotrophic conditions": NEMURO+15N Files: SetParams.m

```
function [param_out,Kzmixed,Kzdeep,MLD,Temperature,SurfIrr]=SetParams(param,Trial,Kzmixed,Kzdeep,MLD,Temperature,SurfIrr)


if Trial==1
elseif Trial==2
    Kzdeep=10^-4;
elseif Trial==3
    Temperature=5;
    SurfIrr=100;
elseif Trial==4
    param(:,4)=0.1;          %KNO3S
    param(:,5)=0.01;        %KNH4S
    param(:,15)=0.3;         %KNO3L
    param(:,16)=0.03;       %KNH4L
elseif Trial==5
    param(:,4)=1;          %KNO3S
    param(:,5)=1;        %KNH4S
    param(:,15)=3;         %KNO3L
    param(:,16)=3;       %KNH4L
elseif Trial==6
    param(:,4)=0.1;          %KNO3S
    param(:,5)=0.1;        %KNH4S
    param(:,15)=0.3;         %KNO3L
    param(:,16)=0.3;       %KNH4L
elseif Trial==7
    param(:,60)=0.3;      %Nit0
elseif Trial==8
    param(:,60)=0.3;      %Nit0
    param(:,4)=0.1;          %KNO3S
    param(:,5)=0.01;        %KNH4S
    param(:,15)=0.3;         %KNO3L
    param(:,16)=0.03;       %KNH4L
elseif Trial==9
    param(:,60)=0.3;      %Nit0
    param(:,4)=0.1;          %KNO3S
    param(:,5)=0.1;        %KNH4S
    param(:,15)=0.3;         %KNO3L
    param(:,16)=0.3;       %KNH4L
elseif Trial==10
    param(:,60)=0.3;      %Nit0
    Kzdeep=10^-4;
elseif Trial==11
    param(:,60)=0.3;      %Nit0
    Temperature=5;
    SurfIrr=100;
elseif Trial==12
    param(:,60)=0.1;      %Nit0
elseif Trial==13
    param(:,6)=15;        %PsiS
elseif Trial==14
    param(:,12)=0.01;    %MorPS0
    param(:,24)=0.01;     %MorPL0
elseif Trial==15
    param(:,4)=1.0;          %KNO3S
    param(:,5)=0.01;        %KNH4S
    param(:,15)=3.0;         %KNO3L
    param(:,16)=0.03;       %KNH4L
elseif Trial==16
    param(:,4)=1.0;          %KNO3S
    param(:,5)=0.01;        %KNH4S
    param(:,15)=3.0;         %KNO3L
    param(:,16)=0.03;       %KNH4L
elseif Trial==17
    param(:,4)=1;          %KNO3S
    param(:,5)=1;        %KNH4S
    param(:,15)=3;         %KNO3L
    param(:,16)=3;       %KNH4L
    param(:,60)=0.1;      %Nit0
elseif Trial==18
    param(:,4)=1;          %KNO3S
    param(:,5)=1;        %KNH4S
    param(:,15)=3;         %KNO3L
    param(:,16)=3;       %KNH4L
    param(:,60)=0.1;      %Nit0
    Temperature=5;
    SurfIrr=100;
elseif Trial==19
    param(:,74)=5;       %setVP
    param(:,75)=5;       %setVO
elseif Trial==20
    param(:,74)=0;       %setVP
    param(:,75)=0;       %setVO
elseif Trial==21
    Kzmixed=10^-4;      
    param(:,74)=0;       %setVP
    param(:,75)=0;       %setVO
elseif Trial==22
    Kzmixed=10^-4;      
elseif Trial==23
    Kzmixed=10^-5;   
elseif Trial==24
    Kzmixed=10^-4;   
    MLD=50;  
elseif Trial==25
    Kzmixed=10^-5;      
    param(:,74)=0;       %setVP
    param(:,75)=0;       %setVO
elseif Trial==26
    Kzdeep=10^-4;
    param(:,62)=0.5;       %VP2N0
    param(:,64)=0.2;      %VP2D0
    param(:,66)=0.2;      %VD2N0
elseif Trial==27
    Kzdeep=10^-4;
    param(:,62)=0.5;       %VP2N0
    param(:,64)=0.2;      %VP2D0
    param(:,66)=0.2;      %VD2N0
    param(:,74)=4;       %setVP
    param(:,75)=4;       %setVO
elseif Trial==28
    Kzdeep=10^-4;
    param(:,62)=0.5;       %VP2N0
    param(:,64)=0.2;      %VP2D0
    param(:,66)=0.2;      %VD2N0
    param(:,4)=1;          %KNO3S
    param(:,5)=1;        %KNH4S
    param(:,15)=3;         %KNO3L
    param(:,16)=3;       %KNH4L
elseif Trial==29
    Kzdeep=10^-4;
    param(:,62)=0.5;       %VP2N0
    param(:,64)=0.2;      %VP2D0
    param(:,66)=0.2;      %VD2N0
    param(:,4)=3;          %KNO3S
    param(:,5)=1;        %KNH4S
    param(:,15)=10;         %KNO3L
    param(:,16)=3;       %KNH4L
elseif Trial==30
    Kzdeep=10^-4;
    param(:,62)=0.5;       %VP2N0
    param(:,64)=0.2;      %VP2D0
    param(:,66)=0.2;      %VD2N0
    param(:,56)=0.5;    %MorZP0
    param(:,42)=0.5;    %MorZL0
    param(:,30)=0.5;    %MorZS0
elseif Trial==31
    Kzmixed=10^-4;
    param(:,62)=0.5;       %VP2N0
    param(:,64)=0.2;      %VP2D0
    param(:,66)=0.2;      %VD2N0
    param(:,56)=0.5;    %MorZP0
    param(:,42)=0.5;    %MorZL0
    param(:,30)=0.5;    %MorZS0
elseif Trial==32
    param(:,4)=3;          %KNO3S
    param(:,5)=0.1;        %KNH4S
    param(:,15)=10;         %KNO3L
    param(:,16)=0.3;       %KNH4L
    param(:,62)=0.5;       %VP2N0
    param(:,64)=0.2;      %VP2D0
    param(:,66)=0.2;      %VD2N0
    param(:,56)=0.5;    %MorZP0
    param(:,42)=0.5;    %MorZL0
    param(:,30)=0.5;    %MorZS0
elseif Trial==33
    Kzdeep=10^-4;
    SurfIrr=50;
    param(:,62)=0.5;       %VP2N0
    param(:,64)=0.2;      %VP2D0
    param(:,66)=0.2;      %VD2N0
    param(:,56)=0.5;    %MorZP0
    param(:,42)=0.5;    %MorZL0
    param(:,30)=0.5;    %MorZS0
elseif Trial==34
    Kzdeep=10^-4;
    SurfIrr=50;
    param(:,62)=0.5;       %VP2N0
    param(:,64)=0.2;      %VP2D0
    param(:,66)=0.2;      %VD2N0
    param(:,56)=0.5;    %MorZP0
    param(:,42)=0.5;    %MorZL0
    param(:,30)=0.5;    %MorZS0
    param(:,32)=1.0;       %AlphaZS
elseif Trial==35
    Kzdeep=10^-4;
    SurfIrr=50;
    param(:,62)=0.5;       %VP2N0
    param(:,64)=0.2;      %VP2D0
    param(:,66)=0.2;      %VD2N0
    param(:,56)=0.5;    %MorZP0
    param(:,42)=0.5;    %MorZL0
    param(:,30)=0.5;    %MorZS0
    param(:,32)=1.0;       %AlphaZS
    param(:,39)=0.0004;      %PS2ZLstar
    param(:,34)=0.01;       %GrmaxLPS
elseif Trial==36
    Kzdeep=10^-4;
    param(:,62)=0.5;       %VP2N0
    param(:,64)=0.2;      %VP2D0
    param(:,66)=0.2;      %VD2N0
    param(:,56)=0.5;    %MorZP0
    param(:,42)=0.5;    %MorZL0
    param(:,30)=0.5;    %MorZS0
    param(:,60)=0.0;      %Nit0
elseif Trial==37    
    param(:,76)=0.0061;     %AlphaPS
    param(:,77)=0.0030;     %AlphaPL
elseif Trial==38    
    param(:,76)=0.002;     %AlphaPS
    param(:,77)=0.008;     %AlphaPL
elseif Trial==39    
    param(:,76)=0.0061;     %AlphaPS
    param(:,77)=0.0030;     %AlphaPL
    param(:,3)=0.8;        %VmaxS
elseif Trial==40    
    param(:,76)=0.0061;     %AlphaPS
    param(:,77)=0.0030;     %AlphaPL
    param(:,3)=0.6;        %VmaxS
elseif Trial==41   
    param(:,76)=0.0061;     %AlphaPS
    param(:,77)=0.0030;     %AlphaPL
    param(:,3)=0.5;        %VmaxS
    param(:,5)=0.05;        %KNH4S
    param(:,16)=1.0;       %KNH4L
elseif Trial==42   
    param(:,76)=0.0030;     %AlphaPS
    param(:,77)=0.0061;     %AlphaPL
    param(:,3)=0.5;        %VmaxS
    param(:,5)=0.05;        %KNH4S
    param(:,16)=1.0;       %KNH4L
elseif Trial==43   
    param(:,76)=0.0030;     %AlphaPS
    param(:,77)=0.0061;     %AlphaPL
    param(:,3)=0.6;        %VmaxS
    param(:,5)=0.05;        %KNH4S
    param(:,16)=1.0;       %KNH4L
elseif Trial==44   
    param(:,76)=0.0030;     %AlphaPS
    param(:,77)=0.0061;     %AlphaPL
    param(:,78)=0.0002;     %BetaPS
    param(:,79)=0.001;     %BetaPL
    param(:,3)=0.6;        %VmaxS
    param(:,5)=0.05;        %KNH4S
    param(:,16)=1.0;       %KNH4L
elseif Trial==45   
    param(:,76)=0.001;     %AlphaPS
    param(:,77)=0.0061;     %AlphaPL
    param(:,78)=0.0002;     %BetaPS
    param(:,79)=0.001;     %BetaPL
    param(:,3)=0.6;        %VmaxS
    param(:,5)=0.05;        %KNH4S
    param(:,16)=1.0;       %KNH4L
elseif Trial==46   
    param(:,76)=0.001;     %AlphaPS
    param(:,77)=0.0061;     %AlphaPL
    param(:,78)=0.0002;     %BetaPS
    param(:,79)=0.001;     %BetaPL
    param(:,3)=0.7;        %VmaxS
    param(:,5)=0.05;        %KNH4S
    param(:,16)=3.0;       %KNH4L
    param(:,56)=0.5;    %MorZP0
    param(:,42)=0.5;    %MorZL0
    param(:,30)=0.5;    %MorZS0
    param(:,62)=0.5;       %VP2N0
    param(:,64)=0.2;      %VP2D0
    param(:,66)=0.2;      %VD2N0
    param(:,32)=1.0;       %AlphaZS
    param(:,39)=0.0004;      %PS2ZLstar
    param(:,34)=0.01;       %GrmaxLPS
elseif Trial==47   
    param(:,76)=0.001;     %AlphaPS
    param(:,77)=0.0061;     %AlphaPL
    param(:,78)=0.0002;     %BetaPS
    param(:,79)=0.001;     %BetaPL
    param(:,3)=0.8;        %VmaxS
    param(:,5)=0.05;        %KNH4S
    param(:,16)=3.0;       %KNH4L
    param(:,56)=0.5;    %MorZP0
    param(:,42)=0.5;    %MorZL0
    param(:,30)=0.5;    %MorZS0
    param(:,62)=0.5;       %VP2N0
    param(:,64)=0.2;      %VP2D0
    param(:,66)=0.2;      %VD2N0
    param(:,32)=1.0;       %AlphaZS
    param(:,39)=0.0004;      %PS2ZLstar
    param(:,34)=0.01;       %GrmaxLPS
elseif Trial==48   
    Kzmixed=10^-4;
    param(:,76)=0.001;     %AlphaPS
    param(:,77)=0.0061;     %AlphaPL
    param(:,78)=0.0002;     %BetaPS
    param(:,79)=0.001;     %BetaPL
    param(:,3)=0.8;        %VmaxS
    param(:,5)=0.05;        %KNH4S
    param(:,16)=3.0;       %KNH4L
    param(:,56)=0.5;    %MorZP0
    param(:,42)=0.5;    %MorZL0
    param(:,30)=0.5;    %MorZS0
    param(:,62)=0.5;       %VP2N0
    param(:,64)=0.2;      %VP2D0
    param(:,66)=0.2;      %VD2N0
    param(:,32)=1.0;       %AlphaZS
    param(:,39)=0.0004;      %PS2ZLstar
    param(:,34)=0.01;       %GrmaxLPS
elseif Trial==49   
    Kzdeep=10^-4;
    param(:,76)=0.001;     %AlphaPS
    param(:,77)=0.0061;     %AlphaPL
    param(:,78)=0.0002;     %BetaPS
    param(:,79)=0.001;     %BetaPL
    param(:,3)=0.8;        %VmaxS
    param(:,5)=0.05;        %KNH4S
    param(:,16)=3.0;       %KNH4L
    param(:,56)=0.5;    %MorZP0
    param(:,42)=0.5;    %MorZL0
    param(:,30)=0.5;    %MorZS0
    param(:,62)=0.5;       %VP2N0
    param(:,64)=0.2;      %VP2D0
    param(:,66)=0.2;      %VD2N0
    param(:,32)=1.0;       %AlphaZS
    param(:,39)=0.0004;      %PS2ZLstar
    param(:,34)=0.01;       %GrmaxLPS
elseif Trial==50
    Kzmixed=10^-4;
    param(:,76)=0.001;     %AlphaPS
    param(:,77)=0.008;     %AlphaPL
    param(:,78)=0.0002;     %BetaPS
    param(:,79)=0.002;     %BetaPL
    param(:,3)=0.8;        %VmaxS
    param(:,5)=0.05;        %KNH4S
    param(:,16)=3.0;       %KNH4L
    param(:,56)=0.5;    %MorZP0
    param(:,42)=0.5;    %MorZL0
    param(:,30)=0.5;    %MorZS0
    param(:,62)=1;       %VP2N0
    param(:,64)=1;      %VP2D0
    param(:,66)=1;      %VD2N0
    param(:,32)=1.0;       %AlphaZS
    param(:,39)=0.0004;      %PS2ZLstar
    param(:,34)=0.01;       %GrmaxLPS
    param(:,74)=4;       %setVP
    param(:,75)=4;       %setVO
elseif Trial==51
    Kzmixed=10^-4;
    param(:,76)=0.001;     %AlphaPS
    param(:,77)=0.008;     %AlphaPL
    param(:,78)=0.0002;     %BetaPS
    param(:,79)=0.002;     %BetaPL
    param(:,3)=0.9;        %VmaxS
    param(:,5)=0.05;        %KNH4S
    param(:,16)=3.0;       %KNH4L
    param(:,56)=0.5;    %MorZP0
    param(:,42)=0.5;    %MorZL0
    param(:,30)=0.5;    %MorZS0
    param(:,62)=1;       %VP2N0
    param(:,64)=1;      %VP2D0
    param(:,66)=1;      %VD2N0
    param(:,32)=1.0;       %AlphaZS
    param(:,39)=0.0004;      %PS2ZLstar
    param(:,34)=0.01;       %GrmaxLPS
    param(:,74)=4;       %setVP
    param(:,75)=4;       %setVO
elseif Trial==52
    Kzmixed=10^-4;
    param(:,76)=0.001;     %AlphaPS
    param(:,77)=0.008;     %AlphaPL
    param(:,78)=0.0002;     %BetaPS
    param(:,79)=0.002;     %BetaPL
    param(:,3)=0.9;        %VmaxS
    param(:,5)=0.05;        %KNH4S
    param(:,16)=3.0;       %KNH4L
    param(:,56)=0.5;    %MorZP0
    param(:,42)=0.1;    %MorZL0
    param(:,30)=0.1;    %MorZS0
    param(:,62)=1;       %VP2N0
    param(:,64)=1;      %VP2D0
    param(:,66)=1;      %VD2N0
    param(:,32)=1.0;       %AlphaZS
    param(:,39)=0.0004;      %PS2ZLstar
    param(:,34)=0.01;       %GrmaxLPS
    param(:,74)=4;       %setVP
    param(:,75)=4;       %setVO
elseif Trial==53
    Kzmixed=10^-4;
    param(:,76)=0.001;     %AlphaPS
    param(:,77)=0.008;     %AlphaPL
    param(:,78)=0.0002;     %BetaPS
    param(:,79)=0.002;     %BetaPL
    param(:,3)=0.9;        %VmaxS
    param(:,5)=0.05;        %KNH4S
    param(:,16)=3.0;       %KNH4L
    param(:,56)=0.5;    %MorZP0
    param(:,42)=0.1;    %MorZL0
    param(:,30)=0.1;    %MorZS0
    param(:,62)=1;       %VP2N0
    param(:,64)=1;      %VP2D0
    param(:,66)=1;      %VD2N0
    param(:,32)=1.0;       %AlphaZS
    param(:,39)=0.0004;      %PS2ZLstar
    param(:,34)=0.01;       %GrmaxLPS
    param(:,28)=0.1;       %lamS   default = 1.4 (lower decreases grazing at low biomass)
    param(:,29)=0.04;      %PS2ZSstar   default = 0.04 (threshold)
    param(:,34)=0.001;       %GrmaxLPS   default = 0.1
    param(:,74)=4;       %setVP
    param(:,75)=4;       %setVO
elseif Trial==54
    Kzmixed=10^-4;
    param(:,76)=0.001;     %AlphaPS
    param(:,77)=0.008;     %AlphaPL
    param(:,78)=0.0002;     %BetaPS
    param(:,79)=0.002;     %BetaPL
    param(:,3)=0.9;        %VmaxS
    param(:,5)=0.05;        %KNH4S
    param(:,16)=3.0;       %KNH4L
    param(:,12)=0.1;    %MorPS0   default is 0.0585
    param(:,56)=0.5;    %MorZP0
    param(:,42)=0.1;    %MorZL0
    param(:,30)=0.1;    %MorZS0
    param(:,62)=1;       %VP2N0
    param(:,64)=1;      %VP2D0
    param(:,66)=1;      %VD2N0
    param(:,32)=1.0;       %AlphaZS
    param(:,39)=0.0004;      %PS2ZLstar
    param(:,34)=0.01;       %GrmaxLPS
    param(:,28)=0.1;       %lamS   default = 1.4 (lower decreases grazing at low biomass)
    param(:,29)=0.04;      %PS2ZSstar   default = 0.04 (threshold)
    param(:,34)=0.001;       %GrmaxLPS   default = 0.1
    param(:,74)=4;       %setVP
    param(:,75)=4;       %setVO
elseif Trial==55
    Kzmixed=10^-4;
    param(:,76)=0.001;     %AlphaPS
    param(:,77)=0.008;     %AlphaPL
    param(:,78)=0.0002;     %BetaPS
    param(:,79)=0.002;     %BetaPL
    param(:,3)=0.9;        %VmaxS
    param(:,5)=0.05;        %KNH4S
    param(:,16)=3.0;       %KNH4L
    param(:,4)=0.6;          %KNO3S  default = 1
    param(:,5)=0.05;        %KNH4S  default = 0.1
    param(:,15)=1;         %KNO3L  default = 3
    param(:,16)=1;       %KNH4L  default = 0.3
    param(:,12)=0.1;    %MorPS0   default is 0.0585
    param(:,56)=0.5;    %MorZP0
    param(:,42)=0.1;    %MorZL0
    param(:,30)=0.1;    %MorZS0
    param(:,62)=1;       %VP2N0
    param(:,64)=1;      %VP2D0
    param(:,66)=1;      %VD2N0
    param(:,32)=1.0;       %AlphaZS
    param(:,39)=0.0004;      %PS2ZLstar
    param(:,34)=0.01;       %GrmaxLPS
    param(:,28)=0.1;       %lamS   default = 1.4 (lower decreases grazing at low biomass)
    param(:,29)=0.04;      %PS2ZSstar   default = 0.04 (threshold)
    param(:,34)=0.001;       %GrmaxLPS   default = 0.1
    param(:,74)=4;       %setVP
    param(:,75)=4;       %setVO
elseif Trial==56
    Kzmixed=10^-4;
    param(:,76)=0.001;     %AlphaPS
    param(:,77)=0.008;     %AlphaPL
    param(:,78)=0.0002;     %BetaPS
    param(:,79)=0.002;     %BetaPL
    param(:,3)=0.9;        %VmaxS
    param(:,5)=0.05;        %KNH4S
    param(:,16)=3.0;       %KNH4L
    param(:,4)=0.3;          %KNO3S  default = 1
    param(:,5)=0.05;        %KNH4S  default = 0.1
    param(:,15)=1;         %KNO3L  default = 3
    param(:,16)=1;       %KNH4L  default = 0.3
    param(:,12)=0.1;    %MorPS0   default is 0.0585
    param(:,56)=0.5;    %MorZP0
    param(:,42)=0.1;    %MorZL0
    param(:,30)=0.1;    %MorZS0
    param(:,62)=2;       %VP2N0
    param(:,64)=2;      %VP2D0
    param(:,66)=2;      %VD2N0
    param(:,32)=1.0;       %AlphaZS
    param(:,39)=0.0004;      %PS2ZLstar
    param(:,34)=0.01;       %GrmaxLPS
    param(:,28)=0.1;       %lamS   default = 1.4 (lower decreases grazing at low biomass)
    param(:,29)=0.04;      %PS2ZSstar   default = 0.04 (threshold)
    param(:,34)=0.001;       %GrmaxLPS   default = 0.1
    param(:,74)=4;       %setVP
    param(:,75)=4;       %setVO
elseif Trial==57
    Kzmixed=10^-4;
    param(:,76)=0.001;     %AlphaPS
    param(:,77)=0.008;     %AlphaPL
    param(:,78)=0.0002;     %BetaPS
    param(:,79)=0.002;     %BetaPL
    param(:,3)=0.9;        %VmaxS
    param(:,5)=0.05;        %KNH4S
    param(:,16)=3.0;       %KNH4L
    param(:,4)=0.6;          %KNO3S  default = 1
    param(:,5)=0.05;        %KNH4S  default = 0.1
    param(:,15)=1;         %KNO3L  default = 3
    param(:,16)=1;       %KNH4L  default = 0.3
    param(:,12)=0.1;    %MorPS0   default is 0.0585
    param(:,56)=0.5;    %MorZP0
    param(:,42)=0.1;    %MorZL0
    param(:,30)=0.1;    %MorZS0
    param(:,62)=1;       %VP2N0
    param(:,64)=1;      %VP2D0
    param(:,66)=1;      %VD2N0
    param(:,32)=1.0;       %AlphaZS
    param(:,39)=0.0004;      %PS2ZLstar
    param(:,34)=0.01;       %GrmaxLPS
    param(:,28)=0.1;       %lamS   default = 1.4 (lower decreases grazing at low biomass)
    param(:,29)=0.04;      %PS2ZSstar   default = 0.04 (threshold)
    param(:,34)=0.001;       %GrmaxLPS   default = 0.1
    param(:,74)=4;       %setVP
    param(:,75)=4;       %setVO
    param(:,60)=0.003;      %Nit0  default is 0.03
elseif Trial==58
    Kzmixed=10^-4;
    param(:,76)=0.001;     %AlphaPS
    param(:,77)=0.008;     %AlphaPL
    param(:,78)=0.0002;     %BetaPS
    param(:,79)=0.002;     %BetaPL
    param(:,3)=0.9;        %VmaxS
    param(:,5)=0.05;        %KNH4S
    param(:,16)=3.0;       %KNH4L
    param(:,4)=0.6;          %KNO3S  default = 1
    param(:,5)=0.05;        %KNH4S  default = 0.1
    param(:,15)=1;         %KNO3L  default = 3
    param(:,16)=1;       %KNH4L  default = 0.3
    param(:,12)=0.01;    %MorPS0   default is 0.0585
    param(:,56)=0.5;    %MorZP0
    param(:,42)=0.1;    %MorZL0
    param(:,30)=0.1;    %MorZS0
    param(:,62)=1;       %VP2N0
    param(:,64)=1;      %VP2D0
    param(:,66)=1;      %VD2N0
    param(:,32)=1.0;       %AlphaZS     assimilation efficiency - default = 0.7
    param(:,33)=0.2;       %BetaZS      GGE default = 0.3
    param(:,39)=0.0004;      %PS2ZLstar
    param(:,34)=0.01;       %GrmaxLPS
    param(:,28)=0.1;       %lamS   default = 1.4 (lower decreases grazing at low biomass)
    param(:,29)=0.04;      %PS2ZSstar   default = 0.04 (threshold)
    param(:,34)=0.001;       %GrmaxLPS   default = 0.1
    param(:,74)=4;       %setVP
    param(:,75)=4;       %setVO
    param(:,60)=0.003;      %Nit0  default is 0.03
elseif Trial==59
    Kzmixed=10^-4;
    param(:,76)=0.001;     %AlphaPS
    param(:,77)=0.008;     %AlphaPL
    param(:,78)=0.0002;     %BetaPS
    param(:,79)=0.002;     %BetaPL
    param(:,3)=0.9;        %VmaxS
    param(:,5)=0.05;        %KNH4S
    param(:,16)=3.0;       %KNH4L
    param(:,4)=0.6;          %KNO3S  default = 1
    param(:,5)=0.05;        %KNH4S  default = 0.1
    param(:,15)=1;         %KNO3L  default = 3
    param(:,16)=1;       %KNH4L  default = 0.3
    param(:,12)=0.01;    %MorPS0   default is 0.0585
    param(:,56)=0.5;    %MorZP0
    param(:,42)=0.3;    %MorZL0
    param(:,30)=0.3;    %MorZS0
    param(:,62)=3;       %VP2N0
    param(:,64)=3;      %VP2D0
    param(:,66)=3;      %VD2N0
    param(:,32)=1.0;       %AlphaZS     assimilation efficiency - default = 0.7
    param(:,33)=0.2;       %BetaZS      GGE default = 0.3
    param(:,39)=0.04;      %PS2ZLstar
    param(:,34)=0.01;       %GrmaxLPS
    param(:,28)=0.1;       %lamS   default = 1.4 (lower decreases grazing at low biomass)
    param(:,29)=0.04;      %PS2ZSstar   default = 0.04 (threshold)
    param(:,34)=0.001;       %GrmaxLPS   default = 0.1
    param(:,74)=4;       %setVP
    param(:,75)=4;       %setVO
    param(:,60)=0.003;      %Nit0  default is 0.03
elseif Trial==60
    Kzmixed=10^-4;
    param(:,76)=0.001;     %AlphaPS
    param(:,77)=0.008;     %AlphaPL
    param(:,78)=0.0002;     %BetaPS
    param(:,79)=0.002;     %BetaPL
    param(:,3)=0.9;        %VmaxS
    param(:,5)=0.05;        %KNH4S
    param(:,16)=3.0;       %KNH4L
    param(:,4)=0.6;          %KNO3S  default = 1
    param(:,5)=0.05;        %KNH4S  default = 0.1
    param(:,15)=1;         %KNO3L  default = 3
    param(:,16)=1;       %KNH4L  default = 0.3
    param(:,12)=0.01;    %MorPS0   default is 0.0585
    param(:,56)=0.5;    %MorZP0
    param(:,42)=0.3;    %MorZL0
    param(:,30)=0.3;    %MorZS0
    param(:,62)=3;       %VP2N0
    param(:,64)=3;      %VP2D0
    param(:,66)=3;      %VD2N0
    param(:,32)=1.0;       %AlphaZS     assimilation efficiency - default = 0.7
    param(:,33)=0.2;       %BetaZS      GGE default = 0.3
    param(:,39)=0.04;      %PS2ZLstar
    param(:,34)=0.01;       %GrmaxLPS
    param(:,28)=0.1;       %lamS   default = 1.4 (lower decreases grazing at low biomass)
    param(:,29)=0.04;      %PS2ZSstar   default = 0.04 (threshold)
    param(:,34)=0.001;       %GrmaxLPS   default = 0.1
    param(:,74)=4;       %setVP
    param(:,75)=4;       %setVO
    param(:,60)=0.003;      %Nit0  default is 0.03
elseif Trial==61
    Kzmixed=10^-4;
    param(:,76)=0.001;     %AlphaPS
    param(:,77)=0.008;     %AlphaPL
    param(:,78)=0.0002;     %BetaPS
    param(:,79)=0.002;     %BetaPL
    param(:,3)=0.9;        %VmaxS
    param(:,5)=0.05;        %KNH4S
    param(:,16)=3.0;       %KNH4L
    param(:,4)=0.6;          %KNO3S  default = 1
    param(:,5)=0.05;        %KNH4S  default = 0.1
    param(:,15)=1;         %KNO3L  default = 3
    param(:,16)=1;       %KNH4L  default = 0.3
    param(:,12)=0.01;    %MorPS0   default is 0.0585
    param(:,56)=0.5;    %MorZP0
    param(:,42)=0.3;    %MorZL0
    param(:,30)=0.3;    %MorZS0
    param(:,62)=3;       %VP2N0
    param(:,64)=3;      %VP2D0
    param(:,66)=3;      %VD2N0
    param(:,32)=1.0;       %AlphaZS     assimilation efficiency - default = 0.7
    param(:,33)=0.2;       %BetaZS      GGE default = 0.3
    param(:,39)=10;      %PS2ZLstar   default = 0.04 (threshold)
    param(:,34)=0.01;       %GrmaxLPS
    param(:,28)=0.1;       %lamS   default = 1.4 (lower decreases grazing at low biomass)
    param(:,29)=0.04;      %PS2ZSstar   default = 0.04 (threshold)
    param(:,34)=0.001;       %GrmaxLPS   default = 0.1
    param(:,74)=4;       %setVP
    param(:,75)=4;       %setVO
    param(:,60)=0.003;      %Nit0  default is 0.03
elseif Trial==62
    Kzmixed=10^-5;
    param(:,76)=0.001;     %AlphaPS
    param(:,77)=0.008;     %AlphaPL
    param(:,78)=0.0002;     %BetaPS
    param(:,79)=0.002;     %BetaPL
    param(:,3)=0.9;        %VmaxS
    param(:,5)=0.05;        %KNH4S
    param(:,16)=3.0;       %KNH4L
    param(:,4)=0.6;          %KNO3S  default = 1
    param(:,5)=0.05;        %KNH4S  default = 0.1
    param(:,15)=1;         %KNO3L  default = 3
    param(:,16)=1;       %KNH4L  default = 0.3
    param(:,12)=0.01;    %MorPS0   default is 0.0585
    param(:,56)=0.5;    %MorZP0
    param(:,42)=0.3;    %MorZL0
    param(:,30)=0.3;    %MorZS0
    param(:,62)=3;       %VP2N0
    param(:,64)=3;      %VP2D0
    param(:,66)=3;      %VD2N0
    param(:,32)=1.0;       %AlphaZS     assimilation efficiency - default = 0.7
    param(:,33)=0.2;       %BetaZS      GGE default = 0.3
    param(:,39)=10;      %PS2ZLstar   default = 0.04 (threshold)
    param(:,34)=0.01;       %GrmaxLPS
    param(:,28)=0.1;       %lamS   default = 1.4 (lower decreases grazing at low biomass)
    param(:,29)=0.04;      %PS2ZSstar   default = 0.04 (threshold)
    param(:,34)=0.001;       %GrmaxLPS   default = 0.1
    param(:,74)=4;       %setVP
    param(:,75)=4;       %setVO
    param(:,60)=0.003;      %Nit0  default is 0.03
elseif Trial==63
    Kzmixed=10^-5;
    param(:,76)=0.001;     %AlphaPS
    param(:,77)=0.008;     %AlphaPL
    param(:,78)=0.0002;     %BetaPS
    param(:,79)=0.002;     %BetaPL
    param(:,3)=0.9;        %VmaxS
    param(:,5)=0.05;        %KNH4S
    param(:,16)=3.0;       %KNH4L
    param(:,4)=0.6;          %KNO3S  default = 1
    param(:,5)=0.05;        %KNH4S  default = 0.1
    param(:,15)=1;         %KNO3L  default = 3
    param(:,16)=1;       %KNH4L  default = 0.3
    param(:,12)=0.01;    %MorPS0   default is 0.0585
    param(:,56)=0.5;    %MorZP0
    param(:,42)=0.3;    %MorZL0
    param(:,30)=0.3;    %MorZS0
    param(:,62)=3;       %VP2N0
    param(:,64)=3;      %VP2D0
    param(:,66)=3;      %VD2N0
    param(:,32)=1.0;       %AlphaZS     assimilation efficiency - default = 0.7
    param(:,33)=0.2;       %BetaZS      GGE default = 0.3
    param(:,39)=10;      %PS2ZLstar   default = 0.04 (threshold)
    param(:,34)=0.01;       %GrmaxLPS
    param(:,28)=0.1;       %lamS   default = 1.4 (lower decreases grazing at low biomass)
    param(:,29)=0.005;      %PS2ZSstar   default = 0.04 (threshold)
    param(:,34)=0.001;       %GrmaxLPS   default = 0.1
    param(:,74)=4;       %setVP
    param(:,75)=4;       %setVO
    param(:,60)=0.003;      %Nit0  default is 0.03
elseif Trial==64
    Kzmixed=10^-5;
    param(:,76)=0.001;     %AlphaPS
    param(:,77)=0.008;     %AlphaPL
    param(:,78)=0.0002;     %BetaPS
    param(:,79)=0.002;     %BetaPL
    param(:,3)=0.9;        %VmaxS
    param(:,5)=0.05;        %KNH4S
    param(:,16)=3.0;       %KNH4L
    param(:,4)=0.6;          %KNO3S  default = 1
    param(:,5)=0.05;        %KNH4S  default = 0.1
    param(:,15)=1;         %KNO3L  default = 3
    param(:,16)=1;       %KNH4L  default = 0.3
    param(:,12)=0.01;    %MorPS0   default is 0.0585
    param(:,56)=0.5;    %MorZP0
    param(:,42)=0.3;    %MorZL0
    param(:,30)=0.3;    %MorZS0
    param(:,62)=3;       %VP2N0
    param(:,64)=3;      %VP2D0
    param(:,66)=3;      %VD2N0
    param(:,32)=1.0;       %AlphaZS     assimilation efficiency - default = 0.7
    param(:,33)=0.2;       %BetaZS      GGE default = 0.3
    param(:,39)=10;      %PS2ZLstar   default = 0.04 (threshold)
    param(:,34)=0.01;       %GrmaxLPS
    param(:,28)=1.4;       %lamS   default = 1.4 (lower decreases grazing at low biomass)
    param(:,29)=0.005;      %PS2ZSstar   default = 0.04 (threshold)
    param(:,34)=0.001;       %GrmaxLPS   default = 0.1
    param(:,74)=4;       %setVP
    param(:,75)=4;       %setVO
    param(:,60)=0.003;      %Nit0  default is 0.03
elseif Trial==65
    Kzmixed=10^-5;
    param(:,76)=0.001;     %AlphaPS
    param(:,77)=0.008;     %AlphaPL
    param(:,78)=0.0002;     %BetaPS
    param(:,79)=0.005;     %BetaPL
    param(:,3)=0.9;        %VmaxS
    param(:,5)=0.05;        %KNH4S
    param(:,16)=3.0;       %KNH4L
    param(:,4)=0.6;          %KNO3S  default = 1
    param(:,5)=0.05;        %KNH4S  default = 0.1
    param(:,15)=1;         %KNO3L  default = 3
    param(:,16)=1;       %KNH4L  default = 0.3
    param(:,12)=0.3;    %MorPS0   default is 0.0585
    param(:,56)=0.5;    %MorZP0
    param(:,42)=0.3;    %MorZL0
    param(:,30)=0.3;    %MorZS0
    param(:,62)=3;       %VP2N0
    param(:,64)=3;      %VP2D0
    param(:,66)=3;      %VD2N0
    param(:,32)=1.0;       %AlphaZS     assimilation efficiency - default = 0.7
    param(:,33)=0.2;       %BetaZS      GGE default = 0.3
    param(:,39)=10;      %PS2ZLstar   default = 0.04 (threshold)
    param(:,34)=0.01;       %GrmaxLPS
    param(:,28)=1.4;       %lamS   default = 1.4 (lower decreases grazing at low biomass)
    param(:,29)=0.5;      %PS2ZSstar   default = 0.04 (threshold)
    param(:,34)=0.001;       %GrmaxLPS   default = 0.1
    param(:,74)=4;       %setVP
    param(:,75)=4;       %setVO
    param(:,60)=0.003;      %Nit0  default is 0.03
elseif Trial==66
    param(:,62)=3;       %VP2N0
    param(:,64)=3;      %VP2D0
    param(:,66)=3;      %VD2N0
    param(:,32)=1.0;       %AlphaZS     assimilation efficiency - default = 0.7
    param(:,33)=0.2;       %BetaZS      GGE default = 0.3
    param(:,39)=10;      %PS2ZLstar   default = 0.04 (threshold)
elseif Trial==67
    param(:,62)=3;       %VP2N0
    param(:,64)=3;      %VP2D0
    param(:,66)=3;      %VD2N0
    param(:,32)=1.0;       %AlphaZS     assimilation efficiency - default = 0.7
    param(:,33)=0.2;       %BetaZS      GGE default = 0.3
    param(:,39)=10;      %PS2ZLstar   default = 0.04 (threshold)
    param(:,60)=0.3;      %Nit0  default is 0.03
elseif Trial==68
    param(:,62)=3;       %VP2N0
    param(:,64)=3;      %VP2D0
    param(:,66)=3;      %VD2N0
    param(:,32)=1.0;       %AlphaZS     assimilation efficiency - default = 0.7
    param(:,33)=0.2;       %BetaZS      GGE default = 0.3
    param(:,39)=10;      %PS2ZLstar   default = 0.04 (threshold)
    param(:,74)=4;       %setVP
    param(:,75)=4;       %setVO
elseif Trial==69
    param(:,62)=3;       %VP2N0
    param(:,64)=3;      %VP2D0
    param(:,66)=3;      %VD2N0
    param(:,32)=1.0;       %AlphaZS     assimilation efficiency - default = 0.7
    param(:,33)=0.2;       %BetaZS      GGE default = 0.3
    param(:,39)=10;      %PS2ZLstar   default = 0.04 (threshold)
    param(:,9)=0.0;       %ResPS0    phytoplankton respiration - default = 0.03 
elseif Trial==69
    param(:,62)=3;       %VP2N0
    param(:,64)=3;      %VP2D0
    param(:,66)=3;      %VD2N0
    param(:,32)=1.0;       %AlphaZS     assimilation efficiency - default = 0.7
    param(:,33)=0.2;       %BetaZS      GGE default = 0.3
    param(:,39)=10;      %PS2ZLstar   default = 0.04 (threshold)
    Kzmixed=10^-4;
elseif Trial==69
    param(:,62)=3;       %VP2N0
    param(:,64)=3;      %VP2D0
    param(:,66)=3;      %VD2N0
    param(:,32)=1.0;       %AlphaZS     assimilation efficiency - default = 0.7
    param(:,33)=0.2;       %BetaZS      GGE default = 0.3
    param(:,39)=10;      %PS2ZLstar   default = 0.04 (threshold)
    Kzmixed=10^-4;
    param(:,60)=0.3;      %Nit0  default is 0.03
elseif Trial==70
    param(:,62)=3;       %VP2N0
    param(:,64)=3;      %VP2D0
    param(:,66)=3;      %VD2N0
    param(:,32)=1.0;       %AlphaZS     assimilation efficiency - default = 0.7
    param(:,33)=0.2;       %BetaZS      GGE default = 0.3
    param(:,39)=10;      %PS2ZLstar   default = 0.04 (threshold)
    Kzmixed=10^-4;
    param(:,60)=0.3;      %Nit0  default is 0.03
    param(:,5)=0.3;        %KNH4S - default = 0.1
    param(:,16)=3.0;       %KNH4L - default - 0.3
    param(:,4)=0.6;          %KNO3S  default = 1
    param(:,15)=3;         %KNO3L - default = 3
elseif Trial==71
    param(:,62)=3;       %VP2N0
    param(:,64)=3;      %VP2D0
    param(:,66)=3;      %VD2N0
    param(:,32)=1.0;       %AlphaZS     assimilation efficiency - default = 0.7
    param(:,33)=0.2;       %BetaZS      GGE default = 0.3
    param(:,39)=10;      %PS2ZLstar   default = 0.04 (threshold)
    Kzmixed=10^-4;
    param(:,60)=0.3;      %Nit0  default is 0.03
    param(:,9)=0.0;       %ResPS0 default is 0.03
    param(:,21)=0.0;      %ResPL0 default is 0.03
elseif Trial==72
    param(:,62)=3;       %VP2N0
    param(:,64)=3;      %VP2D0
    param(:,66)=3;      %VD2N0
    param(:,32)=1.0;       %AlphaZS     assimilation efficiency - default = 0.7
    param(:,33)=0.2;       %BetaZS      GGE default = 0.3
    param(:,39)=10;      %PS2ZLstar   default = 0.04 (threshold)
    param(:,60)=0.3;      %Nit0  default is 0.03
    param(:,9)=0.0;       %ResPS0 default is 0.03
    param(:,21)=0.0;      %ResPL0 default is 0.03
elseif Trial==73
    param(:,32)=1.0;       %AlphaZS     assimilation efficiency - default = 0.7
    param(:,33)=0.2;       %BetaZS      GGE default = 0.3
    param(:,39)=10;      %PS2ZLstar   default = 0.04 (threshold)
    param(:,60)=0.3;      %Nit0  default is 0.03
    param(:,9)=0.0;       %ResPS0 default is 0.03
    param(:,21)=0.0;      %ResPL0 default is 0.03
elseif Trial==74
    param(:,62)=3;       %VP2N0
    param(:,64)=3;      %VP2D0
    param(:,66)=3;      %VD2N0
    param(:,32)=1.0;       %AlphaZS     assimilation efficiency - default = 0.7
    param(:,33)=0.2;       %BetaZS      GGE default = 0.3
    param(:,39)=10;      %PS2ZLstar   default = 0.04 (threshold)
    param(:,9)=0.0;       %ResPS0 default is 0.03
    param(:,21)=0.0;      %ResPL0 default is 0.03
elseif Trial==75
    param(:,32)=1.0;       %AlphaZS     assimilation efficiency - default = 0.7
    param(:,33)=0.2;       %BetaZS      GGE default = 0.3
    param(:,39)=10;      %PS2ZLstar   default = 0.04 (threshold)
    param(:,9)=0.0;       %ResPS0 default is 0.03
    param(:,21)=0.0;      %ResPL0 default is 0.03
elseif Trial==76
    param(:,9)=0.0;       %ResPS0 default is 0.03
    param(:,21)=0.0;      %ResPL0 default is 0.03
elseif Trial==77
    Kzmixed = 10^-4;
    param(:,9)=0.0;       %ResPS0 default is 0.03
    param(:,21)=0.0;      %ResPL0 default is 0.03
elseif Trial==78
    Kzmixed = 10^-4;
    param(:,9)=0.03;       %ResPS0 default is 0.03
    param(:,21)=0.03;      %ResPL0 default is 0.03
    param(:,76)=0.001;     %AlphaPS
    param(:,77)=0.008;     %AlphaPL
    param(:,78)=0.0002;     %BetaPS
    param(:,79)=0.002;     %BetaPL
    param(:,3)=0.9;        %VmaxS
    param(:,4)=0.1;          %KNO3S  default = 1
    param(:,5)=0.1;        %KNH4S  default = 0.1
    param(:,15)=1;         %KNO3L  default = 3
    param(:,16)=1;       %KNH4L  default = 0.3
elseif Trial==79
    Kzmixed = 10^-4;
    param(:,9)=0.03;       %ResPS0 default is 0.03
    param(:,21)=0.03;      %ResPL0 default is 0.03
    param(:,76)=0.001;     %AlphaPS
    param(:,77)=0.008;     %AlphaPL
    param(:,78)=0.0002;     %BetaPS
    param(:,79)=0.002;     %BetaPL
    param(:,3)=0.9;        %VmaxS
    param(:,4)=0.1;          %KNO3S  default = 1
    param(:,5)=0.1;        %KNH4S  default = 0.1
    param(:,15)=1;         %KNO3L  default = 3
    param(:,16)=1;       %KNH4L  default = 0.3
    param(:,60)=0.3;      %Nit0  default is 0.03
elseif Trial==80
    param(:,9)=0.03;       %ResPS0 default is 0.03
    param(:,21)=0.03;      %ResPL0 default is 0.03
    param(:,76)=0.001;     %AlphaPS
    param(:,77)=0.008;     %AlphaPL
    param(:,78)=0.0002;     %BetaPS
    param(:,79)=0.002;     %BetaPL
    param(:,3)=0.9;        %VmaxS
    param(:,4)=0.1;          %KNO3S  default = 1
    param(:,5)=0.1;        %KNH4S  default = 0.1
    param(:,15)=1;         %KNO3L  default = 3
    param(:,16)=1;       %KNH4L  default = 0.3
    param(:,60)=0.3;      %Nit0  default is 0.03
elseif Trial==81
    param(:,9)=0.03;       %ResPS0 default is 0.03
    param(:,21)=0.03;      %ResPL0 default is 0.03
    param(:,76)=0.001;     %AlphaPS
    param(:,77)=0.008;     %AlphaPL
    param(:,78)=0.0002;     %BetaPS
    param(:,79)=0.002;     %BetaPL
    param(:,3)=0.9;        %VmaxS
    param(:,4)=0.1;          %KNO3S  default = 1
    param(:,5)=0.1;        %KNH4S  default = 0.1
    param(:,15)=1;         %KNO3L  default = 3
    param(:,16)=1;       %KNH4L  default = 0.3
    param(:,60)=0.3;      %Nit0  default is 0.03
    param(:,62)=3;       %VP2N0
    param(:,64)=3;      %VP2D0
    param(:,66)=3;      %VD2N0
elseif Trial==82
    param(:,9)=0.03;       %ResPS0 default is 0.03
    param(:,21)=0.03;      %ResPL0 default is 0.03
    param(:,76)=0.001;     %AlphaPS
    param(:,77)=0.008;     %AlphaPL
    param(:,78)=0.0002;     %BetaPS
    param(:,79)=0.002;     %BetaPL
    param(:,3)=0.9;        %VmaxS
    param(:,4)=0.1;          %KNO3S  default = 1
    param(:,5)=0.1;        %KNH4S  default = 0.1
    param(:,15)=1;         %KNO3L  default = 3
    param(:,16)=1;       %KNH4L  default = 0.3
    param(:,60)=0.3;      %Nit0  default is 0.03
    param(:,62)=3;       %VP2N0
    param(:,64)=3;      %VP2D0
    param(:,66)=3;      %VD2N0
    param(:,32)=1.0;       %AlphaZS     assimilation efficiency - default = 0.7
    param(:,33)=0.2;       %BetaZS      GGE default = 0.3
    param(:,39)=10;      %PS2ZLstar   default = 0.04 (threshold)
elseif Trial==83
    Kzmixed=10^-5;
    param(:,9)=0.03;       %ResPS0 default is 0.03
    param(:,21)=0.03;      %ResPL0 default is 0.03
    param(:,76)=0.001;     %AlphaPS
    param(:,77)=0.008;     %AlphaPL
    param(:,78)=0.0002;     %BetaPS
    param(:,79)=0.002;     %BetaPL
    param(:,3)=0.9;        %VmaxS
    param(:,4)=0.1;          %KNO3S  default = 1
    param(:,5)=0.1;        %KNH4S  default = 0.1
    param(:,15)=1;         %KNO3L  default = 3
    param(:,16)=1;       %KNH4L  default = 0.3
    param(:,60)=0.3;      %Nit0  default is 0.03
    param(:,62)=3;       %VP2N0
    param(:,64)=3;      %VP2D0
    param(:,66)=3;      %VD2N0
    param(:,32)=1.0;       %AlphaZS     assimilation efficiency - default = 0.7
    param(:,33)=0.2;       %BetaZS      GGE default = 0.3
    param(:,39)=10;      %PS2ZLstar   default = 0.04 (threshold)
elseif Trial==84
    Kzmixed=10^-5;
    param(:,9)=0.03;       %ResPS0 default is 0.03
    param(:,21)=0.03;      %ResPL0 default is 0.03
    param(:,76)=0.001;     %AlphaPS
    param(:,77)=0.008;     %AlphaPL
    param(:,78)=0.0002;     %BetaPS
    param(:,79)=0.002;     %BetaPL
    param(:,3)=0.9;        %VmaxS
    param(:,4)=0.03;          %KNO3S  default = 1
    param(:,5)=0.03;        %KNH4S  default = 0.1
    param(:,15)=1;         %KNO3L  default = 3
    param(:,16)=1;       %KNH4L  default = 0.3
    param(:,60)=0.5;      %Nit0  default is 0.03
    param(:,62)=3;       %VP2N0
    param(:,64)=3;      %VP2D0
    param(:,66)=3;      %VD2N0
    param(:,32)=1.0;       %AlphaZS     assimilation efficiency - default = 0.7
    param(:,33)=0.2;       %BetaZS      GGE default = 0.3
    param(:,39)=10;      %PS2ZLstar   default = 0.04 (threshold)
elseif Trial==85
    Kzmixed=10^-5;
    param(:,9)=0.03;       %ResPS0 default is 0.03
    param(:,21)=0.03;      %ResPL0 default is 0.03
    param(:,76)=0.001;     %AlphaPS
    param(:,77)=0.008;     %AlphaPL
    param(:,78)=0.0002;     %BetaPS
    param(:,79)=0.002;     %BetaPL
    param(:,3)=0.9;        %VmaxS
    param(:,4)=0.01;          %KNO3S  default = 1
    param(:,5)=0.03;        %KNH4S  default = 0.1
    param(:,15)=1;         %KNO3L  default = 3
    param(:,16)=1;       %KNH4L  default = 0.3
    param(:,60)=0.5;      %Nit0  default is 0.03
    param(:,62)=3;       %VP2N0
    param(:,64)=3;      %VP2D0
    param(:,66)=3;      %VD2N0
    param(:,32)=1.0;       %AlphaZS     assimilation efficiency - default = 0.7
    param(:,33)=0.2;       %BetaZS      GGE default = 0.3
    param(:,39)=10;      %PS2ZLstar   default = 0.04 (threshold)
elseif Trial==86
    Kzmixed=10^-5;
    param(:,9)=0.03;       %ResPS0 default is 0.03
    param(:,21)=0.03;      %ResPL0 default is 0.03
    param(:,76)=0.001;     %AlphaPS
    param(:,77)=0.008;     %AlphaPL
    param(:,78)=0.0002;     %BetaPS
    param(:,79)=0.002;     %BetaPL
    param(:,3)=0.9;        %VmaxS
    param(:,4)=0.01;          %KNO3S  default = 1
    param(:,5)=0.03;        %KNH4S  default = 0.1
    param(:,15)=1;         %KNO3L  default = 3
    param(:,16)=1;       %KNH4L  default = 0.3
    param(:,60)=0;      %Nit0  default is 0.03
    param(:,62)=3;       %VP2N0
    param(:,64)=3;      %VP2D0
    param(:,66)=3;      %VD2N0
    param(:,32)=1.0;       %AlphaZS     assimilation efficiency - default = 0.7
    param(:,33)=0.2;       %BetaZS      GGE default = 0.3
    param(:,39)=10;      %PS2ZLstar   default = 0.04 (threshold)
elseif Trial==87
    Kzmixed=10^-5;
    param(:,9)=0.03;       %ResPS0 default is 0.03
    param(:,21)=0.03;      %ResPL0 default is 0.03
    param(:,76)=0.008;     %AlphaPS
    param(:,77)=0.001;     %AlphaPL
    param(:,78)=0.002;     %BetaPS
    param(:,79)=0.0002;     %BetaPL
    param(:,3)=0.9;        %VmaxS
    param(:,4)=0.1;          %KNO3S  default = 1
    param(:,5)=0.01;        %KNH4S  default = 0.1
    param(:,15)=0.1;         %KNO3L  default = 3
    param(:,16)=1;       %KNH4L  default = 0.3
    param(:,18)=0.5;       %PsiL = ammonium inhibition coeff (smaller is less inhibition) default is 4.6
    param(:,60)=0.5;      %Nit0  default is 0.03
    param(:,62)=3;       %VP2N0
    param(:,64)=3;      %VP2D0
    param(:,66)=3;      %VD2N0
    param(:,32)=1.0;       %AlphaZS     assimilation efficiency - default = 0.7
    param(:,33)=0.2;       %BetaZS      GGE default = 0.3
    param(:,39)=10;      %PS2ZLstar   default = 0.04 (threshold)
elseif Trial==88
    Kzmixed=10^-5;
    param(:,9)=0.01;       %ResPS0 default is 0.03
    param(:,21)=0.01;      %ResPL0 default is 0.03
    param(:,76)=0.001;     %AlphaPS
    param(:,77)=0.008;     %AlphaPL
    param(:,78)=0.0002;     %BetaPS
    param(:,79)=0.002;     %BetaPL
    param(:,3)=0.9;        %VmaxS
    param(:,4)=0.01;          %KNO3S  default = 1
    param(:,5)=0.03;        %KNH4S  default = 0.1
    param(:,15)=1;         %KNO3L  default = 3
    param(:,16)=1;       %KNH4L  default = 0.3
    param(:,60)=0;      %Nit0  default is 0.03
    param(:,62)=3;       %VP2N0
    param(:,64)=3;      %VP2D0
    param(:,66)=3;      %VD2N0
    param(:,32)=1.0;       %AlphaZS     assimilation efficiency - default = 0.7
    param(:,33)=0.2;       %BetaZS      GGE default = 0.3
    param(:,39)=10;      %PS2ZLstar   default = 0.04 (threshold)
elseif Trial==89
    Kzmixed=10^-5;
    param(:,9)=0.00;       %ResPS0 default is 0.03
    param(:,21)=0.00;      %ResPL0 default is 0.03
    param(:,76)=0.001;     %AlphaPS
    param(:,77)=0.008;     %AlphaPL
    param(:,78)=0.0002;     %BetaPS
    param(:,79)=0.002;     %BetaPL
    param(:,3)=0.9;        %VmaxS
    param(:,4)=0.01;          %KNO3S  default = 1
    param(:,5)=0.03;        %KNH4S  default = 0.1
    param(:,15)=1;         %KNO3L  default = 3
    param(:,16)=1;       %KNH4L  default = 0.3
    param(:,60)=0;      %Nit0  default is 0.03
    param(:,62)=3;       %VP2N0
    param(:,64)=3;      %VP2D0
    param(:,66)=3;      %VD2N0
    param(:,32)=1.0;       %AlphaZS     assimilation efficiency - default = 0.7
    param(:,33)=0.2;       %BetaZS      GGE default = 0.3
    param(:,39)=10;      %PS2ZLstar   default = 0.04 (threshold)
elseif Trial==90
    Kzmixed=10^-4;
    param(:,9)=0.03;       %ResPS0 default is 0.03
    param(:,21)=0.03;      %ResPL0 default is 0.03
    param(:,76)=0.001;     %AlphaPS
    param(:,77)=0.008;     %AlphaPL
    param(:,78)=0.0002;     %BetaPS
    param(:,79)=0.002;     %BetaPL
    param(:,3)=0.9;        %VmaxS
    param(:,4)=0.01;          %KNO3S  default = 1
    param(:,5)=0.03;        %KNH4S  default = 0.1
    param(:,15)=1;         %KNO3L  default = 3
    param(:,16)=1;       %KNH4L  default = 0.3
    param(:,60)=0;      %Nit0  default is 0.03
    param(:,62)=3;       %VP2N0
    param(:,64)=3;      %VP2D0
    param(:,66)=3;      %VD2N0
    param(:,32)=1.0;       %AlphaZS     assimilation efficiency - default = 0.7
    param(:,33)=0.2;       %BetaZS      GGE default = 0.3
    param(:,39)=10;      %PS2ZLstar   default = 0.04 (threshold)
elseif Trial==91
    Kzmixed=10^-5;
    param(:,9)=0.03;       %ResPS0 default is 0.03
    param(:,21)=0.03;      %ResPL0 default is 0.03
    param(:,76)=0.001;     %AlphaPS
    param(:,77)=0.008;     %AlphaPL
    param(:,78)=0.0002;     %BetaPS
    param(:,79)=0.002;     %BetaPL
    param(:,3)=0.9;        %VmaxS
    param(:,4)=0.01;          %KNO3S  default = 1
    param(:,5)=0.03;        %KNH4S  default = 0.1
    param(:,15)=1;         %KNO3L  default = 3
    param(:,16)=1;       %KNH4L  default = 0.3
    param(:,60)=0.5;      %Nit0  default is 0.03
    param(:,62)=3;       %VP2N0
    param(:,64)=3;      %VP2D0
    param(:,66)=3;      %VD2N0
    param(:,32)=1.0;       %AlphaZS     assimilation efficiency - default = 0.7
    param(:,33)=0.2;       %BetaZS      GGE default = 0.3
    param(:,39)=10;      %PS2ZLstar   default = 0.04 (threshold)
    param(:,74)=4;       %setVP   sinking speed, default = 40
    param(:,75)=4;       %setVO   sinking speed, default = 40
elseif Trial==92
    Kzmixed=10^-5;
    param(:,9)=0.03;       %ResPS0 default is 0.03
    param(:,21)=0.03;      %ResPL0 default is 0.03
    param(:,76)=0.001;     %AlphaPS
    param(:,77)=0.008;     %AlphaPL
    param(:,78)=0.0002;     %BetaPS
    param(:,79)=0.002;     %BetaPL
    param(:,3)=0.9;        %VmaxS
    param(:,4)=0.01;          %KNO3S  default = 1
    param(:,5)=0.03;        %KNH4S  default = 0.1
    param(:,15)=1;         %KNO3L  default = 3
    param(:,16)=1;       %KNH4L  default = 0.3
    param(:,60)=0.03;      %Nit0  default is 0.03
    param(:,62)=3;       %VP2N0
    param(:,64)=3;      %VP2D0
    param(:,66)=3;      %VD2N0
    param(:,32)=1.0;       %AlphaZS     assimilation efficiency - default = 0.7
    param(:,33)=0.2;       %BetaZS      GGE default = 0.3
    param(:,39)=10;      %PS2ZLstar   default = 0.04 (threshold)
    param(:,74)=4;       %setVP   sinking speed, default = 40
    param(:,75)=4;       %setVO   sinking speed, default = 40
elseif Trial==93
    Kzmixed=10^-5;
    param(:,9)=0.03;       %ResPS0 default is 0.03
    param(:,21)=0.03;      %ResPL0 default is 0.03
    param(:,76)=0.001;     %AlphaPS
    param(:,77)=0.008;     %AlphaPL
    param(:,78)=0.0002;     %BetaPS
    param(:,79)=0.002;     %BetaPL
    param(:,3)=0.9;        %VmaxS
    param(:,4)=0.03;          %KNO3S  default = 1
    param(:,5)=0.03;        %KNH4S  default = 0.1
    param(:,15)=1;         %KNO3L  default = 3
    param(:,16)=1;       %KNH4L  default = 0.3
    param(:,60)=0.03;      %Nit0  default is 0.03
    param(:,62)=3;       %VP2N0
    param(:,64)=3;      %VP2D0
    param(:,66)=3;      %VD2N0
    param(:,32)=1.0;       %AlphaZS     assimilation efficiency - default = 0.7
    param(:,33)=0.2;       %BetaZS      GGE default = 0.3
    param(:,39)=10;      %PS2ZLstar   default = 0.04 (threshold)
    param(:,74)=4;       %setVP   sinking speed, default = 40
    param(:,75)=4;       %setVO   sinking speed, default = 40
elseif Trial==94
    Kzmixed=10^-5;
    param(:,9)=0.03;       %ResPS0 default is 0.03
    param(:,21)=0.03;      %ResPL0 default is 0.03
    param(:,76)=0.001;     %AlphaPS
    param(:,77)=0.008;     %AlphaPL
    param(:,78)=0.0002;     %BetaPS
    param(:,79)=0.002;     %BetaPL
    param(:,3)=0.9;        %VmaxS
    param(:,4)=0.03;          %KNO3S  default = 1
    param(:,5)=0.03;        %KNH4S  default = 0.1
    param(:,15)=1;         %KNO3L  default = 3
    param(:,16)=1;       %KNH4L  default = 0.3
    param(:,60)=0.5;      %Nit0  default is 0.03
    param(:,62)=3;       %VP2N0
    param(:,64)=3;      %VP2D0
    param(:,66)=3;      %VD2N0
    param(:,32)=1.0;       %AlphaZS     assimilation efficiency - default = 0.7
    param(:,33)=0.2;       %BetaZS      GGE default = 0.3
    param(:,39)=10;      %PS2ZLstar   default = 0.04 (threshold)
    param(:,74)=4;       %setVP   sinking speed, default = 40
    param(:,75)=4;       %setVO   sinking speed, default = 40
elseif Trial==95
    Kzmixed=10^-5;
    param(:,9)=0.03;       %ResPS0 default is 0.03
    param(:,21)=0.03;      %ResPL0 default is 0.03
    param(:,76)=0.001;     %AlphaPS
    param(:,77)=0.008;     %AlphaPL
    param(:,78)=0.0002;     %BetaPS
    param(:,79)=0.002;     %BetaPL
    param(:,3)=0.9;        %VmaxS
    param(:,4)=0.1;          %KNO3S  default = 1
    param(:,5)=0.01;        %KNH4S  default = 0.1
    param(:,15)=1;         %KNO3L  default = 3
    param(:,16)=1;       %KNH4L  default = 0.3
    param(:,60)=0.03;      %Nit0  default is 0.03
    param(:,62)=3;       %VP2N0
    param(:,64)=3;      %VP2D0
    param(:,66)=3;      %VD2N0
    param(:,32)=1.0;       %AlphaZS     assimilation efficiency - default = 0.7
    param(:,33)=0.2;       %BetaZS      GGE default = 0.3
    param(:,39)=10;      %PS2ZLstar   default = 0.04 (threshold)
    param(:,74)=4;       %setVP   sinking speed, default = 40
    param(:,75)=4;       %setVO   sinking speed, default = 40
elseif Trial==96
    Kzmixed=10^-5;
    param(:,9)=0.0;       %ResPS0 default is 0.03
    param(:,21)=0.0;      %ResPL0 default is 0.03
    param(:,76)=0.001;     %AlphaPS
    param(:,77)=0.008;     %AlphaPL
    param(:,78)=0.0002;     %BetaPS
    param(:,79)=0.002;     %BetaPL
    param(:,3)=0.9;        %VmaxS
    param(:,4)=0.1;          %KNO3S  default = 1
    param(:,5)=0.01;        %KNH4S  default = 0.1
    param(:,15)=3;         %KNO3L  default = 3
    param(:,16)=0.3;       %KNH4L  default = 0.3
    param(:,60)=0.03;      %Nit0  default is 0.03
    param(:,62)=3;       %VP2N0
    param(:,64)=3;      %VP2D0
    param(:,66)=3;      %VD2N0
    param(:,32)=1.0;       %AlphaZS     assimilation efficiency - default = 0.7
    param(:,33)=0.2;       %BetaZS      GGE default = 0.3
    param(:,39)=10;      %PS2ZLstar   default = 0.04 (threshold)
    param(:,74)=4;       %setVP   sinking speed, default = 40
    param(:,75)=4;       %setVO   sinking speed, default = 40
elseif Trial==97
    Kzmixed=10^-3;
    Kzdeep = 10^-4;
    param(:,9)=0.03;       %ResPS0 default is 0.03
    param(:,21)=0.03;      %ResPL0 default is 0.03
    param(:,18)=15;       %PsiL = ammonium inhibition coeff (smaller is less inhibition) default is 4.6
    param(:,62)=1;       %VP2N0
    param(:,64)=1;      %VP2D0
    param(:,66)=1;      %VD2N0
    param(:,32)=1.0;       %AlphaZS     assimilation efficiency - default = 0.7
    param(:,33)=0.2;       %BetaZS      GGE default = 0.3
    param(:,39)=10;      %PS2ZLstar   default = 0.04 (threshold)
    param(:,74)=4;       %setVP   sinking speed, default = 40
    param(:,75)=4;       %setVO   sinking speed, default = 40
elseif Trial==98
    SurfIrr=100;
    Temperature=5;
    Kzmixed=10^-3;
    Kzdeep = 10^-4;
    param(:,9)=0.03;       %ResPS0 default is 0.03
    param(:,21)=0.03;      %ResPL0 default is 0.03
    param(:,18)=15;       %PsiL = ammonium inhibition coeff (smaller is less inhibition) default is 4.6
    param(:,62)=1;       %VP2N0
    param(:,64)=1;      %VP2D0
    param(:,66)=1;      %VD2N0
    param(:,32)=1.0;       %AlphaZS     assimilation efficiency - default = 0.7
    param(:,33)=0.2;       %BetaZS      GGE default = 0.3
    param(:,39)=10;      %PS2ZLstar   default = 0.04 (threshold)
    param(:,74)=4;       %setVP   sinking speed, default = 40
    param(:,75)=4;       %setVO   sinking speed, default = 40
elseif Trial==99
    Kzmixed=10^-3;
    Kzdeep = 10^-4;
    param(:,9)=0.03;       %ResPS0 default is 0.03
    param(:,21)=0.03;      %ResPL0 default is 0.03
    param(:,11)=0.4;     %gamS   small phyto active excretion rate (default = 0.135)
    param(:,23)=0.4;     %gamL   large phyto active excretion rate (default = 0.135)
    param(:,18)=15;       %PsiL = ammonium inhibition coeff (smaller is less inhibition) default is 4.6
    param(:,62)=1;       %VP2N0
    param(:,64)=1;      %VP2D0
    param(:,66)=1;      %VD2N0
    param(:,32)=1.0;       %AlphaZS     assimilation efficiency - default = 0.7
    param(:,33)=0.2;       %BetaZS      GGE default = 0.3
    param(:,39)=10;      %PS2ZLstar   default = 0.04 (threshold)
    param(:,74)=4;       %setVP   sinking speed, default = 40
    param(:,75)=4;       %setVO   sinking speed, default = 40
elseif Trial==100
    Kzmixed=10^-3;
    Kzdeep = 10^-5;
    param(:,9)=0.03;       %ResPS0 default is 0.03
    param(:,21)=0.03;      %ResPL0 default is 0.03
    param(:,11)=0.4;     %gamS   small phyto active excretion rate (default = 0.135)
    param(:,23)=0.4;     %gamL   large phyto active excretion rate (default = 0.135)
    param(:,18)=15;       %PsiL = ammonium inhibition coeff (smaller is less inhibition) default is 4.6
    param(:,62)=1;       %VP2N0
    param(:,64)=1;      %VP2D0
    param(:,66)=1;      %VD2N0
    param(:,32)=1.0;       %AlphaZS     assimilation efficiency - default = 0.7
    param(:,33)=0.2;       %BetaZS      GGE default = 0.3
    param(:,39)=10;      %PS2ZLstar   default = 0.04 (threshold)
    param(:,74)=4;       %setVP   sinking speed, default = 40
    param(:,75)=4;       %setVO   sinking speed, default = 40
elseif Trial==101
    Kzmixed=10^-4;
    Kzdeep = 10^-5;
    param(:,9)=0.03;       %ResPS0 default is 0.03
    param(:,21)=0.03;      %ResPL0 default is 0.03
    param(:,11)=0.4;     %gamS   small phyto active excretion rate (default = 0.135)
    param(:,23)=0.4;     %gamL   large phyto active excretion rate (default = 0.135)
    param(:,18)=15;       %PsiL = ammonium inhibition coeff (smaller is less inhibition) default is 4.6
    param(:,62)=1;       %VP2N0
    param(:,64)=1;      %VP2D0
    param(:,66)=1;      %VD2N0
    param(:,32)=1.0;       %AlphaZS     assimilation efficiency - default = 0.7
    param(:,33)=0.2;       %BetaZS      GGE default = 0.3
    param(:,39)=10;      %PS2ZLstar   default = 0.04 (threshold)
    param(:,74)=4;       %setVP   sinking speed, default = 40
    param(:,75)=4;       %setVO   sinking speed, default = 40
elseif Trial==102
    Kzmixed=10^-3;
    Kzdeep = 10^-5;
    param(:,9)=0.0;       %ResPS0 default is 0.03
    param(:,21)=0.0;      %ResPL0 default is 0.03
    param(:,11)=0.4;     %gamS   small phyto active excretion rate (default = 0.135)
    param(:,23)=0.4;     %gamL   large phyto active excretion rate (default = 0.135)
    param(:,18)=15;       %PsiL = ammonium inhibition coeff (smaller is less inhibition) default is 4.6
    param(:,62)=1;       %VP2N0
    param(:,64)=1;      %VP2D0
    param(:,66)=1;      %VD2N0
    param(:,32)=1.0;       %AlphaZS     assimilation efficiency - default = 0.7
    param(:,33)=0.2;       %BetaZS      GGE default = 0.3
    param(:,39)=10;      %PS2ZLstar   default = 0.04 (threshold)
    param(:,74)=4;       %setVP   sinking speed, default = 40
    param(:,75)=4;       %setVO   sinking speed, default = 40
elseif Trial==103
    Kzmixed=10^-4;
    Kzdeep = 10^-5;
    param(:,9)=0.0;       %ResPS0 default is 0.03
    param(:,21)=0.0;      %ResPL0 default is 0.03
    param(:,11)=0.4;     %gamS   small phyto active excretion rate (default = 0.135)
    param(:,23)=0.4;     %gamL   large phyto active excretion rate (default = 0.135)
    param(:,18)=15;       %PsiL = ammonium inhibition coeff (smaller is less inhibition) default is 4.6
    param(:,62)=1;       %VP2N0
    param(:,64)=1;      %VP2D0
    param(:,66)=1;      %VD2N0
    param(:,32)=1.0;       %AlphaZS     assimilation efficiency - default = 0.7
    param(:,33)=0.2;       %BetaZS      GGE default = 0.3
    param(:,39)=10;      %PS2ZLstar   default = 0.04 (threshold)
    param(:,74)=4;       %setVP   sinking speed, default = 40
    param(:,75)=4;       %setVO   sinking speed, default = 40
elseif Trial==104
    Kzmixed=10^-3;
    Kzdeep = 10^-5;
    param(:,9)=0.0;       %ResPS0 default is 0.03
    param(:,21)=0.0;      %ResPL0 default is 0.03
    param(:,11)=0.03;     %gamS   small phyto active excretion rate (default = 0.135)
    param(:,23)=0.03;     %gamL   large phyto active excretion rate (default = 0.135)
    param(:,18)=15;       %PsiL = ammonium inhibition coeff (smaller is less inhibition) default is 4.6
    param(:,62)=1;       %VP2N0
    param(:,64)=1;      %VP2D0
    param(:,66)=1;      %VD2N0
    param(:,32)=1.0;       %AlphaZS     assimilation efficiency - default = 0.7
    param(:,33)=0.2;       %BetaZS      GGE default = 0.3
    param(:,39)=10;      %PS2ZLstar   default = 0.04 (threshold)
    param(:,74)=4;       %setVP   sinking speed, default = 40
    param(:,75)=4;       %setVO   sinking speed, default = 40
elseif Trial==105
    Kzmixed=10^-4;
    Kzdeep = 10^-5;
    param(:,9)=0.0;       %ResPS0 default is 0.03
    param(:,21)=0.0;      %ResPL0 default is 0.03
    param(:,11)=0.03;     %gamS   small phyto active excretion rate (default = 0.135)
    param(:,23)=0.03;     %gamL   large phyto active excretion rate (default = 0.135)
    param(:,18)=15;       %PsiL = ammonium inhibition coeff (smaller is less inhibition) default is 4.6
    param(:,62)=1;       %VP2N0
    param(:,64)=1;      %VP2D0
    param(:,66)=1;      %VD2N0
    param(:,32)=1.0;       %AlphaZS     assimilation efficiency - default = 0.7
    param(:,33)=0.2;       %BetaZS      GGE default = 0.3
    param(:,39)=10;      %PS2ZLstar   default = 0.04 (threshold)
    param(:,74)=4;       %setVP   sinking speed, default = 40
    param(:,75)=4;       %setVO   sinking speed, default = 40
elseif Trial==106
    Kzmixed=10^-3;
    Kzdeep = 10^-4;
    param(:,9)=0.03;       %ResPS0 default is 0.03
    param(:,21)=0.03;      %ResPL0 default is 0.03
    param(:,11)=0.4;     %gamS   small phyto active excretion rate (default = 0.135)
    param(:,23)=0.4;     %gamL   large phyto active excretion rate (default = 0.135)
    param(:,18)=15;       %PsiL = ammonium inhibition coeff (smaller is less inhibition) default is 4.6
    param(:,62)=0.1;       %VP2N0
    param(:,64)=0.1;      %VP2D0
    param(:,66)=0.1;      %VD2N0
    param(:,32)=1.0;       %AlphaZS     assimilation efficiency - default = 0.7
    param(:,33)=0.2;       %BetaZS      GGE default = 0.3
    param(:,39)=10;      %PS2ZLstar   default = 0.04 (threshold)
    param(:,74)=4;       %setVP   sinking speed, default = 40
    param(:,75)=4;       %setVO   sinking speed, default = 40
elseif Trial==107
    Kzmixed=10^-3;
    Kzdeep = 10^-5;
    param(:,9)=0.03;       %ResPS0 default is 0.03
    param(:,21)=0.03;      %ResPL0 default is 0.03
    param(:,11)=0.4;     %gamS   small phyto active excretion rate (default = 0.135)
    param(:,23)=0.4;     %gamL   large phyto active excretion rate (default = 0.135)
    param(:,18)=15;       %PsiL = ammonium inhibition coeff (smaller is less inhibition) default is 4.6
    param(:,62)=0.1;       %VP2N0
    param(:,64)=0.1;      %VP2D0
    param(:,66)=0.1;      %VD2N0
    param(:,32)=1.0;       %AlphaZS     assimilation efficiency - default = 0.7
    param(:,33)=0.2;       %BetaZS      GGE default = 0.3
    param(:,39)=10;      %PS2ZLstar   default = 0.04 (threshold)
    param(:,74)=4;       %setVP   sinking speed, default = 40
    param(:,75)=4;       %setVO   sinking speed, default = 40
elseif Trial==108
    Kzmixed=10^-4;
    Kzdeep = 10^-5;
    param(:,9)=0.03;       %ResPS0 default is 0.03
    param(:,21)=0.03;      %ResPL0 default is 0.03
    param(:,11)=0.4;     %gamS   small phyto active excretion rate (default = 0.135)
    param(:,23)=0.4;     %gamL   large phyto active excretion rate (default = 0.135)
    param(:,18)=15;       %PsiL = ammonium inhibition coeff (smaller is less inhibition) default is 4.6
    param(:,62)=0.1;       %VP2N0
    param(:,64)=0.1;      %VP2D0
    param(:,66)=0.1;      %VD2N0
    param(:,32)=1.0;       %AlphaZS     assimilation efficiency - default = 0.7
    param(:,33)=0.2;       %BetaZS      GGE default = 0.3
    param(:,39)=10;      %PS2ZLstar   default = 0.04 (threshold)
    param(:,74)=4;       %setVP   sinking speed, default = 40
    param(:,75)=4;       %setVO   sinking speed, default = 40
elseif Trial==109
    Kzmixed=10^-3;
    Kzdeep = 10^-5;
    param(:,9)=0.0;       %ResPS0 default is 0.03
    param(:,21)=0.0;      %ResPL0 default is 0.03
    param(:,11)=0.4;     %gamS   small phyto active excretion rate (default = 0.135)
    param(:,23)=0.4;     %gamL   large phyto active excretion rate (default = 0.135)
    param(:,18)=15;       %PsiL = ammonium inhibition coeff (smaller is less inhibition) default is 4.6
    param(:,62)=0.1;       %VP2N0
    param(:,64)=0.1;      %VP2D0
    param(:,66)=0.1;      %VD2N0
    param(:,32)=1.0;       %AlphaZS     assimilation efficiency - default = 0.7
    param(:,33)=0.2;       %BetaZS      GGE default = 0.3
    param(:,39)=10;      %PS2ZLstar   default = 0.04 (threshold)
    param(:,74)=4;       %setVP   sinking speed, default = 40
    param(:,75)=4;       %setVO   sinking speed, default = 40
elseif Trial==110
    Kzmixed=10^-4;
    Kzdeep = 10^-5;
    param(:,9)=0.0;       %ResPS0 default is 0.03
    param(:,21)=0.0;      %ResPL0 default is 0.03
    param(:,11)=0.4;     %gamS   small phyto active excretion rate (default = 0.135)
    param(:,23)=0.4;     %gamL   large phyto active excretion rate (default = 0.135)
    param(:,18)=15;       %PsiL = ammonium inhibition coeff (smaller is less inhibition) default is 4.6
    param(:,62)=0.1;       %VP2N0
    param(:,64)=0.1;      %VP2D0
    param(:,66)=0.1;      %VD2N0
    param(:,32)=1.0;       %AlphaZS     assimilation efficiency - default = 0.7
    param(:,33)=0.2;       %BetaZS      GGE default = 0.3
    param(:,39)=10;      %PS2ZLstar   default = 0.04 (threshold)
    param(:,74)=4;       %setVP   sinking speed, default = 40
    param(:,75)=4;       %setVO   sinking speed, default = 40
elseif Trial==111
    Kzmixed=10^-3;
    Kzdeep = 10^-5;
    param(:,9)=0.0;       %ResPS0 default is 0.03
    param(:,21)=0.0;      %ResPL0 default is 0.03
    param(:,11)=0.03;     %gamS   small phyto active excretion rate (default = 0.135)
    param(:,23)=0.03;     %gamL   large phyto active excretion rate (default = 0.135)
    param(:,18)=15;       %PsiL = ammonium inhibition coeff (smaller is less inhibition) default is 4.6
    param(:,62)=0.1;       %VP2N0
    param(:,64)=0.1;      %VP2D0
    param(:,66)=0.1;      %VD2N0
    param(:,32)=1.0;       %AlphaZS     assimilation efficiency - default = 0.7
    param(:,33)=0.2;       %BetaZS      GGE default = 0.3
    param(:,39)=10;      %PS2ZLstar   default = 0.04 (threshold)
    param(:,74)=4;       %setVP   sinking speed, default = 40
    param(:,75)=4;       %setVO   sinking speed, default = 40
elseif Trial==112
    Kzmixed=10^-4;
    Kzdeep = 10^-5;
    param(:,9)=0.0;       %ResPS0 default is 0.03
    param(:,21)=0.0;      %ResPL0 default is 0.03
    param(:,11)=0.03;     %gamS   small phyto active excretion rate (default = 0.135)
    param(:,23)=0.03;     %gamL   large phyto active excretion rate (default = 0.135)
    param(:,18)=15;       %PsiL = ammonium inhibition coeff (smaller is less inhibition) default is 4.6
    param(:,62)=0.1;       %VP2N0
    param(:,64)=0.1;      %VP2D0
    param(:,66)=0.1;      %VD2N0
    param(:,32)=1.0;       %AlphaZS     assimilation efficiency - default = 0.7
    param(:,33)=0.2;       %BetaZS      GGE default = 0.3
    param(:,39)=10;      %PS2ZLstar   default = 0.04 (threshold)
    param(:,74)=4;       %setVP   sinking speed, default = 40
    param(:,75)=4;       %setVO   sinking speed, default = 40
elseif Trial==113
    Kzmixed=10^-3;
    Kzdeep = 10^-4;
    param(:,9)=0.03;       %ResPS0 default is 0.03
    param(:,21)=0.03;      %ResPL0 default is 0.03
    param(:,11)=0.4;     %gamS   small phyto active excretion rate (default = 0.135)
    param(:,23)=0.4;     %gamL   large phyto active excretion rate (default = 0.135)
    param(:,18)=1.5;       %PsiL = ammonium inhibition coeff (smaller is less inhibition) default is 4.6
    param(:,62)=1;       %VP2N0
    param(:,64)=1;      %VP2D0
    param(:,66)=1;      %VD2N0
    param(:,32)=1.0;       %AlphaZS     assimilation efficiency - default = 0.7
    param(:,33)=0.2;       %BetaZS      GGE default = 0.3
    param(:,39)=10;      %PS2ZLstar   default = 0.04 (threshold)
    param(:,74)=4;       %setVP   sinking speed, default = 40
    param(:,75)=4;       %setVO   sinking speed, default = 40
elseif Trial==114
    Kzmixed=10^-3;
    Kzdeep = 10^-5;
    param(:,9)=0.03;       %ResPS0 default is 0.03
    param(:,21)=0.03;      %ResPL0 default is 0.03
    param(:,11)=0.4;     %gamS   small phyto active excretion rate (default = 0.135)
    param(:,23)=0.4;     %gamL   large phyto active excretion rate (default = 0.135)
    param(:,18)=1.5;       %PsiL = ammonium inhibition coeff (smaller is less inhibition) default is 4.6
    param(:,62)=1;       %VP2N0
    param(:,64)=1;      %VP2D0
    param(:,66)=1;      %VD2N0
    param(:,32)=1.0;       %AlphaZS     assimilation efficiency - default = 0.7
    param(:,33)=0.2;       %BetaZS      GGE default = 0.3
    param(:,39)=10;      %PS2ZLstar   default = 0.04 (threshold)
    param(:,74)=4;       %setVP   sinking speed, default = 40
    param(:,75)=4;       %setVO   sinking speed, default = 40
elseif Trial==145
    Kzmixed=10^-4;
    Kzdeep = 10^-5;
    param(:,9)=0.03;       %ResPS0 default is 0.03
    param(:,21)=0.03;      %ResPL0 default is 0.03
    param(:,11)=0.4;     %gamS   small phyto active excretion rate (default = 0.135)
    param(:,23)=0.4;     %gamL   large phyto active excretion rate (default = 0.135)
    param(:,18)=1.5;       %PsiL = ammonium inhibition coeff (smaller is less inhibition) default is 4.6
    param(:,62)=1;       %VP2N0
    param(:,64)=1;      %VP2D0
    param(:,66)=1;      %VD2N0
    param(:,32)=1.0;       %AlphaZS     assimilation efficiency - default = 0.7
    param(:,33)=0.2;       %BetaZS      GGE default = 0.3
    param(:,39)=10;      %PS2ZLstar   default = 0.04 (threshold)
    param(:,74)=4;       %setVP   sinking speed, default = 40
    param(:,75)=4;       %setVO   sinking speed, default = 40
elseif Trial==146
    Kzmixed=10^-3;
    Kzdeep = 10^-5;
    param(:,9)=0.0;       %ResPS0 default is 0.03
    param(:,21)=0.0;      %ResPL0 default is 0.03
    param(:,11)=0.4;     %gamS   small phyto active excretion rate (default = 0.135)
    param(:,23)=0.4;     %gamL   large phyto active excretion rate (default = 0.135)
    param(:,18)=1.5;       %PsiL = ammonium inhibition coeff (smaller is less inhibition) default is 4.6
    param(:,62)=1;       %VP2N0
    param(:,64)=1;      %VP2D0
    param(:,66)=1;      %VD2N0
    param(:,32)=1.0;       %AlphaZS     assimilation efficiency - default = 0.7
    param(:,33)=0.2;       %BetaZS      GGE default = 0.3
    param(:,39)=10;      %PS2ZLstar   default = 0.04 (threshold)
    param(:,74)=4;       %setVP   sinking speed, default = 40
    param(:,75)=4;       %setVO   sinking speed, default = 40
elseif Trial==117
    Kzmixed=10^-4;
    Kzdeep = 10^-5;
    param(:,9)=0.0;       %ResPS0 default is 0.03
    param(:,21)=0.0;      %ResPL0 default is 0.03
    param(:,11)=0.4;     %gamS   small phyto active excretion rate (default = 0.135)
    param(:,23)=0.4;     %gamL   large phyto active excretion rate (default = 0.135)
    param(:,18)=1.5;       %PsiL = ammonium inhibition coeff (smaller is less inhibition) default is 4.6
    param(:,62)=1;       %VP2N0
    param(:,64)=1;      %VP2D0
    param(:,66)=1;      %VD2N0
    param(:,32)=1.0;       %AlphaZS     assimilation efficiency - default = 0.7
    param(:,33)=0.2;       %BetaZS      GGE default = 0.3
    param(:,39)=10;      %PS2ZLstar   default = 0.04 (threshold)
    param(:,74)=4;       %setVP   sinking speed, default = 40
    param(:,75)=4;       %setVO   sinking speed, default = 40
elseif Trial==118
    Kzmixed=10^-3;
    Kzdeep = 10^-5;
    param(:,9)=0.0;       %ResPS0 default is 0.03
    param(:,21)=0.0;      %ResPL0 default is 0.03
    param(:,11)=0.03;     %gamS   small phyto active excretion rate (default = 0.135)
    param(:,23)=0.03;     %gamL   large phyto active excretion rate (default = 0.135)
    param(:,18)=1.5;       %PsiL = ammonium inhibition coeff (smaller is less inhibition) default is 4.6
    param(:,62)=1;       %VP2N0
    param(:,64)=1;      %VP2D0
    param(:,66)=1;      %VD2N0
    param(:,32)=1.0;       %AlphaZS     assimilation efficiency - default = 0.7
    param(:,33)=0.2;       %BetaZS      GGE default = 0.3
    param(:,39)=10;      %PS2ZLstar   default = 0.04 (threshold)
    param(:,74)=4;       %setVP   sinking speed, default = 40
    param(:,75)=4;       %setVO   sinking speed, default = 40
elseif Trial==119
    Kzmixed=10^-4;
    Kzdeep = 10^-5;
    param(:,9)=0.0;       %ResPS0 default is 0.03
    param(:,21)=0.0;      %ResPL0 default is 0.03
    param(:,11)=0.03;     %gamS   small phyto active excretion rate (default = 0.135)
    param(:,23)=0.03;     %gamL   large phyto active excretion rate (default = 0.135)
    param(:,18)=1.5;       %PsiL = ammonium inhibition coeff (smaller is less inhibition) default is 4.6
    param(:,62)=1;       %VP2N0
    param(:,64)=1;      %VP2D0
    param(:,66)=1;      %VD2N0
    param(:,32)=1.0;       %AlphaZS     assimilation efficiency - default = 0.7
    param(:,33)=0.2;       %BetaZS      GGE default = 0.3
    param(:,39)=10;      %PS2ZLstar   default = 0.04 (threshold)
    param(:,74)=4;       %setVP   sinking speed, default = 40
    param(:,75)=4;       %setVO   sinking speed, default = 40
elseif Trial==120
    Kzmixed=10^-3;
    Kzdeep = 10^-4;
    param(:,9)=0.03;       %ResPS0 default is 0.03
    param(:,21)=0.03;      %ResPL0 default is 0.03
    param(:,11)=0.4;     %gamS   small phyto active excretion rate (default = 0.135)
    param(:,23)=0.4;     %gamL   large phyto active excretion rate (default = 0.135)
    param(:,18)=1.5;       %PsiL = ammonium inhibition coeff (smaller is less inhibition) default is 4.6
    param(:,62)=0.1;       %VP2N0
    param(:,64)=0.1;      %VP2D0
    param(:,66)=0.1;      %VD2N0
    param(:,32)=1.0;       %AlphaZS     assimilation efficiency - default = 0.7
    param(:,33)=0.2;       %BetaZS      GGE default = 0.3
    param(:,39)=10;      %PS2ZLstar   default = 0.04 (threshold)
    param(:,74)=4;       %setVP   sinking speed, default = 40
    param(:,75)=4;       %setVO   sinking speed, default = 40
elseif Trial==121
    Kzmixed=10^-3;
    Kzdeep = 10^-5;
    param(:,9)=0.03;       %ResPS0 default is 0.03
    param(:,21)=0.03;      %ResPL0 default is 0.03
    param(:,11)=0.4;     %gamS   small phyto active excretion rate (default = 0.135)
    param(:,23)=0.4;     %gamL   large phyto active excretion rate (default = 0.135)
    param(:,18)=1.5;       %PsiL = ammonium inhibition coeff (smaller is less inhibition) default is 4.6
    param(:,62)=0.1;       %VP2N0
    param(:,64)=0.1;      %VP2D0
    param(:,66)=0.1;      %VD2N0
    param(:,32)=1.0;       %AlphaZS     assimilation efficiency - default = 0.7
    param(:,33)=0.2;       %BetaZS      GGE default = 0.3
    param(:,39)=10;      %PS2ZLstar   default = 0.04 (threshold)
    param(:,74)=4;       %setVP   sinking speed, default = 40
    param(:,75)=4;       %setVO   sinking speed, default = 40
elseif Trial==122
    Kzmixed=10^-4;
    Kzdeep = 10^-5;
    param(:,9)=0.03;       %ResPS0 default is 0.03
    param(:,21)=0.03;      %ResPL0 default is 0.03
    param(:,11)=0.4;     %gamS   small phyto active excretion rate (default = 0.135)
    param(:,23)=0.4;     %gamL   large phyto active excretion rate (default = 0.135)
    param(:,18)=1.5;       %PsiL = ammonium inhibition coeff (smaller is less inhibition) default is 4.6
    param(:,62)=0.1;       %VP2N0
    param(:,64)=0.1;      %VP2D0
    param(:,66)=0.1;      %VD2N0
    param(:,32)=1.0;       %AlphaZS     assimilation efficiency - default = 0.7
    param(:,33)=0.2;       %BetaZS      GGE default = 0.3
    param(:,39)=10;      %PS2ZLstar   default = 0.04 (threshold)
    param(:,74)=4;       %setVP   sinking speed, default = 40
    param(:,75)=4;       %setVO   sinking speed, default = 40
elseif Trial==123
    Kzmixed=10^-3;
    Kzdeep = 10^-5;
    param(:,9)=0.0;       %ResPS0 default is 0.03
    param(:,21)=0.0;      %ResPL0 default is 0.03
    param(:,11)=0.4;     %gamS   small phyto active excretion rate (default = 0.135)
    param(:,23)=0.4;     %gamL   large phyto active excretion rate (default = 0.135)
    param(:,18)=1.5;       %PsiL = ammonium inhibition coeff (smaller is less inhibition) default is 4.6
    param(:,62)=0.1;       %VP2N0
    param(:,64)=0.1;      %VP2D0
    param(:,66)=0.1;      %VD2N0
    param(:,32)=1.0;       %AlphaZS     assimilation efficiency - default = 0.7
    param(:,33)=0.2;       %BetaZS      GGE default = 0.3
    param(:,39)=10;      %PS2ZLstar   default = 0.04 (threshold)
    param(:,74)=4;       %setVP   sinking speed, default = 40
    param(:,75)=4;       %setVO   sinking speed, default = 40
elseif Trial==124
    Kzmixed=10^-4;
    Kzdeep = 10^-5;
    param(:,9)=0.0;       %ResPS0 default is 0.03
    param(:,21)=0.0;      %ResPL0 default is 0.03
    param(:,11)=0.4;     %gamS   small phyto active excretion rate (default = 0.135)
    param(:,23)=0.4;     %gamL   large phyto active excretion rate (default = 0.135)
    param(:,18)=1.5;       %PsiL = ammonium inhibition coeff (smaller is less inhibition) default is 4.6
    param(:,62)=0.1;       %VP2N0
    param(:,64)=0.1;      %VP2D0
    param(:,66)=0.1;      %VD2N0
    param(:,32)=1.0;       %AlphaZS     assimilation efficiency - default = 0.7
    param(:,33)=0.2;       %BetaZS      GGE default = 0.3
    param(:,39)=10;      %PS2ZLstar   default = 0.04 (threshold)
    param(:,74)=4;       %setVP   sinking speed, default = 40
    param(:,75)=4;       %setVO   sinking speed, default = 40
elseif Trial==125
    Kzmixed=10^-3;
    Kzdeep = 10^-5;
    param(:,9)=0.0;       %ResPS0 default is 0.03
    param(:,21)=0.0;      %ResPL0 default is 0.03
    param(:,11)=0.03;     %gamS   small phyto active excretion rate (default = 0.135)
    param(:,23)=0.03;     %gamL   large phyto active excretion rate (default = 0.135)
    param(:,18)=1.5;       %PsiL = ammonium inhibition coeff (smaller is less inhibition) default is 4.6
    param(:,62)=0.1;       %VP2N0
    param(:,64)=0.1;      %VP2D0
    param(:,66)=0.1;      %VD2N0
    param(:,32)=1.0;       %AlphaZS     assimilation efficiency - default = 0.7
    param(:,33)=0.2;       %BetaZS      GGE default = 0.3
    param(:,39)=10;      %PS2ZLstar   default = 0.04 (threshold)
    param(:,74)=4;       %setVP   sinking speed, default = 40
    param(:,75)=4;       %setVO   sinking speed, default = 40
elseif Trial==126
    Kzmixed=10^-4;
    Kzdeep = 10^-5;
    param(:,9)=0.0;       %ResPS0 default is 0.03
    param(:,21)=0.0;      %ResPL0 default is 0.03
    param(:,11)=0.03;     %gamS   small phyto active excretion rate (default = 0.135)
    param(:,23)=0.03;     %gamL   large phyto active excretion rate (default = 0.135)
    param(:,18)=1.5;       %PsiL = ammonium inhibition coeff (smaller is less inhibition) default is 4.6
    param(:,62)=0.1;       %VP2N0
    param(:,64)=0.1;      %VP2D0
    param(:,66)=0.1;      %VD2N0
    param(:,32)=1.0;       %AlphaZS     assimilation efficiency - default = 0.7
    param(:,33)=0.2;       %BetaZS      GGE default = 0.3
    param(:,39)=10;      %PS2ZLstar   default = 0.04 (threshold)
    param(:,74)=4;       %setVP   sinking speed, default = 40
    param(:,75)=4;       %setVO   sinking speed, default = 40
elseif Trial==127
    Kzmixed=10^-4;
    Kzdeep = 10^-5;
    param(:,9)=0.0;       %ResPS0 default is 0.03
    param(:,21)=0.0;      %ResPL0 default is 0.03
    param(:,11)=0.03;     %gamS   small phyto active excretion rate (default = 0.135)
    param(:,23)=0.03;     %gamL   large phyto active excretion rate (default = 0.135)
    param(:,18)=1.5;       %PsiL = ammonium inhibition coeff (smaller is less inhibition) default is 4.6
    param(:,62)=0.1;       %VP2N0
    param(:,64)=0.1;      %VP2D0
    param(:,66)=0.1;      %VD2N0
    param(:,32)=1.0;       %AlphaZS     assimilation efficiency - default = 0.7
    param(:,33)=0.2;       %BetaZS      GGE default = 0.3
    param(:,39)=10;      %PS2ZLstar   default = 0.04 (threshold)
    param(:,74)=4;       %setVP   sinking speed, default = 40
    param(:,75)=4;       %setVO   sinking speed, default = 40
    param(:,28)=0.2;       %lamS   default = 1.4 (lower decreases grazing at low biomass)
elseif Trial==128
    Kzmixed=10^-4;
    Kzdeep = 10^-5;
    param(:,9)=0.0;       %ResPS0 default is 0.03
    param(:,21)=0.0;      %ResPL0 default is 0.03
    param(:,11)=0.03;     %gamS   small phyto active excretion rate (default = 0.135)
    param(:,23)=0.03;     %gamL   large phyto active excretion rate (default = 0.135)
    param(:,18)=15;       %PsiL = ammonium inhibition coeff (smaller is less inhibition) default is 4.6
    param(:,62)=0.1;       %VP2N0
    param(:,64)=0.1;      %VP2D0
    param(:,66)=0.1;      %VD2N0
    param(:,32)=1.0;       %AlphaZS     assimilation efficiency - default = 0.7
    param(:,33)=0.2;       %BetaZS      GGE default = 0.3
    param(:,39)=10;      %PS2ZLstar   default = 0.04 (threshold)
    param(:,74)=4;       %setVP   sinking speed, default = 40
    param(:,75)=4;       %setVO   sinking speed, default = 40
    param(:,28)=0.2;       %lamS   default = 1.4 (lower decreases grazing at low biomass)
elseif Trial==128
    Kzmixed=10^-4;
    Kzdeep = 10^-5;
    param(:,9)=0.0;       %ResPS0 default is 0.03
    param(:,21)=0.0;      %ResPL0 default is 0.03
    param(:,11)=0.03;     %gamS   small phyto active excretion rate (default = 0.135)
    param(:,23)=0.03;     %gamL   large phyto active excretion rate (default = 0.135)
    param(:,18)=1.5;       %PsiL = ammonium inhibition coeff (smaller is less inhibition) default is 4.6
    param(:,62)=1;       %VP2N0
    param(:,64)=1;      %VP2D0
    param(:,66)=1;      %VD2N0
    param(:,32)=1.0;       %AlphaZS     assimilation efficiency - default = 0.7
    param(:,33)=0.2;       %BetaZS      GGE default = 0.3
    param(:,39)=10;      %PS2ZLstar   default = 0.04 (threshold)
    param(:,74)=4;       %setVP   sinking speed, default = 40
    param(:,75)=4;       %setVO   sinking speed, default = 40
    param(:,28)=0.2;       %lamS   default = 1.4 (lower decreases grazing at low biomass)
elseif Trial==129
    Kzmixed=10^-4;
    Kzdeep = 10^-5;
    param(:,9)=0.0;       %ResPS0 default is 0.03
    param(:,21)=0.0;      %ResPL0 default is 0.03
    param(:,11)=0.135;     %gamS   small phyto active excretion rate (default = 0.135)
    param(:,23)=0.135;     %gamL   large phyto active excretion rate (default = 0.135)
    param(:,18)=1.5;       %PsiL = ammonium inhibition coeff (smaller is less inhibition) default is 4.6
    param(:,62)=1;       %VP2N0
    param(:,64)=1;      %VP2D0
    param(:,66)=1;      %VD2N0
    param(:,32)=1.0;       %AlphaZS     assimilation efficiency - default = 0.7
    param(:,33)=0.2;       %BetaZS      GGE default = 0.3
    param(:,39)=10;      %PS2ZLstar   default = 0.04 (threshold)
    param(:,74)=4;       %setVP   sinking speed, default = 40
    param(:,75)=4;       %setVO   sinking speed, default = 40
    param(:,28)=0.2;       %lamS   default = 1.4 (lower decreases grazing at low biomass)
elseif Trial==130
    Kzmixed=10^-4;
    Kzdeep = 10^-5;
    param(:,9)=0.03;       %ResPS0 default is 0.03
    param(:,21)=0.03;      %ResPL0 default is 0.03
    param(:,11)=0.135;     %gamS   small phyto active excretion rate (default = 0.135)
    param(:,23)=0.135;     %gamL   large phyto active excretion rate (default = 0.135)
    param(:,18)=1.5;       %PsiL = ammonium inhibition coeff (smaller is less inhibition) default is 4.6
    param(:,62)=1;       %VP2N0
    param(:,64)=1;      %VP2D0
    param(:,66)=1;      %VD2N0
    param(:,32)=1.0;       %AlphaZS     assimilation efficiency - default = 0.7
    param(:,33)=0.2;       %BetaZS      GGE default = 0.3
    param(:,39)=10;      %PS2ZLstar   default = 0.04 (threshold)
    param(:,74)=4;       %setVP   sinking speed, default = 40
    param(:,75)=4;       %setVO   sinking speed, default = 40
    param(:,28)=0.2;       %lamS   default = 1.4 (lower decreases grazing at low biomass)
elseif Trial==131
    Kzmixed=10^-4;
    Kzdeep = 10^-5;
    param(:,9)=0.03;       %ResPS0 default is 0.03
    param(:,21)=0.03;      %ResPL0 default is 0.03
    param(:,11)=0.135;     %gamS   small phyto active excretion rate (default = 0.135)
    param(:,23)=0.135;     %gamL   large phyto active excretion rate (default = 0.135)
    param(:,18)=1.5;       %PsiL = ammonium inhibition coeff (smaller is less inhibition) default is 4.6
    param(:,62)=0.1;       %VP2N0
    param(:,64)=0.1;      %VP2D0
    param(:,66)=0.1;      %VD2N0
    param(:,32)=1.0;       %AlphaZS     assimilation efficiency - default = 0.7
    param(:,33)=0.2;       %BetaZS      GGE default = 0.3
    param(:,39)=10;      %PS2ZLstar   default = 0.04 (threshold)
    param(:,74)=4;       %setVP   sinking speed, default = 40
    param(:,75)=4;       %setVO   sinking speed, default = 40
    param(:,28)=0.2;       %lamS   default = 1.4 (lower decreases grazing at low biomass)
elseif Trial==132
    Kzmixed=10^-4;
    Kzdeep = 10^-5;
    param(:,9)=0.03;       %ResPS0 default is 0.03
    param(:,21)=0.03;      %ResPL0 default is 0.03
    param(:,11)=0.135;     %gamS   small phyto active excretion rate (default = 0.135)
    param(:,23)=0.135;     %gamL   large phyto active excretion rate (default = 0.135)
    param(:,18)=1.5;       %PsiL = ammonium inhibition coeff (smaller is less inhibition) default is 4.6
    param(:,62)=0.1;       %VP2N0
    param(:,64)=0.1;      %VP2D0
    param(:,66)=0.1;      %VD2N0
    param(:,32)=1.0;       %AlphaZS     assimilation efficiency - default = 0.7
    param(:,33)=0.2;       %BetaZS      GGE default = 0.3
    param(:,39)=10;      %PS2ZLstar   default = 0.04 (threshold)
    param(:,74)=40;       %setVP   sinking speed, default = 40
    param(:,75)=40;       %setVO   sinking speed, default = 40
    param(:,28)=0.2;       %lamS   default = 1.4 (lower decreases grazing at low biomass)
elseif Trial==133
    Kzmixed=10^-3;
    Kzdeep = 10^-5;
    param(:,9)=0.03;       %ResPS0 default is 0.03
    param(:,21)=0.03;      %ResPL0 default is 0.03
    param(:,11)=0.135;     %gamS   small phyto active excretion rate (default = 0.135)
    param(:,23)=0.135;     %gamL   large phyto active excretion rate (default = 0.135)
    param(:,18)=1.5;       %PsiL = ammonium inhibition coeff (smaller is less inhibition) default is 4.6
    param(:,62)=0.1;       %VP2N0
    param(:,64)=0.1;      %VP2D0
    param(:,66)=0.1;      %VD2N0
    param(:,32)=1.0;       %AlphaZS     assimilation efficiency - default = 0.7
    param(:,33)=0.2;       %BetaZS      GGE default = 0.3
    param(:,39)=10;      %PS2ZLstar   default = 0.04 (threshold)
    param(:,74)=40;       %setVP   sinking speed, default = 40
    param(:,75)=40;       %setVO   sinking speed, default = 40
    param(:,28)=0.2;       %lamS   default = 1.4 (lower decreases grazing at low biomass)
elseif Trial==134
    Kzmixed=10^-3;
    Kzdeep = 10^-4;
    param(:,9)=0.03;       %ResPS0 default is 0.03
    param(:,21)=0.03;      %ResPL0 default is 0.03
    param(:,11)=0.135;     %gamS   small phyto active excretion rate (default = 0.135)
    param(:,23)=0.135;     %gamL   large phyto active excretion rate (default = 0.135)
    param(:,18)=1.5;       %PsiL = ammonium inhibition coeff (smaller is less inhibition) default is 4.6
    param(:,62)=0.1;       %VP2N0
    param(:,64)=0.1;      %VP2D0
    param(:,66)=0.1;      %VD2N0
    param(:,32)=1.0;       %AlphaZS     assimilation efficiency - default = 0.7
    param(:,33)=0.2;       %BetaZS      GGE default = 0.3
    param(:,39)=10;      %PS2ZLstar   default = 0.04 (threshold)
    param(:,74)=40;       %setVP   sinking speed, default = 40
    param(:,75)=40;       %setVO   sinking speed, default = 40
    param(:,28)=0.2;       %lamS   default = 1.4 (lower decreases grazing at low biomass)
end


param_out=param;
```
