## Supplementary material for "Investigating equations for measuring dissolved inorganic nutrient uptake in oligotrophic conditions": NEMURO+15N Files: Sinking.m

```
function[tracernew,tracer15Nnew]=Sinking(tracer,tracer15N,thickness,param,dt)

setVP=param(:,74);   
setVO=param(:,75);   

tracernew = tracer;
tracer15Nnew = tracer15N;

sinkN=tracer(:,8).*setVP.*dt./thickness;
sinkSi=tracer(:,11).*setVO.*dt./thickness;
sinkN15=tracer15N(:,8).*setVP.*dt./thickness;

tracernew(:,8)=tracer(:,8)-sinkN;
tracernew(2:end,8)=tracernew(2:end,8)+sinkN(1:end-1);

tracernew(:,11)=tracer(:,11)-sinkSi;
tracernew(2:end,11)=tracernew(2:end,11)+sinkSi(1:end-1);

tracer15Nnew(:,8)=tracer15N(:,8)-sinkN15;
tracer15Nnew(2:end,8)=tracer15Nnew(2:end,8)+sinkN15(1:end-1);
```
